# Efficient translation of chaperones by eIF5A safeguards proteostasis under heat stress

**DOI:** 10.64898/2026.10.27.762258

**Authors:** Vanessa Bernal, Lianqi Chi, Samoa Prieto-Diez, Joan Segui, Marina Barba-Aliaga, Paula Alepuz

**Affiliations:** Instituto de Biotecnología y Biomedicina (Biotecmed), Universitat de València, 46100 València, Spain; Departamento de Bioquímica y Biología Molecular, Universitat de València, 46100 València, Spain

**Keywords:** protein biosynthesis, Hsp70, Hsf1, protein aggregates, codon optimality

## Abstract

The increase in global temperature poses a major threat to cells and organisms because it disrupts cellular proteostasis. We discovered that the evolutionarily conserved and essential translation elongation factor eIF5A is necessary for a rapid response to heat-stress in yeast cells. Lack of eIF5A results in ribosome stalling on heat-stress-induced mRNAs, reducing the synthesis of chaperones, primarily impacting those that undergo robust synthesis via high codon optimality, such as Hsp70 Ssa1 and Hsp12 chaperones. This new eIF5A role in translation appears to be direct, as eIF5A binds preferentially to induced mRNAs with high codon optimality and its polysome association increases under heat-shock. Low levels of Hsp70 in eIF5A mutants delay Hsf1 feedback inhibition during stress adaptation and cause the accumulation of protein aggregates, resulting in low cellular viability to heat-stress. Overall, our results show that eIF5A is crucial for heat-stress resistance by facilitating rapid and efficient translation of high codon-optimal chaperones. Therefore, eIF5A emerges as a key regulator of cellular proteostasis, providing a novel perspective on its significant involvement in neurodegenerative diseases and aging.

## INTRODUCTION

Elevated temperatures are one of the most common stresses that cells and organisms can face. Heat stress endangers cellular proteostasis by causing numerous proteins to unfold and form aggregates, which must be resolved to avoid compromising cell viability. The heat-shock response (HSR) is evolutionarily conserved and involves the reprogramming of gene expression at the transcriptional and translational levels to produce, among other effects, a huge synthesis of Heat shock proteins (Hsps) (Craig & Schlesinger, 1985; Craig *et al*, 2003; Lindquist, 1986; Lindquist & Craig, 1988; Morano *et al*, 2012; Verghese *et al*, 2012). Hsps are primarily molecular chaperones that prevent protein unfolding and facilitate the refolding, aggregation, disaggregation or degradation of damaged proteins (Becker & Craig, 1994; Chen *et al*, 2011; Lindquist & Craig, 1988; Parsell & Lindquist, 1993; Verghese *et al*, 2012; Willmund *et al*, 2013).

In eukaryotes, such as the yeast *Saccharomyces cerevisiae*, the transcriptional reprogramming induced by the HSR is mostly conducted by the heat shock transcription factor 1 (Hsf1). The activation of Hsf1 occurs when its interaction with the chaperone Hsp70 is disrupted by the titration away of Hsp70 by stress-induced clients. The binding of Hsf1 to the promoters of HSP genes activate their transcription, which occurs in parallel to the repression of several hundreds of house-keeping genes and the inhibition of global cytoplasmic translation (Dea & Pincus, 2024; Verghese *et al*, 2012). Differently to the response to most stresses, the phosphorylation of the initiation factor eIF2alpha is not sufficient to generate the reduction of global translation observed under heat-stress (Ghadanian *et al*, 2025; Grouši *et al*, 2009; Spriggs *et al*, 2010). Although still not fully understood how global translation is repressed, the redistribution of ribosomes from house-keeping to HSP mRNAs is considered essential to achieve the huge synthesis of Hsp proteins needed upon heat shock (Crawford & Pavitt, 2018; Hernandez-Elvira & Sunnerhagen, 2022; Yamasaki & Anderson, 2008). Thus, mRNAs encoding house-keeping proteins are redirected from polysomes to storage sites such as P-bodies (PB) and/or stress granules (SG) (Corbet & Parker, 2019; Ivanov *et al*, 2019).

The preferential, abundant and rapid synthesis of Hsp proteins upon heat-stress requires, not only an efficient translation initiation but also a high speed of the ribosomes during elongation (Varenne *et al*, 1984, 1989; and reviewed in Ghadanian *et al*, 2025). As it has been shown in yeast and mammals, one of the factors strongly influencing translation elongation efficiency is the codon optimality of the mRNA coding sequences (CDS) (Brule & Grayhack, 2017; Hussmann *et al*, 2015; Letzring *et al*, 2010; Weinberg *et al*, 2016; and reviewed in Hanson & Coller, 2017; Stein & Frydman, 2018). Optimal codons are those quickly recognized by cognate tRNAs due to both their abundance and pairing efficiency (Presnyak *et al*, 2015). Interestingly, most Hsps are encoded by mRNAs with a high proportion of optimal codons, while only very few show a clear bias to non-optimal codons. Translation by ribosomes through non-optimal codon regions can produce ribosome stalling and ribosome collisions with the formation of two-ribosome or disome structures that signal for mRNA decay (Buschauer *et al*, 2020; Müller *et al*, 2023). Much less is known about the dynamics of the ribosomes in contexts of high density, where theoretical modeling suggests that disomes should stochastically form (Juszkiewicz *et al*, 2018; Park & Subramaniam, 2019). One such context is that of high-codon optimal mRNAs, which leads to heavy translation, but where ribosome elongation progresses efficiently enabling abundant synthesis of proteins (Arava *et al*, 2003). However, it is unclear how ribosomal collisions that lead to translation progress are differentiated from those that result in abrupt termination. It is also unknown whether specific factors are required to facilitate heavy translation.

Several recent works have linked the translation elongation factor eIF5A with different ribosome control pathways. eIF5A is a universally conserved, abundantly expressed and essential protein in eukaryotes (Barba-Aliaga & Alepuz, 2022; Dever *et al*, 2014; Mathews & Hershey, 2015; Tauc *et al*, 2021). The best-know role of eIF5A in ribosome elongation is to promote peptide bond formation when ribosomes stall in codons for consecutive prolines and combinations of proline, glycine and charged amino acids (Pelechano & Alepuz, 2017; Schuller *et al*, 2017). To do this, eIF5A must be activated by hypusination, a unique post-translation modification of this protein, which consists in the addition of the polyamine spermidine to a conserved lysine residue (Park *et al*, 2022). Hypusinated-eIF5A (Hyp-eIF5A) binds to stalled ribosomes through the empty E-site, stabilizing a productive conformation for peptide bond formation (Doerfel *et al*, 2015; Melnikov *et al*, 2016; Schmidt *et al*, 2016). Furthermore, eIF5A also stimulates translation termination of stalled ribosomes in specific STOP codons (Schuller *et al*, 2017).

However, the role of eIF5A in translation elongation seems to be wider, as eIF5A has been found associated to stalled ribosomes under other contexts. Thus, in disomes formed in yeast under amino acid starvation, the Gcn2 Kinase-activator factor Gcn1 interacts with both 80S ribosomes and eIF5A locates in the E-site of the stalled ribosome (Pochopien *et al*, 2021). eIF5A is also member of the ribosome-associated quality control (RQC) pathway, which recognizes ribosome collisions and disome structures within mRNA coding sequences (CDS) (Inada & Beckmann, 2024), binding to the E-site of the free 60S subunit to promote CAT tailing of truncated peptides and subsequent proteasomal degradation (Tesina *et al*, 2023). More recently, work in *Caenorhabditis elegans* and *S. cerevisiae* has proposed that eIF5A is used to identify ribosomal stalls and promote mRNA decay in the No-Go Decay (NGD) pathway (Wohlenberg *et al*, 2026). Additionally, eIF5A has also been found binding to inactive and mRNA-free dormant ribosomes that protect the translation machinery from degradation. Thus, eIF5A was found in the E-site of dormant ribosomes in zebrafish and Xenopus eggs embryos (Leesch *et al*, 2023), in *Schizosaccharomyces pombe* cells under glucose starvation (Gluc *et al*, 2026), and in puromycin-treated neuronal cells (Jiang *et al*, 2026). However, it is still unclear whether eIF5A acts as a stabilizing factor of inactive ribosomes or as a stimulating factor for the reactivation of the dormant ribosomes (Gluc *et al*, 2026). Overall, we still lack a comprehensive understanding of all eIF5A functions in relation to the ribosome, which is relevant as eIF5A is a crucial factor in several diseases in which translation plays a significant role, including several neurodevelopmental syndromes (Faundes *et al*, 2021; Ganapathi *et al*, 2019; Ziegler *et al*, 2022), neurodegeneration (Smeltzer *et al*, 2021), cancer (reviewed in Nakanishi & Cleveland, 2024; Ning *et al*, 2020), as well as in aging (Hofer *et al*, 2021; Liang *et al*, 2021; Luchessi *et al*, 2008; Ma *et al*, 2020; Zhang *et al*, 2019) and viral infection (Ruhl *et al*, 1993). Furthermore, eIF5A has also been linked to stress responses (Li *et al*, 2010; Seoane *et al*, 2024), where translation is profoundly remodeled. Specifically, it is involved in plant thermotolerance (Ma *et al*, 2010; Wang *et al*, 2012; Xu *et al*, 2011) and has a role in translation inhibition and the formation of PBs and SGs under arsenite stress in mammalian cells (Li *et al*, 2010) and under heat-shock in yeast (Seoane *et al*, 2024). However, the underlying mechanism remains unknown.

Here, we discovered that eIF5A is induced and is necessary for a quick response to heat-stress in yeast cells. We show that, under severe heat stress, ribosomes cannot progress without functional eIF5A. Ribosome stalling in eIF5A mutants impacts both, the formation of PBs and SGs, and the synthesis of chaperones. Under mild heat shock, the lack of functional eIF5A primarily affects heat-stress proteins that undergo extensive synthesis via high codon optimality, such as the Hsp70 Ssa1 and Hsp12 chaperones. This new role of eIF5A appears to be direct, resulting in a delayed Hsf1 inhibitory feedback loop and the accumulation of protein aggregates in the cells, thereby reducing cellular viability to heat stress. Overall, our results demonstrate that eIF5A plays a key role in conferring heat stress resistance by binding to and promoting the rapid and efficient synthesis of codon-optimal chaperones. Consequently, eIF5A emerges as a pivotal regulator of cellular proteostasis, shedding light on its significant involvement in neurological disorders and aging.

## RESULTS

### eIF5A is induced and is necessary for heat-stress survival

Although eIF5A is one of the most abundant proteins in yeast cells, we found that its protein levels increased around 2-3 times under mild heat-shock at 37°C and under more severe stress at 41°C. This increase in protein levels was already detected at 30 min upon both heat-shocks and was maintained at longer times of incubation at high temperature (Fig. 1A,B). Increase in protein levels was primarily due to protein stabilization since the mRNA levels of both eIF5A isoforms only increased after prolonged heat shock (Fig. 1C).

**Figure 1.**
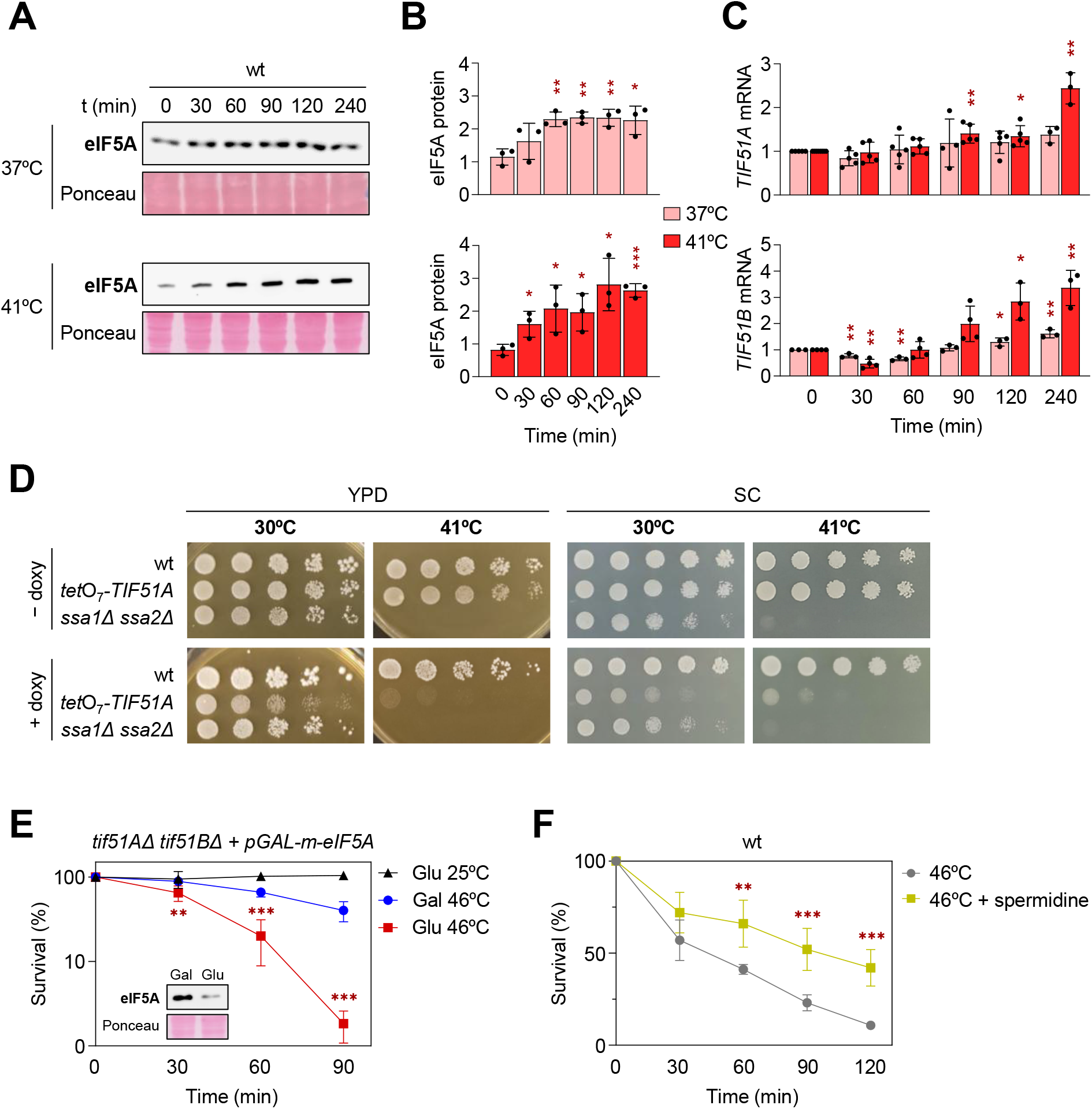
eIF5A is induced and is necessary for survival under heat stress. **(A–C)** Wild type (wt) yeast cells were grown until exponential phase in YPD at 25°C and then, shifted to 37°C or 41°C for the indicated times. **(A)** Representative western blot of eIF5A protein levels in wt cells at 37°C (top) and 41°C (bottom) using an anti-eIF5A antibody. Ponceau S staining is shown as loading control. **(B)** Quantification of eIF5A protein levels from **(A)** normalized to Ponceau S. **(C)** Relative mRNA levels of *TIF51A* and *TIF51B* determined by RT-qPCR and normalized to endogenous *ACT1* mRNA. **(D)** Ten-fold serial dilution spot assays. Wild type, the eIF5A-doxycycline depletable *tet*O_7_ *-TIF51A* strain and the *ssa1Δ ssa2Δ* chaperone double mutant were spotted onto YPD and SC plates with or without doxycycline (0.1 µg/mL) and incubated at the indicated temperatures. **(E)** Survival assay using the *tif51AΔ tif51BΔ* double mutant strain expressing murine eIF5A under a galactose-inducible promoter (pGAL-m-eIF5A). Cells were grown in galactose (SGal-Trp; Gal) at 25°C and shifted to glucose medium to deplete eIF5A (SC-Trp; Glu) for 6 h at 25°C prior to a 46°C heat shock. Control cells were maintained in galactose. The inset confirms eIF5A depletion by western blot using an anti-eIF5A antibody, with Ponceau S staining as loading control. Survival was determined by colony counting and is represented on a logarithmic scale. **(F)** Spermidine supplementation rescues heat shock sensitivity. Wild type cells were cultured in YPD with or without 4 mM spermidine at 25°C to the exponential phase and then shifted to 46°C for the indicated times. Cell viability was determined by colony counting. **Data information:** In (B,C,E,F), data are presented as mean ± SD of independent biological experiments (n = 3 for B; n = 3 to 5 for C; n = 4 to 6 for E; n = 4 to 5 for F). Individual data points are plotted in (B, C), and omitted in (E, F) to preserve graphical clarity. Statistical significance was assessed using a paired two-tailed Student’s t-test. In (B) and (C), asterisks (*P < 0.05, **P < 0.01, ***P < 0.001) indicate significant differences compared to time 0 at the same temperature. In (E), asterisks indicate significant differences between galactose and glucose conditions at 46°C at each corresponding time point. In (F), asterisks indicate significant differences between untreated and spermidine-supplemented cells at each corresponding time point. Exact P values for all comparisons are provided in the associated Source Data file.

We asked then whether the need of more eIF5A protein was connected to a positive effect of the protein in the tolerance to heat stress. Due to the essentiality of eIF5A in all eukaryotes (Park *et al*, 1994; Prieto-Díez *et al*, 2026; Schnier *et al*, 1991), we addressed this first by using a yeast strain with conditional expression of *TIF51A* gene under the control of doxycycline repressible *tet*O_7_ promoter (Bellí *et al*, 1998). The partial depletion of eIF5A protein by doxycycline addition severely reduced the growth of the cells at 41°C, similarly to the growth defect observed in a yeast strain that lacks the highly expressed and partial redundant Hsp70 proteins Ssa1 and Ssa2 (Nelson *et al*, 1992) (Fig. 1D). Second, we used a yeast strain with double deletion of the *TIF51A* and *TIF51B* genes that expressed mouse eIF5A, which complements lack of yeast eIF5A, under the control of a galactose inducible and glucose repressible promoter (Clement *et al*, 2003), and tested survival under severe heat-stress (46°C). We confirmed that survival of eIF5A depleted yeast cells was still not reduced after 90 minutes of transferring the cells from galactose to glucose media at 25°C, according to previous results (Li *et al*, 2014). However, under heat stress conditions (46°C), yeast cells expressing eIF5A (in galactose media) had significantly higher survival than cells in which eIF5A had been depleted (glucose media) (Fig. 1E). Moreover, we observed that the addition of the polyamine spermidine, which is essential for eIF5A activation by hypusination (Park, 2006), increased the tolerance of wild type yeast cells to severe heat-shock, supporting the positive role of eIF5A to heat-stress survival (Fig. 1F).

Altogether, our results suggest that the levels of the eIF5A protein modulate the survival of yeast cells at high temperatures. These results are in line with previous observations that over-expression of eIF5A increases and its depletion decreases tolerance to heat-stress in plants (Ma *et al*, 2010; Wang *et al*, 2012; Xu *et al*, 2011); and with the correlation between the loss of active eIF5A and heat-stress-induced cell death observed in human colon and pancreatic cancer cells (Gosslau *et al*, 2009; Takeuchi *et al*, 2002).

### Translational but not transcriptional reprogramming depends on eIF5A under heat stress

As mentioned above, previous studies from our group have shown defects in the inhibition of global translation during heat shock in eIF5A-depleted yeast cells (Seoane *et al*, 2024). Moreover, a study with mammalian cells showed that hypusinated-eIF5A is necessary for the disassembly of polysomes and the formation of PB and SG under arsenite stress (Li *et al*, 2010). Given the important role of eIF5A in thermotolerance and survival, we aimed to conduct a comprehensive analysis of eIF5A function in response to heat stress. First, we studied the transcriptional reprogramming that occurs in yeast cells when transferred from 25°C to 37°C or 41°C. To do this, we employed two temperature-sensitive yeast strains, *tif51A-1* and *tif51A-3*, containing one-or two-point mutations in the *TIF51A* gene that encodes eIF5A, respectively. These eIF5A mutant proteins get rapidly degraded at 37°C or a higher temperature, with Tif51A-3 protein degrading faster and resulting in a stronger growth defect in the *tif51A-3* mutant (Fig. EV1A) (Li *et al*, 2014). Similar or even higher levels of induction of various heat-stress-induced genes were observed in the eIF5A mutants than in the wild type under both heat stress conditions (Fig. 2A). With respect to Hsp70 chaperones, in all strains, induction was higher for *SSA3* and *SSA4* (the HSP70 genes originally classified as inducible), and lower for *SSA1* and *SSA2* (which are expressed under normal conditions and originally classified as non-inducible) (Fig. 2A). Similarly, the mRNA levels of other chaperones such as Hsp12, Hsp26, Hsp42, Hsp104 and Hsp150, as well as other stress response proteins such as the catalase Ctt1, the hexokinase Hxk1 and the transcription factor Hsf1, were induced to the same or a greater extent in cells eIF5A-depleted than in wild type cells under heat stress. Regarding the repression of house-keeping genes, the mRNA levels of the ribosomal proteins Rpl28 and Rpl30 were reduced in both eIF5A mutants and wild type cells under heat stress. However, after 2–3 hours of heat stress, we observed that the initial mRNA levels of many stress-induced and repressed genes were recovered in wild type cells. In contrast, stress induced mRNA levels remained higher for longer in *tif51A-1* and *tif51A-3* mutants (Fig. 2A). In summary, eIF5A mutants were found to be able to reshape the transcriptional landscape in response to heat stress; however, recovery back to initial levels appears to be slower in the eIF5A mutants.

**Figure 2.**
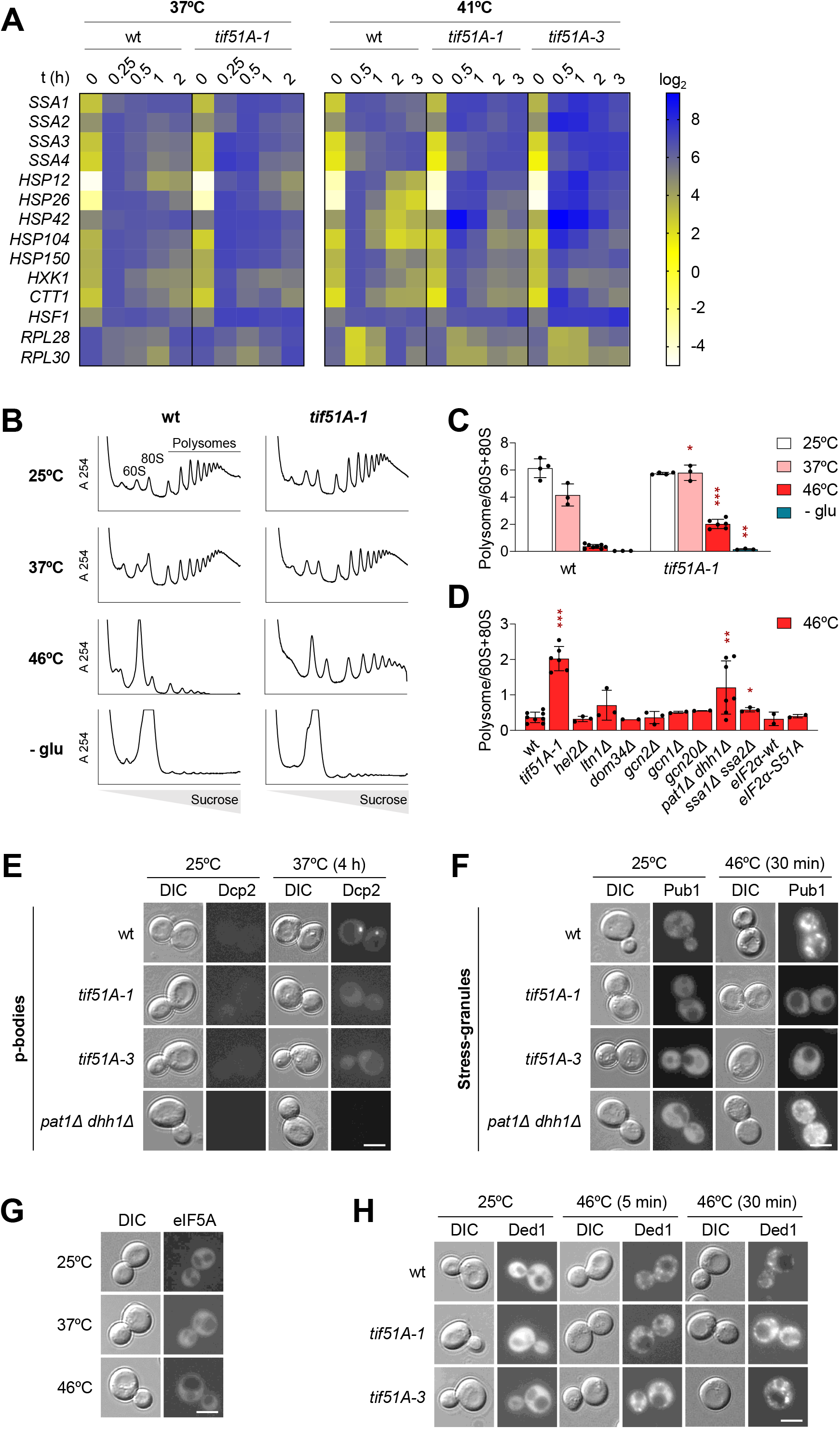
Effect of eIF5A depletion over the transcriptional and translational reprogramming under heat stress. **(A)** Heatmap of relative mRNA expression over time. Wild type (wt), *tif51A-1*, and *tif51A-3* strains were grown in YPD at 25°C to the exponential phase and then shifted to 37°C or 41°C for the indicated times. mRNA levels were determined by RT-qPCR using endogenous *ACT1* mRNA for normalization. Data were normalized for each gene to its highest value observed in the wt strain, set to 100%. Then, a log_2_ transformation was applied to the mean of three biological replicates. **(B)** Polysome profiles of wt and *tif51A-1* strains. Cells were cultured in SC medium at 25°C and subjected to the following stress conditions: 37°C for 10 min (labeled as 37°C), 37°C for 4 h followed by 30 min at 46°C (46°C), or the same sequential heat stress with glucose depletion during the final 15 min (-glu). A representative profile from three independent experiments is shown. **(C)** Quantification of the polysome/(60S+80S) area ratio calculated from the profiles in **(B)**. **(D)** Polysome/(60S+80S) area ratio for the indicated strains. Cells were grown at 25°C in SC medium to the exponential phase and shifted to 37°C for 4 h followed by 30 min at 46°C. **(E, F)** Fluorescence microscopy of p-bodies and stress granules. The wt, *tif51A-1*, *tif51A-3*, and *pat1Δ dhh1Δ* strains expressing either Dcp2-RFP **(E)** or Pub1-mCherry **(F)** were grown in SC-Leu or SC-Ura, respectively. Cells were grown at 25°C and shifted to 37°C for 4 h **(E)**, or to 37°C for 4 h followed by a heat shock at 46°C for 30 min **(F). (G)** Fluorescence microscopy evaluating the localization of endogenously tagged GFP-eIF5A in wt cells at 25°C, after incubation at 37°C for 4 h or after 4 h at 37°C followed by a 30-min heat shock at 46°C. Representative images from three independent experiments are shown for all microscopy panels. **(H)** Fluorescence microscopy evaluating endogenous Ded1-GFP localization in wt and the temperature-sensitive mutants *tif51A-1* and *tif51A-3* at 25°C or after a direct heat shock at 46°C for 5 or 30 min. Scale bars: 5 µm. **Data information:** In (C, D), data are presented as mean ± SD of independent biological experiments (n = 3 to 8 for C; n = 2 to 8 for D). Individual data points shown as dots. Statistical significance was assessed using an unpaired two-tailed Student’s t-test for conditions with n ≥ 3. Exact P values for all comparisons are provided in the associated Source Data file. In (C), asterisks (*P < 0.05, **P < 0.01, ***P < 0.001) indicate significant differences between the *tif51A-1* mutant and the wt under the same condition. In (D), asterisks denote significant differences for each strain compared to the wt at 46°C.

Next, we analyzed the translational reprogramming by studying the polysomal profiles over the heat stress. In wild type cells grown at 25°C and transferred to 37°C, a reduction in the polysomal fractions was observed, reaching a minimum after 10 min of heat stress. The original levels recovered between 15 and 20 min after stress, indicating a transient and partial reduction in global translation (Figs. 2B,C and EV1B,C). However, no reduction in the polysomal fractions was observed in *tif51A-1* mutant cells at any point during heat stress at 37°C (Figs. 2B,C and EV1B,C). Because translation inhibition under mild heat stress (37°C) is slight and transient, we aimed to investigate the response to severe heat stress at 46°C. To do this, cells were grown at permissive temperature (25°C) and subsequently depleted of eIF5A by incubation of 4 h at 37°C. These cells were then exposed to 30 min of severe heat stress at 46°C. Wild type cells showed a significant reduction in the polysomal levels, whereas eIF5A mutant cells exhibited much less inhibition of global translation under severe heat stress (Fig. 2B,C). Although we have previously shown that *tif51A-1* mutant cells do not lose viability after incubation of 6 hours at 37°C (Li *et al*, 2014), we wondered whether these cells were still responsive after 30 min incubation at 46°C. To check this, we additionally incubated cells in a free glucose media, a stress condition that produces a major inhibition of global translation (Ashe *et al*, 2000). Indeed, in wild type cells this additional stress produced an additional reduction in the polysomal levels (90.8% reduction in polysomal/80S+60S ratio). eIF5A depleted cells reacted to this stress showing an important inhibition of global translation (92.4% reduction in polysomal/80S+60S ratio) (Fig. 2B,C). This result pointed to a specific defect in translational reprograming associated to heat stress. To demonstrate that the lack of eIF5A is the primary cause of the inability to inhibit translation under heat stress, we obtained polysomal profiles in the *tet*O_7_-*TIF51A* yeast strain. Consistent with the results obtained with temperature sensitive eIF5A mutants, global translation was not inhibited by heat-stress upon doxycycline-induced eIF5A depletion (Fig. EV1D). This result correlated well with the growth defects of doxycycline-treated *tet*O_7_-*TIF51A* cells under heat stress (Fig. 1D).

To have some clues about how eIF5A controls translation under heat stress, we analyzed the effect on translation of mutations in proteins functionally related to eIF5A. First, given the recent role attributed to eIF5A in ribosome-associated quality control pathways, we investigated whether mutants in the RQC factors Hel2 (homologue to human ZNF598 (Juszkiewicz *et al*, 2018), Ltn1 (homologue to human LTN1) (Bengtson & Joazeiro, 2010), and in the ribosome recycling factor Dom34 (homologue to human Pelota) (Becker *et al*, 2012), were able to reprogram translation upon heat shock. However, none showed defects in global translation inhibition (Fig. 2D and Appendix Fig. S1A), indicating that, the RQC pathway is not required for translational reprogramming under heat stress. Additionally, we confirmed that both a non-phosphorylable eIF2alpha mutant (sui S51A) and a Gcn2 kinase mutant were able to inhibit global translation under 46°C heat stress (Fig. 2D and Appendix Fig. S1B). Moreover, mutants in the Gcn2 activating factors Gcn1 and Gcn20 (Vazquez De Aldana *et al*, 1995) were also inhibiting translation normally under severe heat stress (Fig. 2D and Appendix Fig. S1A). This again excludes a role of the described ribosome collision sensors Hel2 and Gcn1 (Inada & Beckmann, 2024) in regulating global translation under heat stress. Further, a slight defect in translation inhibition was observed in the double mutant *ssa1Δ ssa2Δ*, although it is sensitive to heat stress, thus discarding a main role for these Hsp70 chaperones in the repression of global translation. Meanwhile and as previously reported (Coller & Parker, 2005), a double mutant in the mRNA decay and general translation repression factors, Dhh1 and Pat1 (*dhh1Δ pat1Δ*), was not able to inhibit global translation under heat stress (Fig. 2D and Appendix Fig. S1C). Furthermore, the absence of global translation inhibition in the *dhh1Δ pat1Δ* and eIF5A mutants was correlated with defects in the formation of PB under mild heat stress and SG under severe heat stress, and also correlated with growth defects under heat stress. However, only eIF5A mutants showed a complete inability to form both mRNA heat storage condensates, correlating with the stronger defect in global translation inhibition (Figs. 2E,F and EV1E,F and Appendix Fig. S1D). Additionally, although the localization of eIF5A in SG has been described (Smeltzer *et al*, 2021), we did not observe the formation of condensates with a GFP-eIF5A fusion under heat stress (Fig. 2G).

Next, we checked whether the lack of eIF5A impaired the condensation of the Ded1 helicase, as Ded1 condensation under severe heat stress has been shown to promote house-keeping mRNAs repression and, therefore, global translation inhibition (Iserman *et al*, 2020). However, Ded1 protein condensates were visualized quickly after 5 min of 46°C incubation in yeast cells both, with and without eIF5A. Indeed, more abundant and intense Ded1 condensates were visualized in eIF5A mutant cells than in wild type cells (Fig. 2H).

Altogether, these results indicate that eIF5A is involved in the translation inhibition, but not in the initial transcriptional reprogramming, under mild and severe heat stress, which in turn leads to the formation of the mRNA storage PB and SG granules, respectively. This function appears to be specific to heat stress, as translation inhibition was only minimally impacted under glucose depletion in eIF5A depleted cells. The eIF5A-controlled mechanism of global translation inhibition appears to be independent of the RQC pathway, Hel2 and Gcn1 recognition of stalled disomes, and Ded1 helicase condensation.

### eIF5A depletion causes ribosome stalling upon heat stress

It is generally assumed that, under stress, reprogramming of translation releases ribosomes from house-keeping mRNAs, making them available to translate newly synthesized mRNA (Crawford & Pavitt, 2018; Hernandez-Elvira & Sunnerhagen, 2022; Yamasaki & Anderson, 2008). If this is true, then in eIF5A mutants, where global translation is not inhibited by heat stress, stress induced mRNAs will not be found in the polysomal fractions. To investigate this, we studied the distribution of mRNAs along the polysomal profile in wild type and eIF5A mutant cells. The distribution of the mRNA encoding the house-keeping ribosomal protein Rpl28 showed the expected pattern for wild type cells, being mainly associated with the polysomal fractions at 25°C and quickly moving, partially at 37°C and mostly at 46°C, from polysomal to monosomal and lighter fractions (Fig. 3A). However, in line with the global polysomal profiles (Fig. 2B), *RPL28* mRNA was significantly retained in the polysomal fractions in the eIF5A mutant (Fig. 3A). We then studied the mRNA distribution of the heat stress-induced chaperones *SSA1* and *HSP12*; as well as the mRNA distribution of other non-chaperone heat-stress induced proteins, such as the hexokinase *HXK1* and the catalase *CTT1* mRNAs. We observed a similar distribution pattern for all of these mRNAs, which were predominantly present in the polysomal fractions under mild heat stress in both wild type and eIF5A mutant strains. In the case of the Hsp70 *SSA1* mRNA, which is highly expressed under non-stress conditions and further induced under heat stress, we also observed it to be mostly associated with polysomal fractions at 25°C in both strains, as expected. Under severe heat shock, the four heat stress mRNAs showed a similar or stronger association with polysomal fractions in eIF5A mutant cells than in wild type cells (Fig. 3A). Therefore, these results demonstrated that, although ribosomes are defectively maintained in association with house-keeping mRNAs in eIF5A mutants under heat stress, they are not limiting, as heat stress mRNAs are associated with the polysomal fraction in eIF5A mutants to a similar extent as in wild type cells under mild heat stress. An even greater association was observed under severe heat stress.

**Figure 3.**
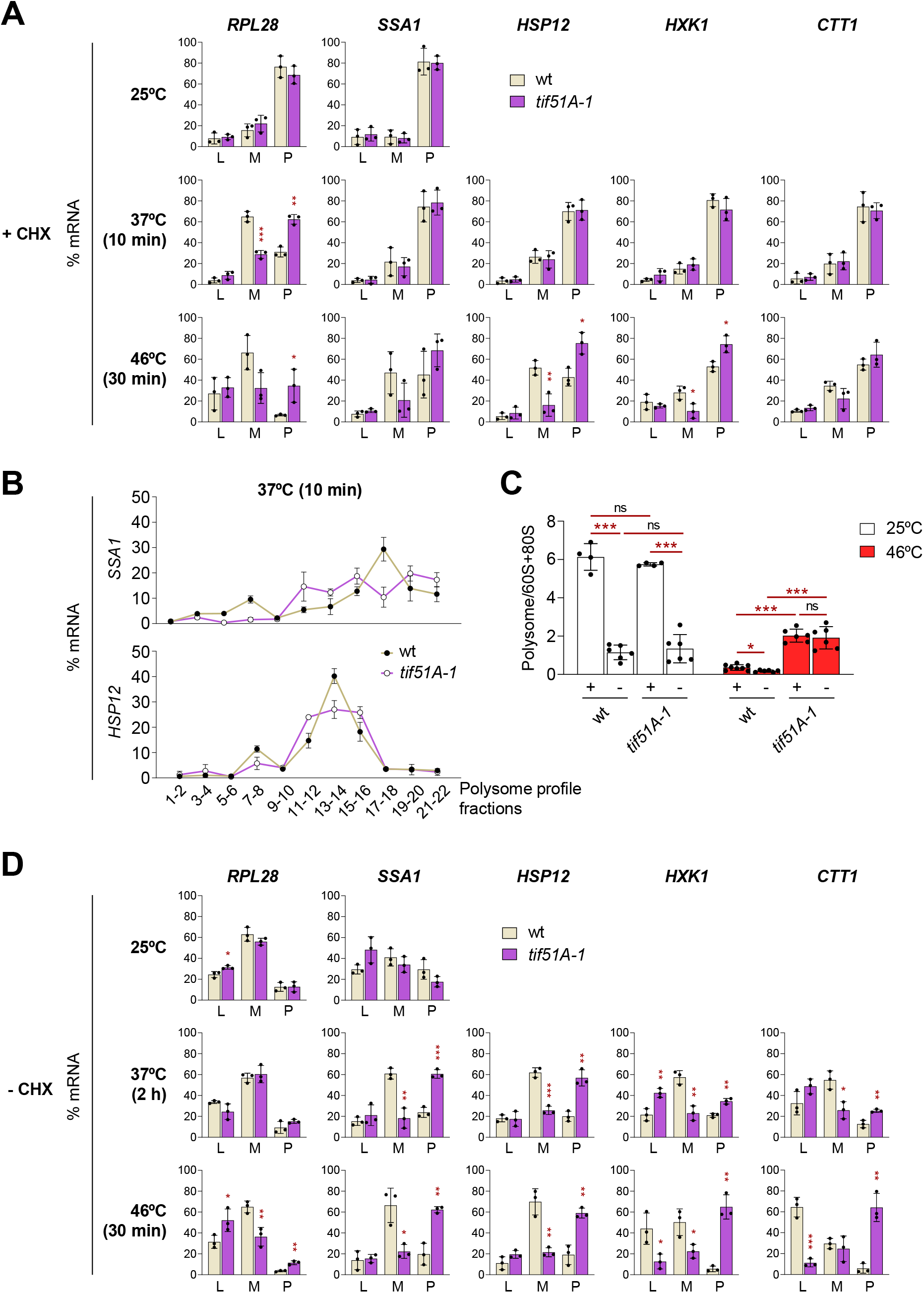
In eIF5A mutants, ribosomes stall to different extents depending on the mRNA and the severity of the heat stress. **(A)** Relative mRNA distribution in polysome profiles from wild type (wt) and *tif51A-1* strains treated with cycloheximide (CHX) to arrest translating ribosomes. Cells were cultured in SC medium at 25°C to the early exponential phase and then maintained at 25°C, shifted to 37°C for 10 min, or shifted to 37°C for 4 h followed by 30 min at 46°C. Gradient fractions were pooled into free RNA (L), monosomes (M), and polysomes (P), and the relative levels of the indicated mRNAs were analyzed by RT-qPCR. **(B)** Relative distribution of *SSA1* and *HSP12* mRNAs across the polysome profile in wt and *tif51A-1* strains. Cells were grown and treated with CHX as in **(A)** and subjected to heat stress at 37°C for 10 min. Fractions were pooled in pairs prior to RNA extraction and RT-qPCR analysis. **(C)** Quantification of the polysome/(60S+80S) area ratio from polysome profiles obtained from wt and *tif51A-1* strains. Cells were grown at 25°C and then incubated at 37°C or 46°C as described in **(A),** and treated with (+) or without (−) CHX prior to harvesting. **(D)** Relative mRNA distribution in polysome profiles analyzed as in **(A)**, but without CHX treatment to promote ribosome run-off. Cells were cultured in SC medium at 25°C to the early exponential phase and then maintained at 25°C, shifted to 37°C for 2 h, or shifted to 37°C for 4 h followed by 30 min at 46°C. **Data information:** In (A–D), data are presented as the mean ± SD from independent biological experiments (n = 3 for A, B, and D; n = 3 to 8 for C). Individual data points are plotted as dots in (A, C, D) and omitted in (B) to preserve graphical clarity. In (A, B, D), the amount of mRNA in each pool or fraction is expressed as a percentage of the total amount of that specific mRNA across the entire gradient. Statistical significance was determined using a two-tailed unpaired Student’s t-test (\**P* < 0.05, \*\**P* < 0.01, \*\*\**P* < 0.001). In (A) and (D), asterisks indicate significant differences relative to the wt strain within each specific gradient region (L, M, or P). In (C), asterisks located directly above the bars indicate significant differences between the *tif51A-1* mutant and the wt strain under the same temperature and CHX condition. Asterisks over horizontal lines indicate significant differences between the +CHX and −CHX treatments within the same strain and temperature. Exact *P* values for all comparisons are provided in the associated Source Data file.

As we analyzed all polysomal fractions together, we wondered whether a possible limitation in ribosome availability could cause a movement of mRNAs from heavier to lighter polysomal fractions, which would not be detectable with our previous analysis. Therefore, we studied the distribution of heat-stress induced mRNAs along the different polysomal fractions. In wild type cells, *SSA1* mRNA peak was detected in fractions 17-18, whereas the shorter *HSP12* mRNA peak was detected in fractions 13-14 at 37°C (Fig. 3B). In eIF5A mutant cells, the peaks for both mRNAs were in similar fractions to those in the wild type, however, the mRNAs did not show a narrow peak; instead, the mRNAs were distributed at similar levels in the heavier and lighter fractions around the peak, suggesting defects in translation elongation (Fig. 3B).

The association of heat-stress mRNAs with polysomal fractions, and thus with putatively active translation and synthesis of proteins for survival this stress, was contradictory with the low heat-stress tolerance of eIF5A mutants, and suggested that ribosomes may be stalled. To investigate this further, we obtained new polysomal profiles without adding cycloheximide (–CHX). Under these conditions, active ribosomes can continue to elongate during the extraction process. Consequently, they will run-off and the mRNA will move from the polysomal to the monosomal and free fractions. Conversely, inactive elongating ribosomes remain stalled and mRNA stays in the polysomal fractions. To confirm that the ribosome run-off was occurring under –CHX conditions, we calculated the polysomal/80S+60S ratio after obtaining the polysomal profiles, and observed the expected drop in this ratio in both wild type and eIF5A mutant cells at 25°C. However, a drop in this ratio was not observed in the eIF5A mutant cells incubated at 46°C, while it was observed in wild type cells (Figs. 3C and EV2A). This result confirmed a general problem in ribosome elongation under severe heat stress in eIF5A depleted cells. We next studied the distribution of house-keeping and heat stress mRNAs in the polysomal profile under run-off conditions. For mild heat stress, we incubated the cells for 2 hours at 37°C. At this time, the levels of heat stress mRNAs are sufficiently high and are still being actively translated, while the level of eIF5A protein has decreased significantly in the temperature sensitive strain *tif51A-1* (Figs. 2A, EV2A and EV1A). Under these conditions, the run-off of ribosomes in *SSA1* and *HSP12* mRNAs was low in eIF5A depleted cells, as indicated by a still higher percentage of these mRNAs in the polysomal fractions compared to wild type samples. Ribosome run-off was higher in *HXK1* and *CTT1* mRNAs than in the chaperone mRNAs, and a complete ribosome run-off was observed in *RPL28* mRNA in eIF5A mutants. However, under severe heat shock, defects in ribosome run-off were detected for all the studied mRNAs, including *RPL28* mRNA (Fig. 3D).

It has been proposed that disomes formed from colliding ribosomes under stress conditions are recognized by Gcn1, which activates the Gcn2 kinase, resulting in the phosphorylation of eIF2alpha and the subsequent inhibition of translation initiation (Chang & Choe, 2026). eIF2alpha is also phosphorylated under heat stress (Grouši *et al*, 2009), although this is not sufficient to cause global translation inhibition. As our results showed different ribosome dynamics between wild type and eIF5A mutants, we wondered whether this causes differences in signaling for eIF2alpha phosphorylation. In wild type cells, we observed a strong eIF2alpha phosphorylation signal under 30 min of 46°C severe heat stress. However, only a weak eIF2alpha phosphorylation signal was observed in eIF5A mutant cells under severe stress (Fig. EV2B,C). This was not due to a general loss of signaling capacity, as a control signal for heat stress-mediated Hog1 phosphorylation (Winkler *et al*, 2002) was observed in the eIF5A mutants (Fig. EV2B,C). Therefore, these results indicate that, in cells under heat stress, eIF5A-depletion disturbs signaling for eIF2alpha phosphorylation and this may be linked to different ribosome dynamics and/or disome conformation. Furthermore, defective translation initiation inhibition due to low eIF2alpha phosphorylation, together with a ribosome elongation defect, explain why eIF5A mutants do not exhibit global translation inhibition under heat stress.

In summary, our results analyzing ribosome movement are consistent with those of translational reprogramming and PB/SG formation and predict different outcomes. Thus, under severe heat stress, ribosomes in eIF5A-depleted cells, are stalled in elongation across all mRNAs. Therefore, ribosomes are retained on house-keeping mRNA, impairing their condensation into SGs. On the other hand, stalled ribosomes on newly transcribed mRNA must impede the synthesis of heat stress proteins. Under mild heat stress, the effect of eIF5A depletion on translation is milder and mRNA-dependent. Quick and transient inhibition of global translation does not occur in eIF5A mutant cells. Moreover, elongating ribosomes appear to be partially inactive in heat-stress-induced mRNA at 37°C, with a more significant run-off defect observed in *SSA1* and *HSP12* mRNA than in *HXK1* and *CTT1* mRNA (Fig. 3D). These differences must also impact the synthesis of the corresponding proteins differently.

### eIF5A is necessary for the efficient synthesis of chaperones under heat-stress

Our previous results predicted that synthesis of heat stress proteins must be compromised in eIF5A mutants. To investigate this, we examined the production of heat shock proteins in wild type and *tif51A-1* and *tif51A-3* mutants under eIF5A depletion conditions, both at mild heat stress (cells grown at 25°C and then incubated for four hours at 37°C) and at severe heat stress (cells grown at 25°C and then preincubated for four hours at 37°C to fully deplete eIF5A, followed by severe heat shock treatment involving incubation for 30 minutes at 46°C). Due to the lack of specific antibodies, we used the C-terminal tags integrated into the loci of heat stress-induced proteins. When examining Hsp70 chaperones, we observed huge Ssa1 protein synthesis under mild and severe heat stress in wild type cells; however, only slight or no increase in Ssa1 protein levels was observed in eIF5A-depleted cells. Nevertheless, a similar increase in *SSA1* mRNA levels was observed in wild type and eIF5A mutant strains under mild and severe heat stress, respectively, confirming a translational defect in the mutants (Fig. 4A,B,C). Ssa2 protein levels increased slightly in wild type cells at 37°C and 46°C, but no increase was observed in eIF5A mutants under either condition, despite similar increases in *SSA2* mRNA levels in both wild type and eIF5A mutant cells. Of the Hsp70 chaperones, *SSA4* mRNA showed the highest increase at 46°C and modest induction at 37°C. However, in accordance with previous data (Boopathy *et al*, 2023), only modest levels of Ssa4 protein were obtained in wild type cells at both temperatures. In the *tif51A-1* mutant cells, the Ssa4 protein increased to a similar extent as in the wild type cells under mild heat stress, and lower Ssa4 protein levels were observed in the more severe eIF5A mutant, *tif51A-3*. Under severe heat stress, however, lower levels of Ssa4 protein were observed in both eIF5A mutants compared to the wild type, although the corresponding *SSA4* mRNA levels were similar in the mutants (Fig. 4A,B,C). The results for Ssa3 protein were similar to those for Ssa4. Similar Ssa3 protein and mRNA levels were observed in cells with or without eIF5A under mild stress, but lower protein levels were detected in the eIF5A mutant under severe heat stress (Fig. EV3A,B,C).

**Figure 4.**
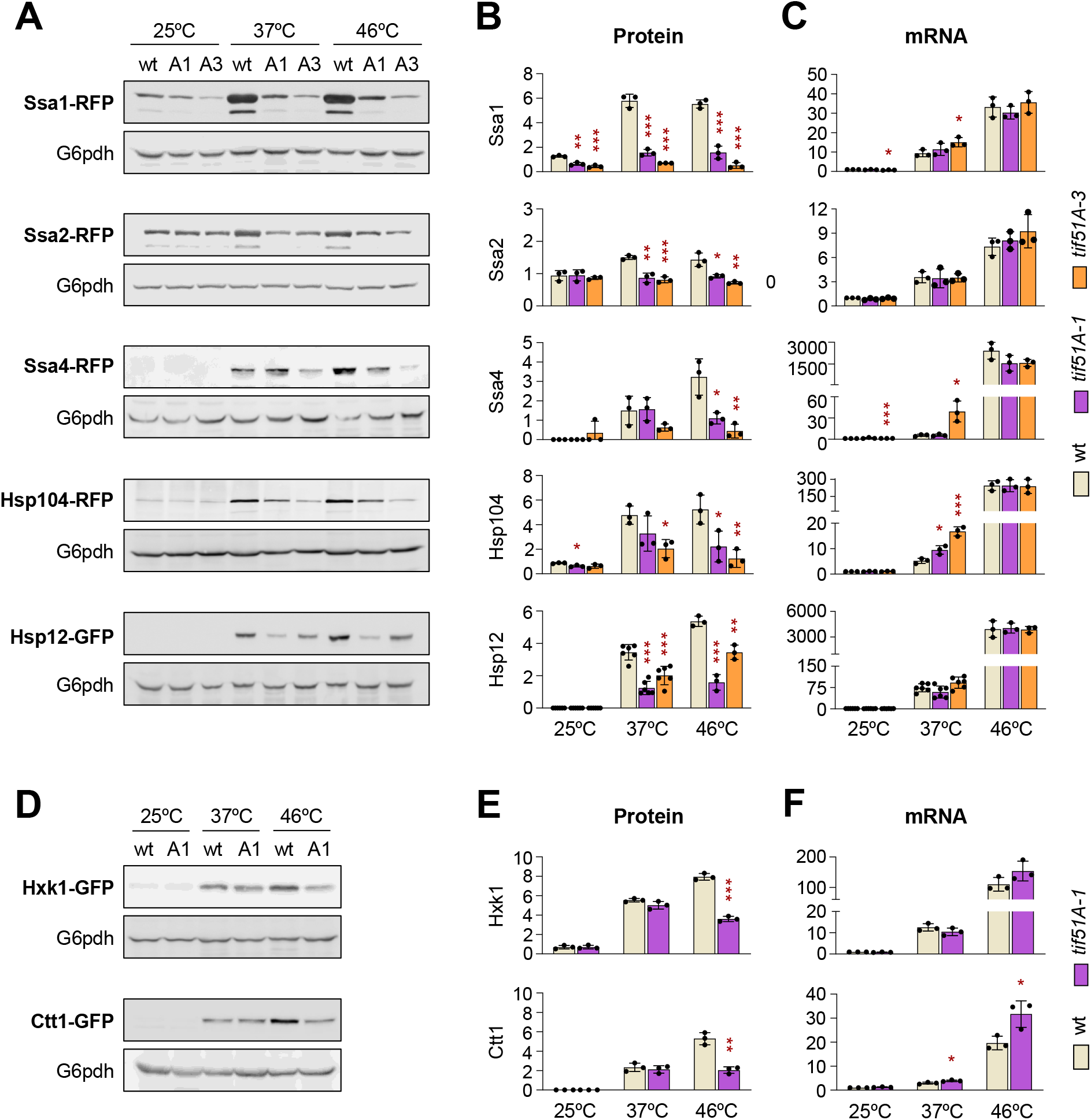
Lack of functional eIF5A impairs the synthesis of specific chaperones under heat stress. **(A,B,C)** Wild type (wt), *tif51A-1* (A1) and *tif51A-3* (A3) strains were cultured in YPD at 25°C to the early exponential phase and shifted to 37°C for 4 h or to 37°C for 4 h followed by 30 min at 46°C. **(A)** Representative western blots evaluating the protein levels of endogenously tagged Ssa1, Ssa2, Ssa4, Hsp104, and Hsp12 using anti-RFP or anti-GFP antibodies. G6pdh was used as a loading control. **(B)** Quantification of protein levels from the western blots shown in **(A)**, normalized to the G6pdh loading control. **(C)** Relative mRNA levels of *SSA1*, *SSA2*, *SSA4*, *HSP104*, and *HSP12* determined by RT-qPCR. **(D)** Representative western blots evaluating Hxk1 and Ctt1 protein levels in wt and *tif51A-1* cells, using an anti-GFP antibody and G6pdh as a loading control. **(E)** Quantification of relative protein levels from the western blots in **(D). (F)** Relative mRNA levels of *HXK1* and *CTT1* determined by RT-qPCR, in wt and *tif51A-1* cells. **Data information:** For all RT-qPCR experiments (C, F), mRNA levels were normalized to endogenous *ACT1* mRNA and are expressed relative to the wt strain at 25°C. In (B, C, E, F), data represent the mean ± SD from independent biological experiments (n = 3 to 6). Individual data points are plotted as dots. Statistical significance was assessed using a two-tailed unpaired Student’s t-test (\**P* < 0.05, \*\**P* < 0.01, \*\*\**P* < 0.001). Asterisks indicate statistically significant differences between the mutant and the wt strain under the same temperature condition. Exact *P* values for all comparisons are provided in the associated Source Data file.

We investigated next the synthesis of other chaperones and other stress responsive proteins. The results for the chaperones Hsp12, Hsp26, Hsp104 and Hsp150 were similar to those obtained with Ssa1 and Ssa2: lower protein levels under mild and severe heat stress were observed in eIF5A mutant with respect to wild type cells. These lower chaperone protein levels in eIF5A mutant under mild heat stress were not due to lower levels of the corresponding mRNAs (Figs. 4A,B,C and EV3A,B,C). However, for the stress induced Hxk1 and Ctt1 proteins, similar increased levels were observed in eIF5A mutants and wild type cells under mild heat stress, and significant lower protein levels were observed in the mutant cells under severe heat stress (Fig. 4D,E,F). Thus, for these last two genes, the expression pattern was similar to that of the Ssa3 and Ssa4 chaperones.

To rule out the possibility of artefacts caused by the type of tag used in each case, we confirmed that the results were very similar when using GFP or HA tags with Hsp12 and Hsp26 proteins (Figs. 4A,B,C and EV3A-F), and for Ssa1 fused to RFP or GFP (Figs. 4A and EV3G,H,I,J).

Among all proteins analyzed, Ssa1 was the one most abundantly produced under heat stress, as it was observed an increase of around 6-fold under mild and severe heat stress in wild type cells. This means the synthesis of an enormous amount of protein under heat stress, as Ssa1 is one of the most abundant proteins in the cells under normal conditions (around 200000 copies of protein per cell, *Saccharomyces* Genome Database; https://www.yeastgenome.org/). As Ssa1 was one of the stress responsive proteins whose increase was more affected by eIF5A depletion (Fig. 4A,B,C), we chose Ssa1 for further studies. First, we confirmed by western and fluorescence microscopy that the Ssa1 protein levels were lower in eIF5A mutants than in the wild type along a kinetic of heat stress at 37°C (Fig. EV3G,H,I). Also according to the previous results, the *SSA1* mRNA levels were similar or even higher for the eIF5A mutant than the wild type (Fig. EV3J), supporting a defect in mRNA translation. We also confirmed that the lower levels of Ssa1 protein were not due to a lower stability of the protein in eIF5A depleted cells (Fig. EV3K,L). Additionally, we confirmed that defects in the Ssa1 synthesis were not linked to the use of temperature-sensitive eIF5A mutants, as similar defects were observed in the tetO_7_-*TIF51A* yeast strain in which eIF5A depletion was done by repressing the expression of the *TIF51A* gene by the addition of doxycycline (Fig. EV4A-E).

As the synthesis of the Ssa1 chaperone is the most severely affected in eIF5A mutants under heat stress, and as Ssa1 is also involved in translation (Horton *et al*, 2001), we wondered whether the observed defects in the synthesis of other chaperones in eIF5A mutants were an indirect consequence of the low Ssa1 production under heat stress. To investigate this, we studied Hsp12 synthesis in eIF5A and *ssa1Δ ssa2Δ* mutants containing a high-copy plasmid expressing Ssa1-FLAG. In control cells containing an empty plasmid, we observed lower increases in Hsp12 protein levels under heat stress in both the eIF5A mutant and the double *ssa1Δ ssa2Δ* mutant than in wild type cells (Fig. EV4F). This confirms the role of Ssa1 in synthesizing the Hsp12 chaperone. However, induction of Hsp12 protein levels under stress was not restored in the eIF5A mutant, though it was restored in the *ssa1Δ ssa2Δ* mutant, when a Ssa1-Flag protein was over-expressed. It should be noted that although Ssa1-Flag expression was lower in the eIF5A mutant than in the other strains due to its defects in Ssa1 synthesis, it was still notable and did not improve Hsp12 production (Fig. EV4F). These results were consistent with the observed growth phenotypes, where the *ssa1Δ ssa2Δ* double mutant was complemented by the plasmid-mediated Ssa1-Flag overexpression, whereas no complementation was observed in the eIF5A mutants (Fig. EV4G). Consequently, these findings strongly suggest that defects in eIF5A-mediated chaperone synthesis under heat stress are not merely an indirect consequence of defective Ssa1 synthesis.

In sum, the results of protein synthesis were in accordance with those with polysomes analyzing translation. Thus, under severe heat stress, where ribosomes were stalled in all heat stress mRNAs tested in eIF5A mutant cells, we see that cells are unable to synthesize heat-stress-responsive proteins in the absence of eIF5A. Under mild heat stress, however, the defect was observed in the synthesis of certain stress-induced proteins (e.g. Ssa1, Ssa2, Hsp12, Hsp26, Hsp104 and Hsp150), but not in the synthesis of others (e.g. Ssa3, Ssa4, Hxk1 and Ctt1). These results are also in agreement with the analysis of polysomes, as ribosomes in *SSA1* and *HSP12* mRNAs were found mostly stalled, and therefore did not run-off, and almost no increase in the corresponding proteins was observed in eIF5A mutants under mild heat stress. Furthermore, the defective synthesis of most chaperone due to the stall of ribosomes explains the temperature sensitivity observed in eIF5A mutants.

### Synthesis defects of heat shock proteins in eIF5A-depleted cells are independent of specific peptide motifs but contingent on the coding region

Our results have shown that under severe heat stress there is a general problem in ribosome elongation in eIF5A mutants, while under mild heat stress, the defect seems to be mRNA specific. We aimed to understand why some heat stress mRNAs are most affected than others. Given that eIF5A has been described as a translation elongation factor promoting peptide bound formation when ribosomes encounter codons encoding for consecutive prolines or combination of proline, glycine and charged amino acids (Pelechano & Alepuz, 2017; Schuller *et al*, 2017), we determined whether the most affected proteins had a higher content of eIF5A-dependent peptide motifs. For this, we used the previously described Protein Pause Index (PPI) (Barba-Aliaga *et al*, 2026), which gives an eIF5A-dependent pausing score for each gene by counting the number of tripeptides that cause pausing under eIF5A depletion in the protein encoded sequence, and takes into account the strength pause value of each tripeptide revealed by 5PSeq analysis (Pelechano & Alepuz, 2017). The higher the PPI for the gene, the more likely the translation of the corresponding mRNA will stall upon eIF5A deficiency. In the yeast proteome, the average PPI value is 10.84 and bona-fide targets of eIF5A such as the proline-rich proteins Vrp1 (Gutierrez *et al*, 2013), Bni1 (Li *et al*, 2014) and Tim50 (Barba-Aliaga *et al*, 2024) have a PPIs of 465.9, 217.0 and 48.5, respectively. Ssa1-4 proteins, although having highly similar protein sequences (identities of 96% for Ssa2, 74% for Ssa3 and 74% for Ssa4, with Ssa1), show higher PPIs for Ssa1 (49.01) and Ssa2 (36.31) than for Ssa3 (19.27) and Ssa4 (23.86). Other chaperones with poor synthesis in eIF5A mutant cells at 37°C have lower PPIs, such as Hsp12 (3.82) and Hsp150 (11.23), similar or lower than that of non-affected proteins such as Hxk1 (12.44) and Ctt1 (10.14). Therefore, there is no good correlation between the defect in translation in eIF5A mutants under mild heat shock and the presence of eIF5A-dependent peptide motifs.

However, because the synthesis of Ssa1 protein is one of the most affected, we investigated whether removing eIF5A-dependent motifs renders its synthesis independent of eIF5A. The high PPI score of full Ssa1 protein (642 amino acids) is mostly due to the presence of two pentapeptides putatively dependent on eIF5A (Pelechano & Alepuz, 2017) (Fig. 5A). However, the synthesis of a C-terminal truncated Ssa1 protein (502 amino acids), which does not contain these two motifs (resulting PPI of 10.2), was strongly affected under mild heat stress in eIF5A mutant cells (Figs. 5A–D and EV5A,B). We next obtained shorter Ssa1 version (197 amino acids) with a PPI of 0. A defect in the synthesis of this Ssa1 truncated protein was observed in eIF5A mutants, although this truncated protein showed a more reduced synthesis than the native Ssa1 in wild type cells at 37°C. The synthesis of a truncated Ssa1 protein containing only the N-terminal 83 amino acids (PPI 0) increased slightly under heat stress in wild type cells. However, a slight defect was still detected in eIF5A mutants. Finally, a version with depletion of the full *SSA1* coding sequence (CDS) but expressing the RFP tag under the control of the *SSA1* promoter showed a very low increase in the corresponding protein levels in wild type cells under mild heat stress. Remarkably, higher protein levels were observed in eIF5A mutants (Figs. 5A–D and EV5A,B). We confirmed that the differences in increased Ssa1 levels under heat stress between wild type and eIF5A mutant cells were not due to a lower stability of full-length or truncated Ssa1 versions in eIF5A mutants (Fig. EV5C).

**Figure 5.**
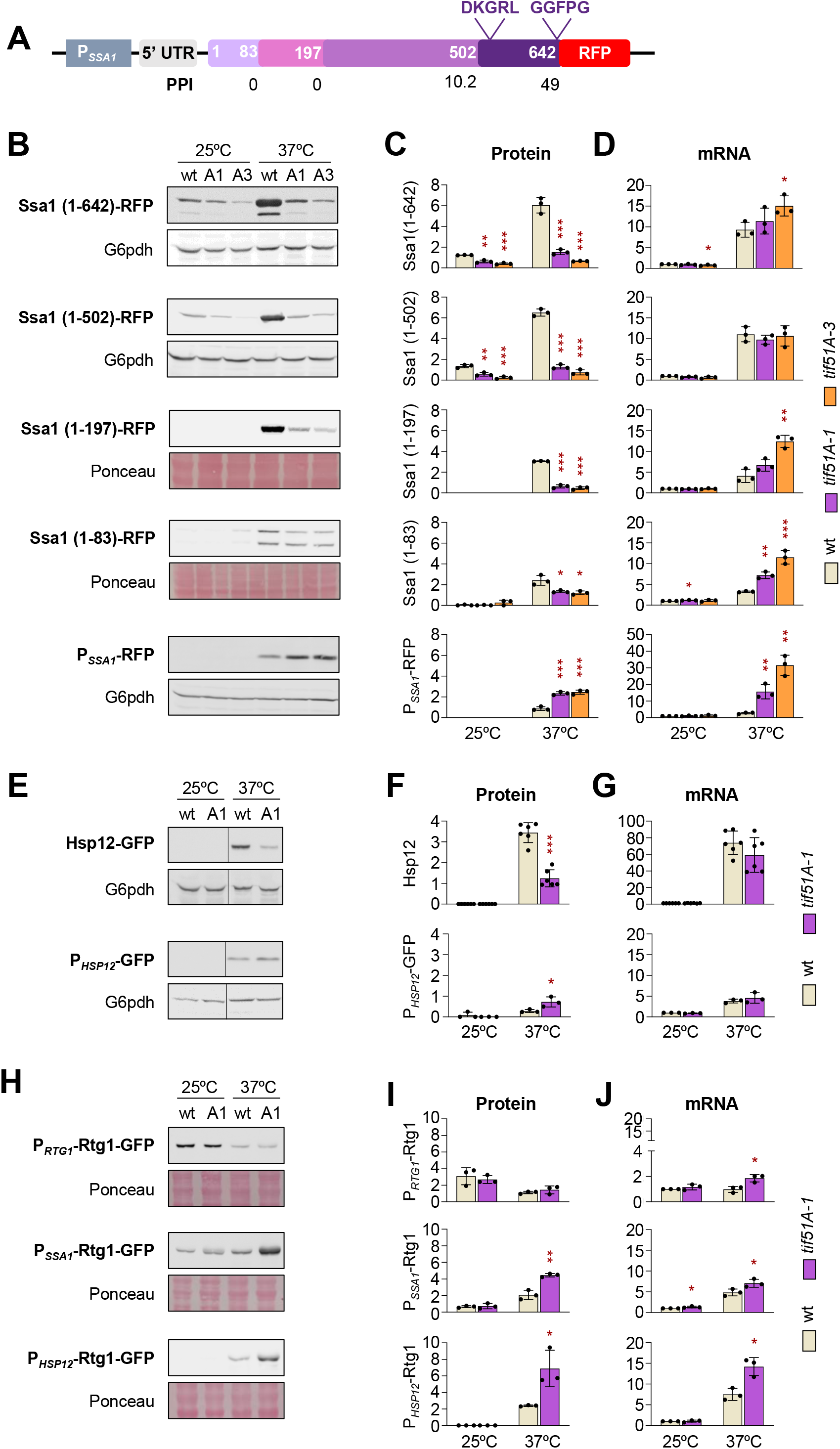
The chaperone coding sequence (CDS), independently of the eIF5A-peptide pausing motifs, dictates the translational eIF5A requirement during heat stress. **(A)** Schematic representation of *SSA1* locus encoding full Ssa1 protein and truncated protein versions, C-terminal fused to RFP. All constructs were expressed under the control of the endogenous *SSA1* promoter (P*_SSA1_*). Numbers inside boxes indicate the amino acid in the Ssa1 protein sequence. The positions of putative eIF5A-dependent motifs (DKGRL and GGFPG) and the calculated protein pausing index (PPI) for each construct are also indicated. **(B-D)** Wild type (wt), *tif51A-1* (A1), and *tif51A-3* (A3) strains expressing the indicated reporter constructs were grown in YPD at 25°C to the early exponential phase and then shifted to 37°C for 4 h. Representative western blots **(B)**, protein levels quantification normalized to loading controls **(C)**, and relative mRNA levels **(D)** of Ssa1-RFP full-length and truncated Ssa1 versions, and of a P*_SSA1_*-RFP reporter expressed in wt, *tif51A-1*, and *tif51A-3* strains. Proteins were detected using an anti-RFP antibody. G6pdh or Ponceau S staining was used as a loading control. **(E-G)** Representative western blots **(E)**, protein quantification normalized to G6pdh **(F)**, and relative mRNA levels **(G)** of Hsp12-GFP and a P*_HSP12_*-GFP reporter in wt and *tif51A-1* strains. Western blots were developed using an anti-GFP antibody, with G6pdh as a loading control. In **(E)**, vertical dividing lines indicate spliced non-adjacent lanes from the same membrane. **(H-J)** Representative western blots **(H)**, protein quantification normalized to Ponceau S **(I)**, and relative mRNA levels **(J)** of Rtg1-GFP expressed under the control of its endogenous promoter (P*_RTG1_*), the *SSA1* promoter (P*_SSA1_*), or the *HSP12* promoter (P*_HSP12_*) in wt and *tif51A-1* strains. Proteins were detected using an anti-GFP antibody, with Ponceau S staining as a loading control. **Data information:** For all RT-qPCR experiments (D, G, J), mRNA levels were normalized to endogenous *ACT1* mRNA and are expressed relative to the wt strain at 25°C. Quantitative data in (C, D, F, G, I, J) represent the mean ± SD from independent biological experiments (n = 3 to 6). Individual data points are plotted as dots. Representative western blot images are shown in (B, E, H). Statistical significance was assessed using a two-tailed unpaired Student’s t-test (\**P* < 0.05, \*\**P* < 0.01, \*\*\**P* < 0.001). Asterisks indicate significant differences between the eIF5A mutant and the wt strain under the same temperature condition. Exact *P* values for all comparisons are provided in the associated Source Data file.

Altogether, these results suggested that, first, the defect in the Ssa1 protein synthesis in the eIF5A mutant is mostly independent of the presence of eIF5A-peptide motifs in the Ssa1 protein sequence. Second, the defect is not dependent of the 5’ untranslated region (5’UTR) of *SSA1* mRNA, but is contingent to the CDS of *SSA1* mRNA. These conclusions were further supported by two additional experiments. In the first one, we obtained a genomic construct with *HSP12* promoter fused to GFP, thus with no *HSP12* CDS region. Again, we observed that the defect in Hsp12-GFP synthesis was lost in the eIF5A mutant when only GFP was translated under the control of the *HSP12* 5’UTR (Fig. 5E,F,G). In a second experiment, we expressed the CDS of the *RTG1* gene under the control of the *SSA1* promoter. The idea behind was to check heat-stress mediated synthesis of a protein such as Rtg1, which is not induced by heat stress in native conditions (Fig. 5H,I,J) and presumably has no eIF5A dependency (PPI index of 12.93). Therefore, this new strain rendered a heat stress inducible mRNA containing the *SSA1* 5’UTR and the *RTG1* CDS fused to GFP. First, we observed that Rtg1 protein levels increased very little in both wild type and eIF5A mutant strain under mild heat stress, pointing again to the *SSA1* CDS, and not to its 5’UTR, as mainly responsible for the huge synthesis of the Ssa1 protein seen under heat stress. Second, no less but even slightly more Rtg1 protein was synthesized in the eIF5A mutant with respect to the wild type from this hybrid mRNA containing the *SSA1* 5’UTR (Fig. 5H,I,J). To be able to generalize these results, we made a similar genomic fusion of *RTG1* CDS but in *HSP12* locus. Therefore, Rtg1 protein expression was induced by the *HSP12* promoter and the resulting mRNA contained the *HSP12* 5’UTR. Again, we observed not lower but higher protein levels of Rtg1 in eIF5A mutant under mild heat stress (Fig. 5H,I,J). These results pointed again to the CDS of the *SSA1* or *HSP12* mRNAs as the *cis* elements causing both, high protein synthesis and eIF5A-dependeny for translation.

In summary, our results support the idea that defects in the synthesis of specific heat stress proteins in eIF5A-depleted cells during mild heat shock are not caused by the presence of codons in the mRNA for the eIF5A-dependent amino acid motifs, nor by problems in translation initiation due to the specific mRNA 5’ UTR. Rather, they are linked to a different feature of these particular coding regions.

### Under mild heat-stress eIF5A preferentially binds and impacts the translation of heat-stress mRNAs with heavy translation via high codon optimality

We conducted further research into the characteristics that differentiated between the CDS of mRNAs that were dependent and independent of eIF5A for translation under mild heat shock. A striking difference was the percentage of optimal codons: eIF5A-dependent mRNAs under mild heat stress showed high codon optimality, such as *SSA1* and *HSP12* (87.1% and 85.5%, respectively); in contrast, eIF5A-independent mRNAs such as *SSA3*, *SSA4*, *HXK1*, *CCT1* and the control gene *RTG1* showed lower codon optimality (50.8%, 53.7%, 71.6%, 59.0% and 48.3%, respectively) (see Appendix Fig. S2A). It is noteworthy that high codon optimality appears to be an important feature of mRNA that promotes heavy translation and high synthesis of the corresponding chaperone proteins. However, codon optimality alone does not appear to be sufficient for high protein synthesis, as the RFP protein, whose CDS is also optimal (92.8%), was poorly synthesized under the control of the *SSA1* promoter (see Fig. 5B–E). We reasoned that, if high codon optimality was driving heavy translation of *SSA1* and *HSP12* mRNAs and thus, eIF5A dependency under mild heat stress, then reducing the optimality would eliminate eIF5A dependency in synthesizing the corresponding proteins. To test this, we created *SSA1*-GFP and *HSP12*-GFP genomic fusions with the native nucleotide sequences of the CDS (Ssa1(87opt) and Hsp12(86opt)), as well as with sub-optimized CDSs in which the *SSA1* optimality was reduced to 54% (Ssa1(54opt)) and that of *HSP12* to 45% (Hsp12(45opt)). The changes were made to synonymous codons, thus maintaining the encoded amino acid sequence, and were distributed throughout the CDS. Furthermore, in the case of *SSA1*, where the amino acid was conserved between the Ssa1 and Ssa4 proteins but the corresponding codons differed, we replaced the optimal *SSA1* codons for the non-optimal *SSA4* codons (see Appendix Fig. S3 and details in Methods). Additionally, we confirmed that the expression of GFP alone at the *SSA1* loci, where the full *SSA1* CDS has been deleted, was unaffected by eIF5A depletion (see Appendix Fig. S2B,C,D), similar to the results obtained with RFP (Fig. 5B,C,D). Analysis of the Ssa1(54opt) synthesis under mild heat stress revealed lower levels of the protein and the corresponding mRNA than those of the native Ssa1(87opt) in wild type cells. These results again indicated the strong link between the mRNA codon optimality and its stability and translation efficiency. Under the same stress conditions, we observed a lower synthesis of Ssa1(54opt) in eIF5A mutant cells than in wild type cells, although the difference was smaller than that observed with Ssa1(87opt) (Fig. 6A,B,C). These results do not conclusively reject our hypothesis that eIF5A dependency in translation is linked to codon optimality because both the Ssa1(87opt) and Ssa1(54opt) proteins contain peptide motifs that are putative eIF5A targets. To eliminate this variable, we next studied the translation of the low-optimal version of *HSP12* mRNA, since this Hsp12 protein has a PPI index of 0 and its amino acid sequence should not confer eIF5A dependency. As expected, we observed a lower increase in the levels of Hsp12(45opt) protein and mRNA than in those corresponding to Hsp12(87opt) in wild type cells under mild heat shock. Remarkably, no difference in the levels of synthesized Hsp12(45opt) was observed between wild type and eIF5A mutant cells under mild heat-shock (Fig. 6D,E,F). Overall, the results strongly suggest that eIF5A is necessary for the high efficient synthesis of stress proteins under mild heat stress driven, among other factors, by high codon optimality.

**Figure 6.**
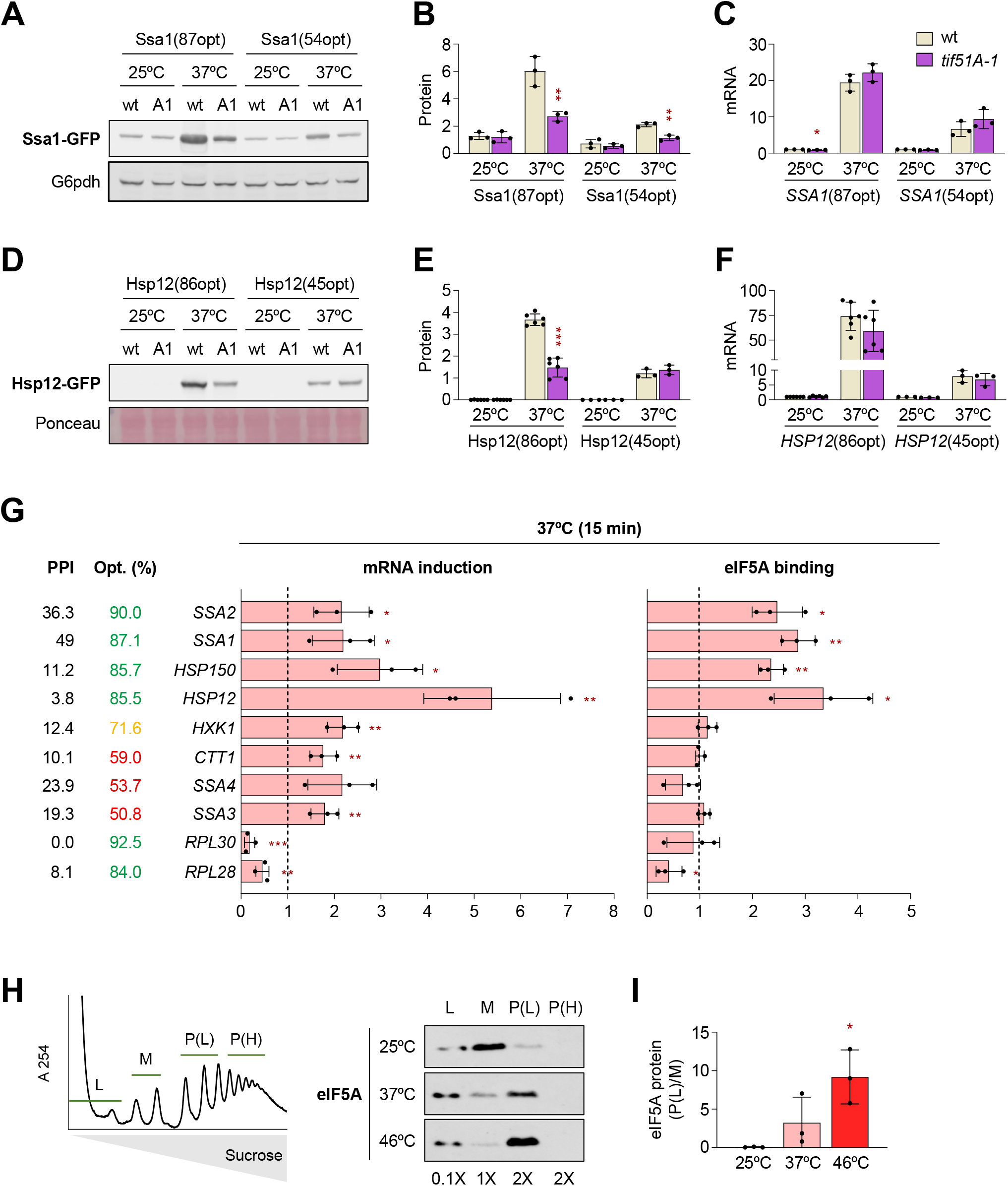
Under mild heat stress, eIF5A binds specifically to and is required for the optimal codon-driven efficient translation of chaperone mRNAs. **(A–F) Translation of native and codon-deoptimized chaperones under heat stress.** Wild type (wt) and *tif51A-1* strains expressing endogenously tagged native (Ssa1(87opt)-GFP, Hsp12(86opt)-GFP) or codon-deoptimized (Ssa1(54opt)-GFP, Hsp12(45opt)-GFP) constructs for Ssa1 **(A–C)** and Hsp12 **(D–F)** were grown in YPD at 25°C until the exponential phase and then shifted to 37°C for 4 h. **(A, D)** Representative western blots evaluating protein levels using an anti-GFP antibody. G6pdh or Ponceau S staining was used as loading controls. **(B, E)** Quantification of protein levels from the western blots in **(A)** and **(D)**, normalized to the respective loading control. **(C, F)** Relative mRNA levels determined by RT-qPCR, normalized to endogenous *ACT1* mRNA and expressed relative to the wt strain at 25°C for each respective construct. **(G)** eIF5A binds selectively to heat stress induced codon-optimal mRNAs during mild heat stress. RNA immunoprecipitation (RIP) was performed following formaldehyde crosslinking in wt cells grown in YPD at 25°C and shifted to 37°C for 15 min, using a specific anti-eIF5A antibody. Bar graphs show the fold change relative to 25°C for both total mRNA induction and eIF5A binding. The vertical dashed line at 1 marks the baseline level. The table on the left displays the eIF5A protein pausing index (PPI) of each encoded protein and the codon optimality (Opt. %) of each mRNA. The color indicates high (green), intermediate (yellow) or low (red) optimality. **(H)** eIF5A associates with polysomal fractions under heat stress. The wt strain was grown in SC medium to the exponential phase at 25°C and transferred to 37°C for 30 min (labeled as 37°C) or 37°C for 4 h followed by 30 min at 46°C (labeled as 46°C). Polysome profiling was performed and fractions were pooled into free RNA (L), monosomes (M), light polysomes (P(L)) and heavy polysomes (P(H)). eIF5A protein levels in each pool were analyzed by western blotting using an anti-eIF5A antibody. The relative volume of each pooled fractions loaded into the gel is indicated in the lower part. **(I)** Quantification of the eIF5A polysome/monosome ratio from the western blot shown in (H). **Data information:** In (B, C, E–G, I), data are presented as the mean ± SD from independent biological experiments (n = 3, except for E and F where n = 3 to 6). Individual data points are plotted as dots. In (B-F), statistical significance was assessed using a two-tailed unpaired Student’s t-test (\**P* < 0.05, \*\**P* < 0.01, \*\*\**P* < 0.001). In (G,I), statistical significance was assessed using a two-tailed paired Student’s t-test (\**P* < 0.05, \*\**P* < 0.01, \*\*\**P* < 0.001). In (G), statistical significance was determined by comparing the fold change of each specific gene to the fold change of *ACT1* under the same temperature conditions. In (I), asterisks indicate significant differences in the polysome/monosome ratio at 37°C or 46°C compared to 25°C. Exact *P* values for all comparisons are provided in the associated Source Data file.

Next, we wanted to investigate whether the role of eIF5A in the efficient translation of chaperone mRNAs under heat stress is direct. For this purpose, we first performed RNA immunoprecipitation (RIP) experiments to check for binding of the eIF5A protein to mRNAs under heat stress. Following a brief incubation of wild type yeast cells at 37°C, we detected an increase in the mRNA levels of heat stress responsive genes, including the *SSA1-4*, *HSP12*, *HSP150*, *HXK1*, and *CTT1* mRNAs. Conversely, the levels of *RPL30* and *RPL28* mRNAs decreased as expected (Fig. 6G). Notably, we detected an enrichment of eIF5A association with induced mRNAs that exhibited high codon optimality. This was particularly evident in the case of the Hsp70 *SSA1-4* mRNAs, which all increased to similar level under these heat stress conditions, however, only the highly optimal and heavy translated *SSA1* and *SSA2* mRNAs, but not the lower optimal and poorly translated *SSA3* and *SSA4* mRNAs, were bound by eIF5A (Fig. 6G). Similarly, eIF5A bound to the optimal *HSP12* and *HSP150* mRNAs, but not to lower optimal *HXK1* and *CTT1* mRNAs. This correlated with the requirement for eIF5A in the synthesis of Hsp12 and Hsp150, but not for Hxk1 and Ctt1 synthesis, under mild heat stress (Figs. 4 and EV3).

Under non-stress conditions, eIF5A acts as a translation elongation factor for specific mRNAs (Pelechano & Alepuz, 2017; Schuller *et al*, 2017). Due to its impact on only a small proportion of total mRNA, it is barely detectable in polysomal fractions (Zanelli *et al*, 2006). However, as we have demonstrated, large quantities of chaperone proteins are synthesized in a partially or totally eIF5A-dependent manner under mild and severe heat stress, respectively. Therefore, we wondered whether eIF5A would become more associated with translationally active fractions under heat stress. Analysis of the protein content of the polysome fractions showed that this was indeed the case, with increased eIF5A association with polysomes observed at 37°C and 46°C (Fig. 6H,I). This result was consistent with our previous observation that eIF5A remains mainly cytosolic, and not accumulated in condensates, during heat stress (Fig. 2G).

In sum, these results suggest a direct role for eIF5A in the translation of heat stress mRNAs. Moreover, the results suggest that, under mild heat stress, eIF5A specifically binds to heat stress mRNAs under heavy translation driven by high codon optimality, where it promotes the efficient synthesis of the corresponding proteins.

### Defects in chaperone synthesis under heat shock in eIF5A mutants result in a transcriptional over-response via Hsf1 and impaired proteostasis

Our results have shown that the translation, and therefore the synthesis, of heat stress inducible chaperones such as the Hsp70 Ssa1 and Ssa2 depends on eIF5A. During the heat shock response, the inhibitory interaction between Hsp70 and Hsf1 is lost as Hsp70 is titrated away by stress specific chaperone clients. This enables Hsf1 to bind to the heat shock elements (HSEs) at the chaperone promoters. Induction of Hsp70 expression results in an excess of Hsp70 protein, which can then bind back and inhibit Hsf1, creating a negative feedback loop that shuts down the transcriptional response (Dea & Pincus, 2024; Krakowiak *et al*, 2018). As eIF5A is necessary for the synthesis of Ssa1 and Ssa2 under heat stress (Fig. 4A,B,C), and as we also observed higher and more sustained levels of chaperone mRNAs under heat stress in eIF5A mutants (Fig. 2A), we hypothesize that this Hsp70-dependent negative feedback loop over Hsf1 could be altered in eIF5A mutants (Fig. 7A). To test this, we first obtained yeast wild type and eIF5A mutant strains with a genome-integrated yellow fluorescent protein (YFP) under the control of a promoter containing four repeats of the heat-shock element (4xHSE-YFP) (Krakowiak *et al*, 2018; Zheng *et al*, 2016), and studied the induction kinetics under heat stress following the authors induction conditions. We observed a significant increase in YFP fluorescence that peaked at 1 h of heat stress and then slowly decreased in the wild type. In the eIF5A mutant, the increase in YFP was slower but reached similar levels to those in the wild type cells after 90 min. Later on, the YFP fluorescence continued increasing in the mutant cells, with levels of YFP more than doubling compared to the wild type after 5 hours of heat stress (Fig. 7B). However, Hsf1 protein levels and kinetics were similar in both strains (Fig. 7C). As these results supported our model (Fig. 7A), we next examined Hsf1 binding to the *SSA1* promoter, which contains two HSE, during heat stress using chromatin immunoprecipitation (ChIP). In wild type cells, we observed an increase in Hsf1 binding that reached its maximum at 10-20 minutes of heat stress, which was lost after 30 minutes (Fig. 7D). This modest increase in Hsf1 binding to the *SSA1* promoter was similar to that previously described (Pincus *et al*, 2018). However, in the eIF5A mutant cells, Hsf1 showed higher and prolonged binding to the *SSA1* promoter that remained significantly high after 30 and 45 minutes of heat stress (Fig. 7D). Overall, these results support the model in which the absence of eIF5A reduces Ssa1 chaperone synthesis, consequently affecting the transcriptional induction and shutdown of heat-stress-induced genes by altering the inhibitory effect of Ssa1 on Hsf1 (see Fig. 7A).

**Figure 7.**
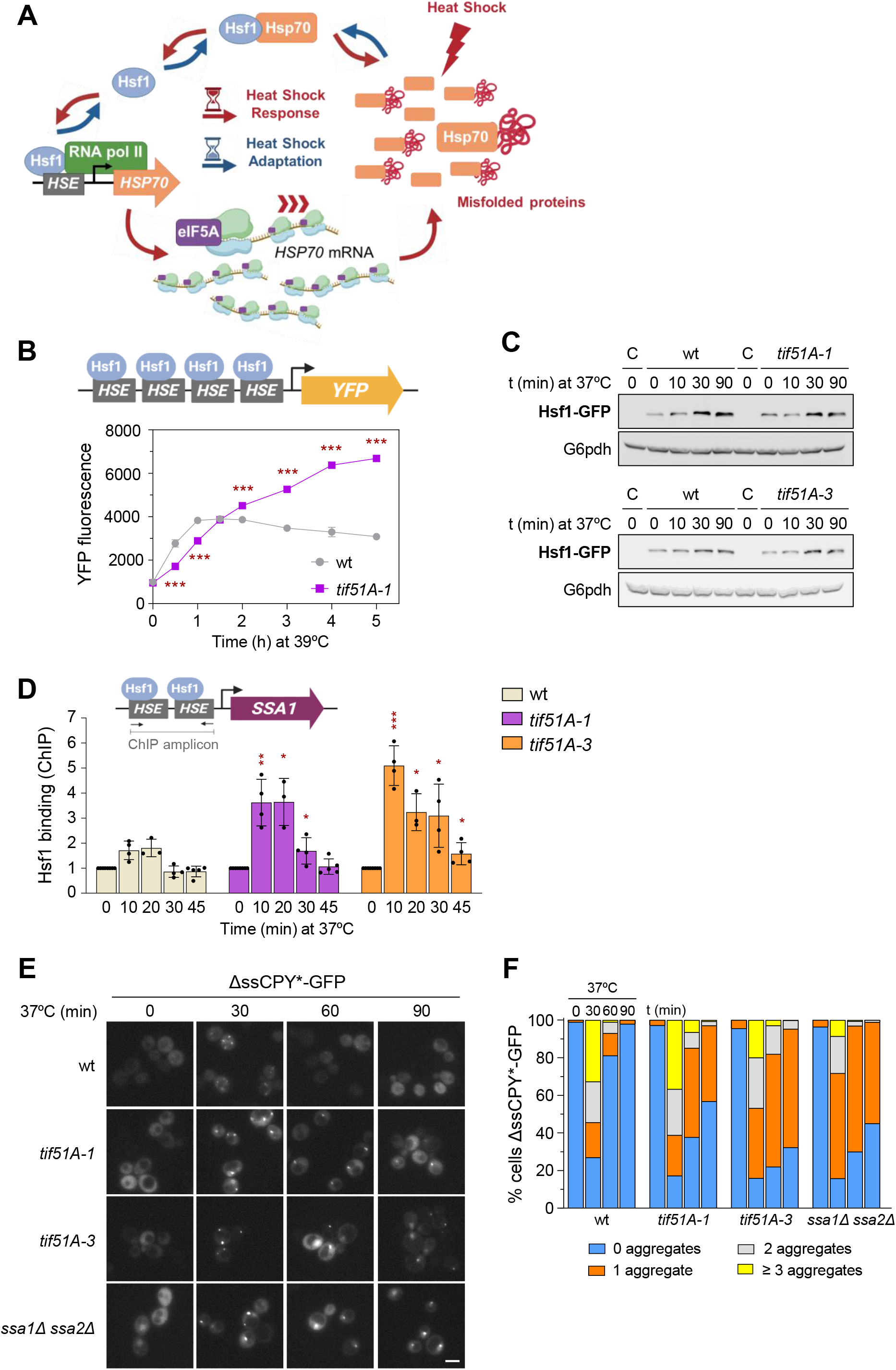
Impaired eIF5A function results in Hsf1 over-activation of transcription and proteostasis disruption under heat stress. **(A)** Model for eIF5A in the Hsp70-mediated inhibitory feed-back loop over Hsf1 during the adaptation to heat stress. **(B)** Effect of eIF5A dysfunction in the induction of 4xHSE-YFP reporter by Hsf1. Cultures were grown until early exponential phase in YPD medium at 25°C and then incubated at 39°C for the indicated times. To allow for complete YFP fluorophore maturation, translation was halted with cycloheximide and samples were incubated for 2 h at 25°C prior to analysis. It is shown the quantification of the fluorescence intensity determined by flow cytometry in wild type (wt) and *tif51A-1* cells. **(C)** Representative Western blot analysis evaluating Hsf1-GFP protein levels in wt, *tif51A-1,* and *tif51A-3* strains. Cultures were grown until exponential phase in YPD at 25°C and then shifted to 37°C for the indicated times. Untagged strains for each corresponding genetic background harvested at time 0 were included as negative controls (C). Specific antibodies against GFP and G6pdh were employed, with G6pdh used as a loading control. **(D)** Chromatin immunoprecipitation (ChIP) assay evaluating the recruitment of Hsf1-GFP to the *SSA1* promoter region. Exponentially growing cells in YPD at 25°C were subjected to heat stress at 37°C for the specified times. Hsf1-GFP ChIP was performed in wild type (wt) and *tif51A-1*, and *tif51A-3* mutant strains using an anti-GFP antibody. The top scheme shows the *SSA1* promoter region with HSE elements and the location of the ChIP-qPCR amplicon. **(E,F)** Wild type (wt), eIF5A mutants (*tif51A-1*, *tif51A-3*), and chaperone deletion strains *ssa1Δ ssa2Δ, hsp12Δ, hsp26Δ*, *hsp42Δ*, *hsp104Δ*, containing a plasmid with the ΔssCPY*-GFP protein aggregation reporter were cultured in SC-Ura medium at 25°C until the early exponential phase and then incubated at 37°C for the indicated times. **(E)** Representative fluorescence microscopy images of wt, *tif51A-1*, *tif51A-3* and *ssa1Δ ssa2Δ* cells taken at 0, 30, 60 and 90 min after the temperature shift. Scale bar: 5 µm. **(F)** Quantification of ΔssCPY*-GFP aggregates. The stacked bar charts represent the percentage of cells containing 0, 1, 2 or ≥ 3 fluorescent foci at each time point for all indicated strains. **Data information:** In (B, D), data represent the mean ± SD from at least three independent biological experiments (n = 3 for B; n = 3 to 5 for D). Individual data points are plotted as dots. ChIP data in (D) represent relative enrichment normalized against total input DNA and expressed relative to the t = 0 value for each respective strain. Statistical significance in (B, D) was determined using a two-tailed unpaired Student’s t-test (*P < 0.05, **P < 0.01, ***P < 0.001). Asterisks indicate significant differences relative to the wt control under the same time condition. Exact P values for all comparisons are provided in the associated Source Data file. Images in (E) are representative of three independent biological experiments. Quantitative data in (F) represent the mean from three independent biological experiments (n ≥ 100 cells were scored per strain and condition).

The rapid and enormous increase in chaperones that occurs upon heat shock regulates cellular proteostasis, as chaperones prevent thermally damaged proteins from aggregating and resolve protein aggregates by unfolding, refolding or targeting proteins for degradation (Verghese *et al*, 2012). As chaperone synthesis is affected in eIF5A mutants under heat stress, we examined the formation and resolution of protein aggregates at elevated temperatures. To do this, we used the protein aggregation reporter ΔssCPY*-GFP, which is a mutant derivative of the yeast endoplasmic reticulum (ER) carboxypeptidase (Medicherla *et al*, 2004). This derivative lacks the ER-directing signal and therefore remains in the cytoplasm, where it is prone to unfolding and aggregation. Furthermore, it has been demonstrated that the Hsp70 chaperone Ssa1 recognizes this misfolded protein and conducts it towards proteasome degradation (Park *et al*, 2007). We expressed ΔssCPY*-GFP in both wild type and eIF5A mutant yeast cells and subjected them to heat stress at 37°C. In wild type cells, we observed transient ΔssCPY*-GFP aggregates that were visible at 30 minutes and had disappeared after 60 minutes of heat stress. However, more intense aggregates were visualized in the eIF5A mutant cells at 30 minutes of stress, and were still present after 90 minutes (Fig. 7E,F). We also studied the dynamics of the ΔssCPY*-GFP aggregates in cells with mutations in chaperones: *ssa1Δ ssa2Δ*, *hsp12Δ*, *hsp26Δ*, *hsp42Δ* and *hsp104Δ*. As previously described, among the chaperone mutants tested, a lack of the partially redundant Ssa1 and Ssa2 chaperones resulted in the most severe defects in resolving the ΔssCPY*-GFP aggregates. Notably, the pattern and dynamics of the ΔssCPY*-GFP aggregates were highly similar between the *ssa1Δ ssa2Δ* mutant and the eIF5A mutants (Fig. 7E,F and Appendix Fig. S4). These results support the role of eIF5A in promoting the translation, and therefore the synthesis, of the Hsp70 Ssa1 and Ssa2 chaperones. Furthermore, the results imply that eIF5A regulates proteostasis during heat stress by facilitating the efficient production of chaperones.

## DISCUSSION

In this study, we discovered that the translation elongation factor eIF5A plays a pivotal role in maintaining ribosome elongation activity under heat stress, thereby facilitating the abundant synthesis of chaperones that regulate cellular proteostasis. While it was previously thought that global translation shutdown was essential for sustaining a high translation rate of stress-response genes, our results suggest that eIF5A, rather than ribosomes, could be the limiting factor. Thus, the disassembly of ribosomes from house-keeping mRNAs and their targeting to PB and SG may release eIF5A, which, together with newly eIF5A synthesized protein, could then associate with heat stress induced mRNAs and promote their heavy translation under heat stress.

Loss of functional eIF5A has a severe impact at high temperatures, where a lack of ribosome movement is observed across all mRNAs. However, at temperatures that induce moderate stress, the absence of eIF5A affects ribosome mobility depending on the particular features of each stress-response mRNA. This negative effect is more pronounced in heavily translated mRNAs with high codon optimality, such as those encoding most heat shock proteins (Hsp). In these mRNAs, heat stress leads to increased translation efficiency due to easier ribosome engagement with these transcripts (Carneiro *et al*, 2025). Our results demonstrate greater binding of eIF5A to these mRNAs; however, we cannot ascertain whether this association and specificity originates directly from the specific codons or the resulting high ribosome density. Interestingly, slow ribosome decoding at the A-site, occurring in regions with non-optimal codons, generates stalled ribosomes recognized by the Ccr4-Not complex, which mediates mRNA decay (Buschauer *et al*, 2020). In these stalled ribosomes, the Not5 subunit of the complex binds to the empty E-site of the ribosome when the A-site is also empty (-/PP/-). It has therefore been proposed that Not5 and eIF5A compete for the empty E-site in an A-site-dependent manner. Not5 binds to stalled ribosomes in the -/PP/-conformation, whereas eIF5A binds to those in the AA/PP/-conformation (Buschauer *et al*, 2020). It remains unknown whether heavy translation under heat stress, particularly in regions of high codon optimality, leads to the formation of transiently stalled AA/PP/-ribosomes. This could occur if empty tRNA dissociates from the E-site faster than peptide bond formation occurs. The specific role of eIF5A in promoting ribosome elongation at optimal codons is further supported by ribosome profiling studies in yeast, in which an inverse correlation was observed between ribosome occupancy at optimal and non-optimal codons in eIF5A and Not5 mutants. Thus, optimal codons were enriched among those with increased ribosome occupancy under eIF5A depletion (Allen *et al*, 2021).

In sum, our results demonstrate a new function of eIF5A in translation elongation that is independent of the previously described in promoting peptide bond formation between specific amino acids (Pelechano & Alepuz, 2017; Schuller *et al*, 2017). Conversely, dependency on eIF5A appears to be associated with highly efficient ribosome elongation of mRNA under heat stress. At mild heat stress (37°C), eIF5A-dependent fast ribosome elongation is primarily associated with codon optimality in mRNA. However, at more severe heat stress (46°C), the increased temperature might also accelerate ribosomal speed as translation efficiency increases (Carneiro *et al*, 2025), resulting in a broader eIF5A dependency. The fact that the bacterial homologue of eIF5A, EF-P, is only necessary in *Escherichia coli* under conditions of high translational demand supports this new function. Thus, *E. coli* lacking EF-P grows well at 25°C but exhibits reduced growth at a high temperature (37°C) (Tollerson *et al*, 2018). Whether this new eIF5A function is strictly linked to ribosome elongation at high temperatures or is common to other contexts of heavy translation, independent of temperature, remains to be determined.

The new role of eIF5A in supporting efficient translation, together with its previously described function in facilitating the translation of specific, hard-to-translate peptide motifs, are both advantageous for protein synthesis. However, these roles contrast with those of eIF5A in ribosome quality pathways, which repress protein production in response to ribosome stalls. A reconciled model would position eIF5A as a ribosome controller that determines whether stalled ribosomes progress in translation or get disassembled. Along these lines, recent work in *C. elegans* and yeast has proposed that eIF5A may act as a sensor that distinguishes between different types of ribosome stalls, determining the fate of ribosomes and mRNA in quality control pathways (Wohlenberg *et al*, 2026). This model is supported by data from mammalian cells which suggest that eIF5A promotes efficient translation by preventing the activation of the RQC pathway, which disassembles stalled ribosomes (Schneider-Poetsch *et al*, 2025). However, eIF5A also participates in RQC once the ribosome has been disassembled (Tesina *et al*, 2023). As proposed by Wohlenberg *et al*. (Wohlenberg *et al*, 2026), the role of eIF5A as a ribosome stall sensor would explain its ubiquity in different disome structures (Pochopien *et al*, 2021; Schmidt *et al*, 2016). Moreover, eIF5A is also found at the E-site of ribosomes that have paused but reinitiate translation (Zeng *et al*, 2021). Our results open up the possibility that eIF5A acts as a sensor and resolution factor for ribosome collisions in the context of heavy and rapid translation (Juszkiewicz *et al*, 2018; Park & Subramaniam, 2019). Interestingly, ribosome collisions do not always impede fast and efficient translation, but rather promote it. *In vitro* translation studies have suggested that a transient collision between a tailing and a stalled ribosome could help the latter move forward, thus reducing ribosome recycling and increasing processivity (Madern *et al*, 2025). All these results bolster the idea that eIF5A plays a role in this ribosome cooperation by detecting collisions and promoting the reactivation of stalled ribosomes.

Our results show that the Ssa1 Hsp70, one of the most inducible HSPs in terms of promptness and robustness (Cuesta *et al*, 2000; Theodorakis & Morimoto, 1987), is the heat stress protein whose synthesis is most affected in eIF5A mutants. Interestingly, the mammalian eEF1A translation elongation factor eEF1A, whose canonic role is to deliver aminoacylated tRNAs to the A-site of the ribosome during translation elongation, has been shown to positively coordinate *HSP70* expression by binding to *HSP70* mRNA during transcription (Vera *et al*, 2014). This result suggests that the abundant synthesis of Hsp70 requires a coordinated regulation of its expression, from transcription to translation. It cannot be ruled out that eF5A binding to *SSA1* mRNA occurs in the nucleus, as we have previously demonstrated eIF5A binding to specific genes in the nucleus in an RNA-dependent manner (Barba-Aliaga *et al*, 2026). However, further research is needed to fully clarify the specific interaction dynamics between eIF5A and chaperone mRNAs.

In light of the role of eIF5A as a factor required for rapid and efficient translation and for the synthesis of chaperones under heat stress, the connections between eIF5A and stress- and disease-related contexts can be reconsidered. In plants, defects in eIF5A result in thermosensitivity, whereas overexpression increases heat tolerance. Furthermore, increasing active eIF5A provides protection against other abiotic stresses (Ma *et al*, 2010; Wang *et al*, 2012; Xu *et al*, 2011). As rising global temperatures are a major constraint limiting plant growth and food production (Janni *et al*, 2024; Tran *et al*, 2025), it would be relevant to investigate whether the link between eIF5A activity and the abundant synthesis of chaperones is conserved in plants.

Neurological disorders are often caused by the misfolding, aggregation, and accumulation of specific disease proteins in the brain and central nervous system. Thus, disturbing the key role of chaperones in maintaining cellular proteostasis can lead to proteinophaties (Sinnige *et al*, 2020; Sood *et al*, 2025). In recent years, the role of eIF5A in neurological disorders has emerged. Heterozygous deleterious variants of eIF5A have been shown to cause the Faundes-Banka syndrome, characterized by developmental delay, microcephaly, micrognathia and dysmorphism (Faundes *et al*, 2021). Furthermore, biallelic deleterious variants of the eIF5A hypusination enzymes DHS and DOHH are associated with a rare neurodevelopmental disorder, causing global developmental delay and seizures (Beltrán-Corbellini *et al*, 2025; Ganapathi *et al*, 2019; Ziegler *et al*, 2022). eIF5A has also been linked to the pathological accumulation and aggregation of tau and TDP-43 proteins, which is a characteristic of the brains of patients with Alzheimer disease (Smeltzer *et al*, 2021). A recent study of amyotrophic lateral sclerosis (ALS)-associated mutations in the RNA-binding protein fused in sarcoma (FUS), showed that eIF5A controls local axon translation and *in vivo* treatment with spermidine mitigates defects in a *Drosophila* FUS-ALS model (Piol *et al*, 2025). In a mouse model of Huntington’s disease, it was demonstrated that the mutant huntingtin (Htt) protein depletes eIF5A in affected brains, causing increased ribosome stalling and impairing stress responses (Aviner *et al*, 2024). Moreover, aging is associated with neurological diseases and loss of proteostasis (López-Otín *et al*, 2023), and levels of eIF5A decrease in aged cells (Liang *et al*, 2021). It is therefore reasonable to speculate that eIF5A dysfunction may lead to deficient chaperone synthesis and loss of cellular proteostasis in a variety of circumstances. Future work will probe whether manipulating eIF5A activity could improve cellular proteostasis in disease contexts and thus mitigate proteinopathies.

## METHODS

### Yeast strains, plasmids and growth conditions

All *S. cerevisiae* strains and plasmids used in this study are listed in Appendix Tables S1 and S2, respectively. Cells were cultured in liquid YPD (1% yeast extract, 2% peptone, 2% glucose), YP (1% yeast extract, 2% peptone), or synthetic complete (SC) drop-out medium (0.7% YNB, Drop-Out mixture [Kaiser, Formedium], and 2% glucose or galactose). Solid media were prepared by adding 2% agar. For all experimental assays, cells were grown to the exponential phase at 25°C or 30°C. To deplete eIF5A, temperature-sensitive strains grown at 25°C were shifted to restrictive temperatures (37°C, 39°C, 41°C, or 46°C) as indicated (Li *et al*, 2014). Where specified, media were supplemented with 4 mM spermidine (Merck Life Science; S0266). For survival assays using the *tif51AΔ tif51BΔ* strain expressing murine eIF5A (p*GAL*-m eIF5A), cells were grown in SGal-Trp at 25°C to the exponential phase. Then, to deplete eIF5A, cultures were shifted to glucose medium (SC-Trp) for 6 h at 25°C prior to a 46°C heat shock. To repress *TIF51A* expression in the *tetO₇-TIF51A* strain, liquid cultures were grown in the dark with 15 μg/mL doxycycline until the early exponential phase; doxycycline was then refreshed prior to a temperature shift (37°C for 4 h, followed by 46°C for 30 min). For solid media assays, 0.1 μg/mL doxycycline was added. See Appendix Supplementary Methods for more details.

### Strain engineering and plasmid transformation

For genomic tagging of full-length open reading frames (ORFs) with HA, GFP, or RFP and complete ORF replacements, a PCR-based homologous recombination approach was employed using the plasmids pFA6a-3HA-HIS3MX, pFA6a-GFP-HIS3MX (Longtine *et al*, 1998), and pYM43-Redstar2-clonNAT (Euroscarf; P30255) as templates, along with the primers listed in Appendix Table S3.

All genetic modifications and plasmid introductions were performed using the standard lithium acetate method (Gietz *et al*, 1992). Transformants were selected on appropriate auxotrophic media or media supplemented with 100 μg/mL nourseothricin/clonNAT (GoldBio; N-500-1). Integrations were verified by PCR, western blotting, and/or Sanger sequencing (Macrogen Inc.).

To construct yeast strains with codon-deoptimized chaperones, synthetic *SSA1* and *HSP12* ORFs were designed *in silico* according to the codon optimality classification described by (Presnyak *et al*, 2015). These fragments were synthesized by Integrated DNA Technologies (IDT) and fused to a GFP-tagging cassette via overlap extension PCR (Ho *et al*, 1989). prior to yeast transformation. For detailed information about strain construction, see Appendix Supplementary Methods.

### Growth and viability analyses

To evaluate growth on solid media, cells were grown in YPD or SC medium at 25°C to the early exponential phase. Serial 10-fold dilutions of normalized cultures were spotted onto agar plates and incubated at the indicated temperatures for 2–3 days. To determine cell viability, colony-forming unit (CFU) assays were performed. Briefly, treated cultures were collected at specified time points, serially diluted, and plated onto solid media. Following incubation at 25°C for 2–3 days, colonies were counted to determine survival relative to the initial time point (t = 0).

### Polyribosome profile analysis

Cell extractions and polysome gradients were performed as previously described (Garre *et al*, 2012; Ramos-Alonso *et al*, 2018) Briefly, cells grown to early exponential phase at 25°C were treated with 100 μg/mL CHX for 5 min to arrest translation, harvested, and mechanically disrupted with glass beads in lysis buffer. For translational run-off experiments, CHX was omitted during harvesting and lysis. Lysates were cleared by sequential centrifugation, supplemented with 5% glycerol, and stored at −80°C.

For polysome profiling, 8.5 A_260nm_ units of cleared lysate were loaded onto 10–50% sucrose gradients and separated by ultracentrifugation (35,000 rpm, 2 h 40 min, 4°C) using a Beckman SW41Ti rotor. Gradients were fractionated into 22 aliquots using a density gradient fractionation system (Teledyne ISCO) with continuous monitoring at 240 nm. Total RNA from pooled fractions was extracted using the SpeedTools Total RNA Extraction Kit (Biotools; 21211) with integrated rDNase treatment. Target mRNAs were quantified by RT-qPCR (primers in Appendix Table S3). The relative abundance of each mRNA was expressed as a percentage of the total transcript distributed across the fractions. All polysome profiling experiments were performed at least in biological triplicates and a representative profile is shown. For detailed buffer compositions and procedures, see Appendix Supplementary Methods.

### RT-qPCR analysis

To analyze mRNA levels, total RNA was isolated from yeast cells following the phenol-chloroform-based extraction method described in (Barba-Aliaga & Alepuz, 2022). DNAse treatment, reverse transcription and quantitative PCR (qPCR) reactions were performed as detailed previously (Garre *et al*, 2013). Amplification and fluorescence detection were carried out on a CFX96 Touch Real-Time PCR Detection System (Bio-Rad) using the Supreme qPCR Green Master Mix (NZYtech; MB22403). Endogenous *ACT1* levels were used for normalization, and at least three biological replicates were analyzed for each sample. Specific primers are listed in Appendix Table S3, and further methodological details can be found in the Appendix Supplementary Methods.

### Western blotting

Whole-cell protein extracts were prepared using an alkaline lysis method (Zuzuarregui *et al*, 2015). For protein extraction from sucrose gradient fractions (obtained during polysome profiling), the 22 collected fractions were pooled into four distinct groups: free RNA (L), 60S+80S (M), light polysomes (P(L)), and heavy polysomes (P(H)) groups, as indicated in the corresponding figures. Proteins from these fractions were precipitated with 10% trichloroacetic acid (TCA), washed with cold acetone, and resuspended in SDS lysis buffer. Proteins were resolved by SDS-PAGE on polyacrylamide gels (ranged from 8% to 15%) and transferred to 0.45 μm nitrocellulose membranes (Cytiva; 10600001). After blocking with 5% skim milk, membranes were incubated overnight at 4 °C with primary antibodies: anti-GFP (Roche; 11814460001), anti-RFP (Thermo Fisher Scientific; A00682-40), anti-HA (Roche; 12013819001), anti-eIF5A (Abcam; ab32407), anti-Hog1-P (Cell Signaling Technology; 4511), anti-eIF2alpha-total (Santa Cruz; sc-11386), anti-eIF2alpha-P(Ser52) (Santa Cruz; sc-101670) or anti-G6pdh (Sigma-Aldrich; A9521). Bound antibodies were detected using HRP-conjugated anti-rabbit (Promega; W4011) or anti-mouse (Promega; W4021) secondary antibodies (complete list and working dilutions in Appendix Table S4).

Chemiluminescent signals were developed using Amersham ECL Prime (Cytiva; 10308449) and captured with an ImageQuant LAS 4000 system (Cytiva). Band intensities were quantified using ImageJ (Schindelin *et al*, 2012). To capture variation across all samples, the signal in each lane was normalized to the mean signal across all lanes in a single blot. Then, the resulting signal was normalized against the corresponding G6pdh signal or total protein (Ponceau staining), as indicated in each specific experiment. At least three biological replicates of each sample were analyzed.

For detailed buffer compositions and procedures, see Appendix Supplementary Methods.

### Protein stability determinations

To assess protein stability, translation was arrested by supplementing cultures with 100 μg/mL cycloheximide (CHX; Sigma-Aldrich; C7698). Samples were harvested over a 5-h period post-CHX addition, and protein levels were analyzed by western blotting. Protein quantification data were normalized to the initial time point (t = 0, set as 100%). The degradation rate constant (*k*_d_) for each protein was determined by fitting the normalized data to an exponential decay model (y = Ae^−*k*d t^).

### Fluorescence microscopy

For live-cell imaging, cells were grown to the early exponential phase and after treatment, cells were centrifuged and washed. Fluorescence images were acquired using a motorized Axio Imager Z1 microscope (Carl Zeiss, Oberkochen, Germany) equipped with a Plan-Apochromat 63x/1.4 oil-immersion objective, a 100-W mercury lamp, and an AxioCam MRm digital camera. Image acquisition was controlled via AxioVision v4.7 software (Carl Zeiss). Fluorophores were detected at the following excitation/emission wavelengths: GFP (475/509 nm), RFP (555/583 nm), and mCherry (587/610 nm). Exposure times were kept constant across all comparable samples.

Image processing and quantification were performed using ImageJ software (Schindelin *et al*, 2012). To quantify protein aggregation, cells were manually classified based on the number of visible inclusions per cell (0, 1, 2, or ≥ 3). A minimum of 100 cells per strain and condition were evaluated, and data was obtained from at least three independent biological replicates.

### Flow cytometry

Cells expressing the *4xHSE-YFP* reporter were grown to the early exponential phase at 25 °C and subjected to heat stress at 39 °C. At the indicated time points, translation was halted by adding 50 µg/mL cycloheximide, followed by a 2-h incubation at 25 °C to allow for complete YFP fluorophore maturation (Zheng *et al*, 2016). Cells were subsequently washed, fixed in 4% paraformaldehyde, and analyzed using a BD FACSVerse™ flow cytometer (50,000 events acquired per sample). All experiments were performed at least in biological triplicates. For detailed procedures see Appendix Supplementary Methods.

### RNA immunoprecipitation (RIP)

The RNA immunoprecipitation (RIP) protocol was adapted from (Gilbert & Svejstrup, 2006). Briefly, exponentially growing cells were cross-linked with 1% formaldehyde, quenched with 0.14 M glycine, and disrupted with glass beads in lysis buffer using a Precellys 24 homogenizer. Lysates were sonicated (Bioruptor, Diagenode), cleared by centrifugation, and treated with DNase I (Roche; 4716728001) and Ribonuclease Inhibitor (Promega; N2111) at 37 °C for 15 min. An input fraction was saved, and the remaining lysate was incubated with Dynabeads M-280 Sheep Anti-Rabbit IgG (Invitrogen; 11204D) pre-conjugated with an anti-eIF5A antibody (Selleck Chemicals; F2900). Following washes, RNA complexes were eluted, cross-linking was reversed at 65 °C for 1 h, and samples were treated with Proteinase K (Roche; RPROTKSOL-RO). Finally, RNA was purified using a Qiagen RNA Purification Kit (Qiagen; 74104).

The specific binding of eIF5A to target RNAs was analyzed by RT-qPCR using the specific primers listed in Appendix Table S3. The relative binding of eIF5A to each specific mRNA was quantified by normalizing the values obtained in the IP samples against their corresponding INPUT values. For more detailed procedures, see Appendix Supplementary Methods.

### Chromatin immunoprecipitation (ChIP)

Chromatin immunoprecipitation (ChIP) assays were performed as previously described (Li *et al*, 2014), with minor modifications (Barba-Aliaga & Alepuz, 2022). Briefly, exponentially growing cells were subjected to heat stress at 37°C, cross-linked with 1% formaldehyde, quenched with 0.14 M glycine, and disrupted with glass beads using a Precellys 24 homogenizer. After chromatin was sonicated (Bioruptor, Diagenode) and cleared by centrifugation, an input fraction was saved, and the remaining lysate was incubated overnight with Dynabeads Pan Mouse IgG (Invitrogen; 11041) pre-conjugated with anti-GFP (Roche; 11814460001)) for detecting Hsf1-GFP binding (antibodies and working dilutions in Appendix Table S4). Following sequential washes, protein-chromatin complexes were eluted, cross-linking was reversed at 65 °C, and DNA was purified using the NZYGelpure kit (NZYtech; MB01101).

Hsf1 binding to the *SSA1* gene was quantified by RT-qPCR using the specific primers designed to amplify the heat shock element (HSE) sequences located within the 5’ UTR regions (listed in Appendix Table S3). Data were normalized against total INPUT DNA value in the corresponding whole cell extract. At least three independent biological replicates were analyzed for each sample. More detailed procedures can be found in the Appendix Supplementary Methods.

### Condon optimality and PPI index determinations

A list of optimal and non-optimal codons in *S. cerevisiae* was procured from (Presnyak *et al*, 2015) and the CDSs of the specific yeast mRNAs were obtained from the Saccharomyces Genome Database (https://www.yeastgenome.org/). The occurrence of each optimal codon was counted and divide it by the total number of codons to calculate the percentages of optimal codons.

The translation dependency of each yeast gene on eIF5A was estimated by calculating the Protein Pause Index (PPI) as described in (Barba-Aliaga *et al*, 2026). Briefly, the number of the top 43 eIF5A-dependent tripeptide motifs present in the encoded protein amino acid sequence was determined. Then, the number of each motif was multiplied by the pause strength value observed when eIF5A is depleted, as revealed by 5PSeq analysis (Pelechano & Alepuz, 2017). Thirdly, the PPI for each gene was obtained as the sum of motifs x strength.

### Statistical analyses

Quantitative data are expressed as the mean ± standard deviation (SD). All graphs and statistical analyses were generated using GraphPad Prism (version 8.0.2; GraphPad Software, Boston, MA, USA). Statistical significance was determined using a two-tailed Student’s *t*-test. Specifically, an unpaired *t*-test was applied when comparing independent groups, whereas a paired *t*-test was utilized when comparing the same strain subjected to different experimental conditions or treatments. A *p*-value < 0.05 was considered statistically significant. Unless specified, all experiments were obtained from at least three independent biological replicates performed on different days. For microscopy analyses, at least 100 cells per strain and condition were evaluated, and data were pooled from at least three independent biological replicates.

## Data availability

This study includes no data deposited in external repositories

## Author Contributions

P.A. designed research; V.B., L.C., M.B-A., J.S. and S.P.-D. performed research; V.B., L.C., M.B-A., J.S., S.P.-D. and P.A. analyzed data; P.A. wrote the paper. All authors revised the paper.

## Acknowledgements

This work was supported by grants PID2020-120066RB-I00 and PID2023-152214NB-100 by Spanish Ministry of Sciences and Innovation MCIN/AEI/10.13039/501100011033 to PA. This research was also funded by Generalitat Valenciana (CIAICO/2022/237, INVEST/2022/334 and CIACO/2026/124 to PA). MB-A. was a recipient of a predoctoral fellowship (FPU2017/03542) funded by MCIN/AEI/10.13039/501100011033. We thank the Cytometry Service of the SCSIE-Universitat de València for their help with the cytometry experiments.

## EXPANDED VIEW FIGURE LEGENDS

**Figure EV1.**
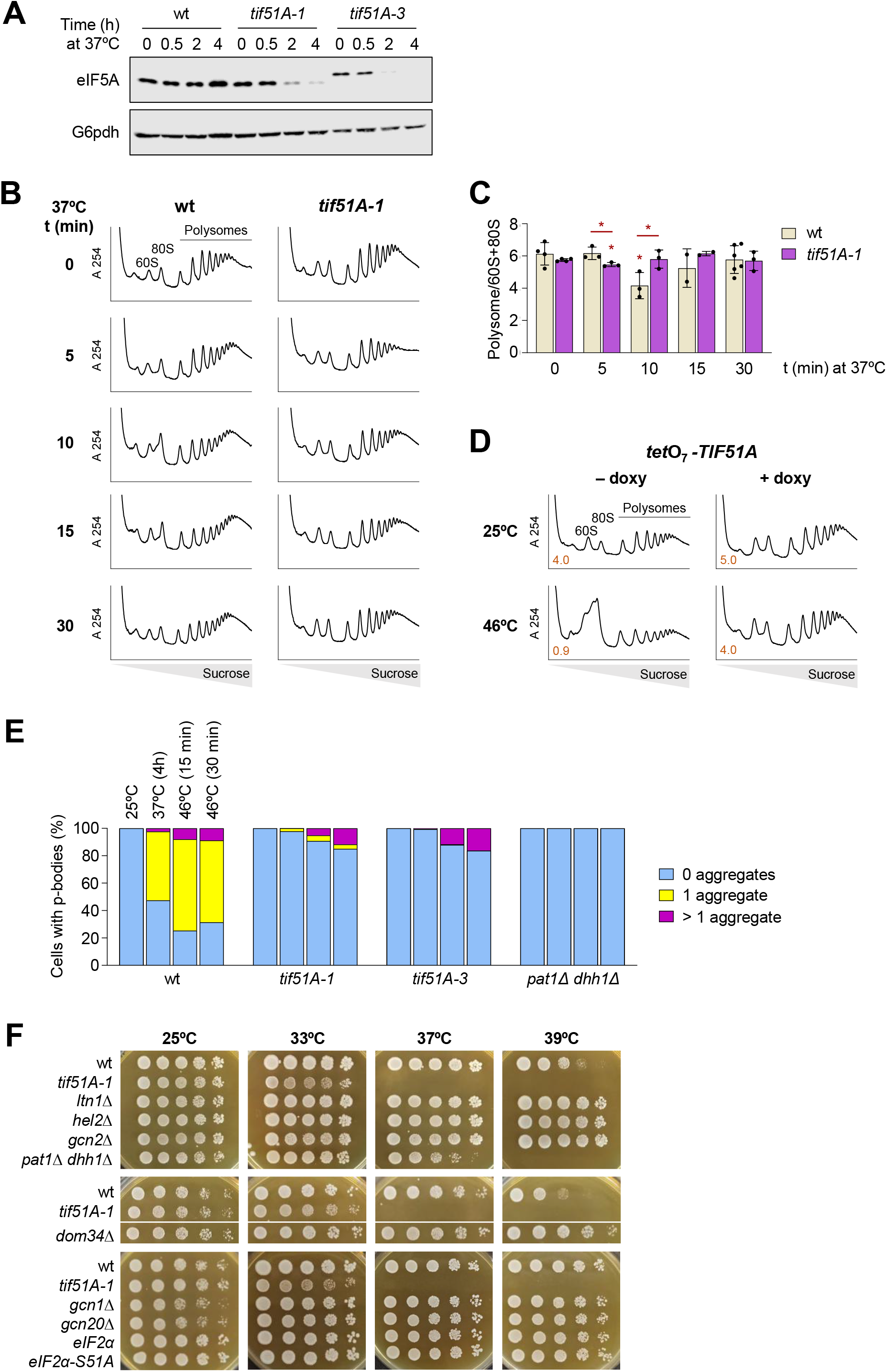
Impact of eIF5A depletion on global translation and p-body dynamics during heat stress. **(A)** Western blot analysis of eIF5A protein levels in wild type (wt), *tif51A-1*, and *tif51A-3* strains. Cells were cultured in YPD medium at 25°C to exponential phase and transferred to 37°C for the indicated times. Specific antibodies against eIF5A and G6pdh were employed, with G6pdh used as a loading control. Note that eIF5A protein in *tif51A-3* mutant runs slower than in wt. **(B)** Polysome profiles of wild type (wt) and *tif51A-1* strains cultured in SC medium at 25°C until the exponential phase and then shifted to 37°C for the indicated times (0, 5, 10, 15, and 30 min). Representative profiles from independent experiments are shown. **(C)** Quantification of the polysome/(60S+80S) area ratio calculated from the polysome profiles shown in **(B). (D)** Polysome profiles under doxycycline depletion of *TIF51A*. The *tetO_7_ -TIF51A* strain was cultured in SC medium at 25°C in the dark with or without 15 µg/mL doxycycline to deplete eIF5A. The following day, doxycycline was added again and cells were maintained at 25°C or transferred to 37°C for 4 h followed by 30 min at 46°C. The calculated polysome/(60S+80S) area ratio is indicated in orange in the lower-left corner of each profile. **(E)** Quantification of p-bodies formation by fluorescence microscopy analysis. The wt, *tif51A-1*, *tif51A-3,* and *pat1*Δ *dhh1*Δ strains expressing Dcp2-RFP were cultured in SC-Leu at 25°C until early exponential phase and shifted to 37°C for 4 h followed by 46°C for 15 and 30 min. Stacked bars represent the percentage of cells displaying 0, 1, or >1 Dcp2-RFP foci (n ≥100 cells per condition). **(F)** Serial dilution spot assays of cells grown in YPD medium at 25°C to the exponential phase and spotted onto YPD plates incubated at the indicated temperatures. Strains eIF2α wild type and eIF2α-S51A denote a *sui2Δ* mutant carrying a plasmid-borne wild type or mutant *sui2-S51A* allele, respectively. The black horizontal line indicates non-adjacent rows grouped from the same original agar plate. **Data information:** In (C), data represent the mean ± SD from independent biological experiments (n = 2 to 4). Individual data points are plotted as dots. Statistical significance was assessed using a two-tailed unpaired Student’s t-test for conditions with n ≥ 3; no statistical tests were performed for the 15 min time point due to limited sample size (n = 2). Asterisks located directly above the bars indicate statistically significant differences compared to time 0 for each respective strain. Asterisks over horizontal lines indicate significant differences between the *tif51A-1* mutant and the wt strain at that specific time point. (\**P* < 0.05). Exact *P* values for all comparisons are provided in the associated Source Data file.

**Figure EV2.**
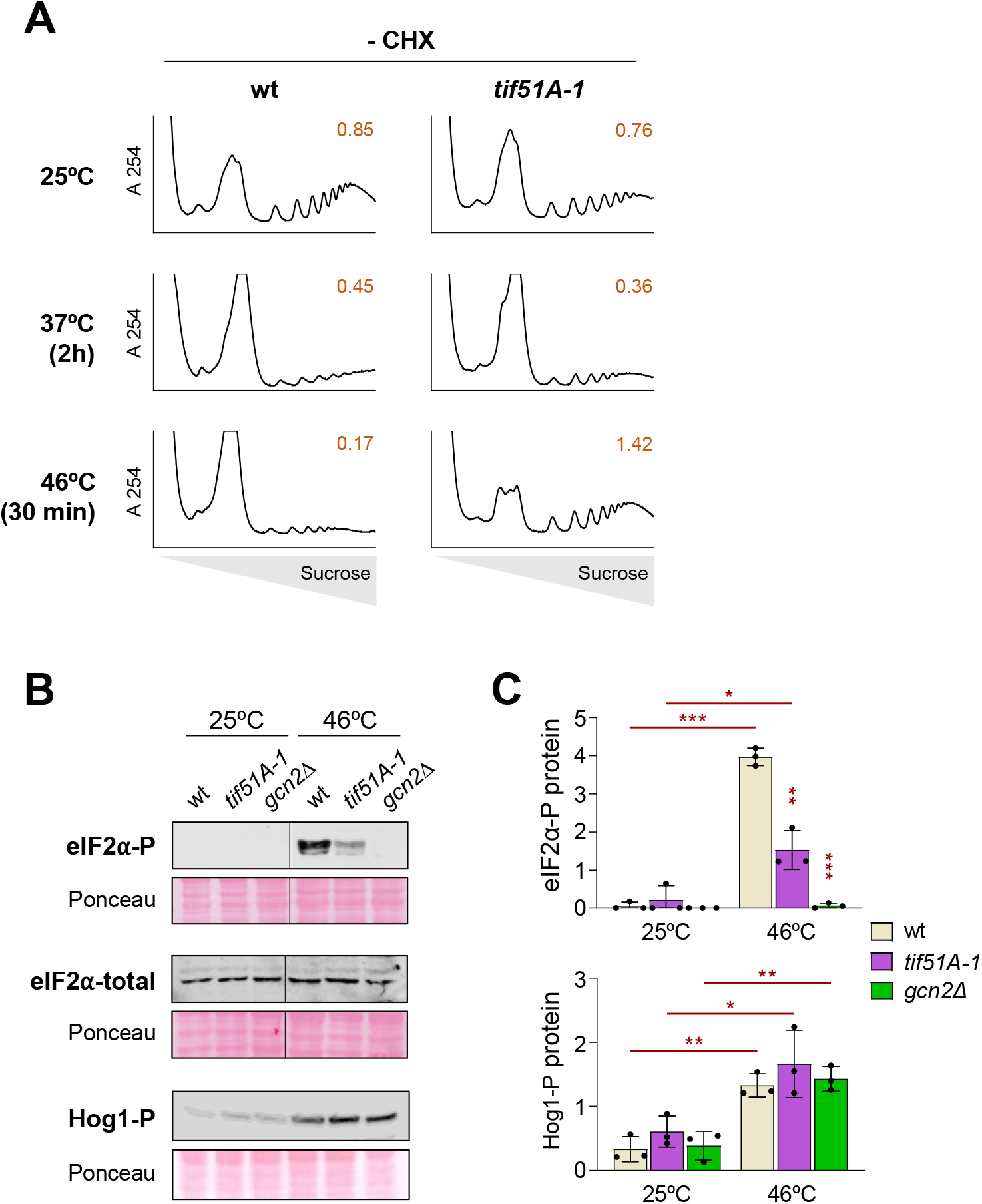
Effect of eIF5A dysfunction in polysomal profiles and eIF2α signalling. **(A)** Representative polysome profiles performed on samples prepared without cycloheximide (CHX). Wild type (wt) and *tif51A-1* strains were cultured in SC medium at 25°C until the exponential phase and then maintained at 25°C, shifted to 37°C for 2 h, or shifted to 37°C for 4 h followed by 30 min at 46°C. The polysome/(60S+80S) area ratio is indicated in orange in the upper-right corner of each profile. **(B)** Wild type, *tif51A-1*, and *gcn2Δ* strains were grown at 25°C to exponential phase and then transferred to 46°C for 30 min. Representative western blots evaluating protein levels of phosphorylated eIF2α (eIF2α-P), total eIF2α, and phosphorylated Hog1 (Hog1-P) using specific antibodies are shown. Ponceau S staining was used as a loading control. Vertical black lines indicate non-adjacent lanes spliced together from the same blot for presentation purposes. **(C)** Quantification of eIF2α and Hog1 phosphorylation levels from the western blots shown in **(B),** normalized to the Ponceau S loading control. **Data information:** In (C), results are presented as the mean ± SD from three independent biological experiments (n = 3). Individual data points are plotted as dots. Statistical significance was determined using a two-tailed unpaired Student’s t-test (\**P* < 0.05, \*\**P* < 0.01, \*\*\**P* < 0.001). Asterisks located directly above individual bars indicate statistically significant differences between the indicated mutant strain and the wt under the same temperature condition. Asterisks positioned above horizontal lines denote statistically significant differences for a given strain when comparing 25°C and 46°C conditions. Exact *P* values for all comparisons are provided in the associated Source Data file.

**Figure EV3.**
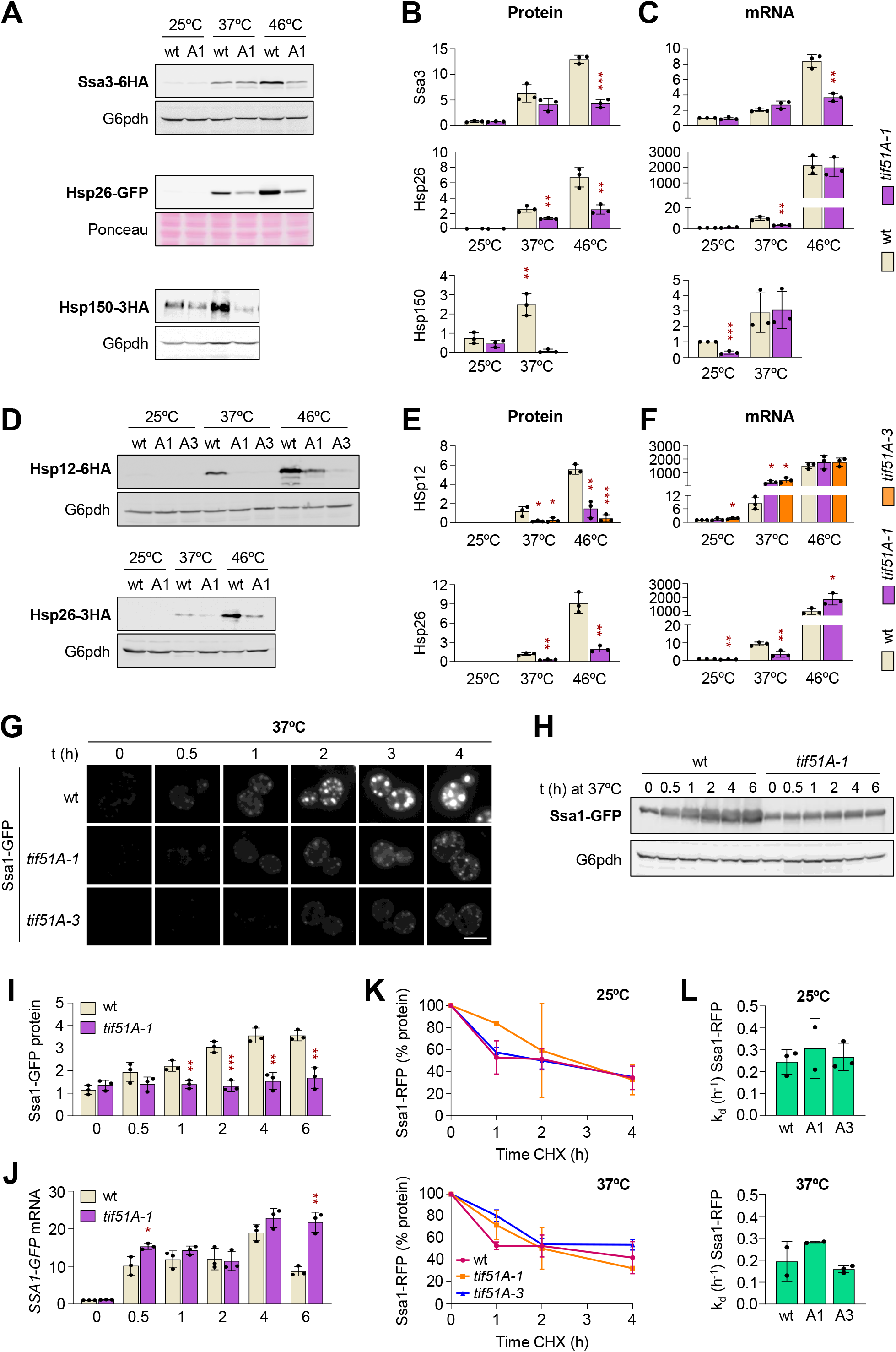
Defective chaperone accumulation in eIF5A mutants under heat stress is independent of protein tags, mRNA induction, and protein stability. **(A-F)** Wild type (wt), *tif51A-1* (A1), and *tif51A-3* (A3) strains were cultured in YPD medium at 25°C until early exponential phase and then shifted to 37°C for 4 h or to 37°C for 4 h followed by 30 min at 46°C. **(A)** Representative Western blots evaluating the protein levels of endogenously tagged Ssa3-6HA, Hsp26-GFP, and Hsp150-3HA in wt and *tif51A-1* strains using anti-HA or anti-GFP antibodies. G6pdh levels or Ponceau S staining was used as a loading control. For Hsp150, only the 25°C and 37°C conditions are shown. **(B)** Quantification of chaperone protein levels from the Western blots shown in **(A),** normalized to the G6pdh or Ponceau S loading control. **(C)** Relative mRNA levels of *SSA3*, *HSP26*, and *HSP150* determined by RT-qPCR under the conditions described in **(A)**, expressed relative to the wt strain at 25°C. **(D)** Representative Western blots evaluating endogenously tagged Hsp12-6HA (in wt, *tif51A-1*, and *tif51A-3*) and Hsp26-3HA (in wt and *tif51A-1*) protein levels using an anti-HA antibody and G6pdh as a loading control. **(E)** Quantification of Hsp12-6HA and Hsp26-3HA protein levels from the Western blots in **(D)**, normalized to G6pdh. **(F)** Relative mRNA levels of *HSP12* and *HSP26* determined by RT-qPCR, expressed relative to the wt strain at 25°C. **(G)** Fluorescence microscopy analysis of Ssa1-GFP over time. The wt, *tif51A-1*, and *tif51A-3* strains expressing Ssa1-GFP were grown in YPD at 25°C to early exponential phase and shifted to 37°C for the indicated times. Representative images from three independent experiments are shown (scale bar: 5 µm). **(H, I)** The wt and *tif51A-1* strains cultured as in **(G)** were harvested at the indicated time points. Ssa1-GFP protein levels were analyzed by Western blotting using an anti-GFP antibody with G6pdh as a loading control **(H)** and quantified **(I)**. **(J)** Relative *SSA1-GFP* mRNA levels determined by RT-qPCR under the heat stress conditions described in **(G)**, expressed relative to the wt strain at time 0. **(K)** Ssa1 protein stability assessed by a cycloheximide (CHX) chase assay. The wt, *tif51A-1*, and *tif51A-3* strains expressing Ssa1-RFP were grown in YPD until exponential phase and then incubated at 25°C or 37°C for 4 h. Then, CHX (100 µg/mL) was added and Ssa1-RFP protein levels were analyzed and quantified by Western blotting at the indicated times. **(L)** Constants of the protein degradation rate (k_d_) were calculated from the CHX chase experiments in **(K)**. **Data information:** For all RT-qPCR experiments (C, F, J), mRNA levels were normalized to endogenous *ACT1* mRNA. In (B, C, E, F, I, J), data represent the mean ± SD from three independent biological experiments (n = 3). In (K, L), data represent the mean ± SD from independent biological experiments (n = 3, except for *tif51A-1* at 25°C and 37°C, and wt at 37°C, where n = 2). Individual data points are plotted as dots in (B, C, E, F, I, J, L) and omitted in (K) to preserve graphical clarity. Statistical significance was assessed using a two-tailed unpaired Student’s *t*-test for conditions with n ≥ 3 (\**P* < 0.05, \*\**P* < 0.01, \*\*\**P* < 0.001). Asterisks indicate statistically significant differences between the indicated mutant and the corresponding wt strain under the same condition or time point. Exact *P* values for all comparisons are provided in the associated Source Data file.

**Figure EV4.**
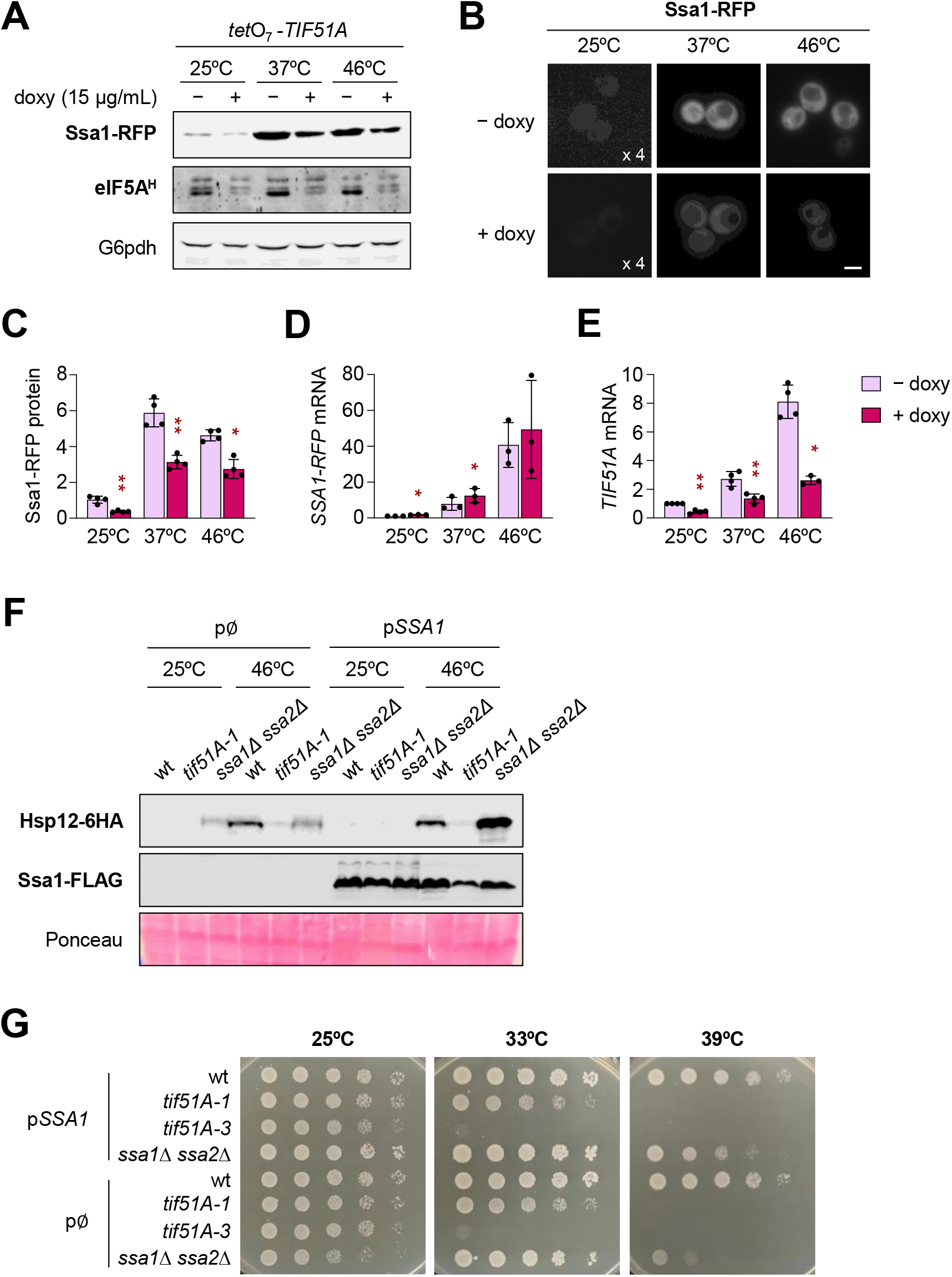
Depletion of *tetO_7_-TIF51A* with doxycycline impairs Ssa1 synthesis under heat stress, but *SSA1* overexpression fails to rescue eIF5A thermosensitivity. **(A-E)** To deplete eIF5A conditionally, a strain expressing *TIF51A* under the control of a doxycycline-repressible promoter (*tetO_7_-TIF51A*) was used. Cells were cultured in YPD medium at 25°C in the dark with or without doxycycline (15 µg/mL) until early exponential phase. The following day, doxycycline was refreshed and cells were shifted to 37°C for 4 h or to 37°C for 4 h followed by 30 min at 46°C. **(A)** Western blot analysis of Ssa1-RFP and hypusinated eIF5A (eIF5A^H^) levels under the indicated temperature and doxycycline conditions using antibodies against RFP or hypusine, with G6pdh used as a loading control. **(B)** Fluorescence microscopy images of yeast cells expressing Ssa1-RFP grown at 25°C, 37°C, and 46°C as previously described with or without doxycycline (scale bar: 5 µm). The label in the 25°C panels indicates that signal brightness was digitally enhanced 4-fold (x4) to allow visualization of basal expression levels. **(C)** Quantification of Ssa1-RFP protein levels from the Western blots in **(A)**, normalized to the G6pdh loading control. **(D, E)** Relative mRNA levels of *SSA1-RFP* (D) and *TIF51A* (E) determined by RT-qPCR under the indicated conditions. Values are normalized to endogenous *ACT1* mRNA and expressed relative to the untreated control (− doxy) at 25°C. **(F)** Representative Western blot evaluating Hsp12-6HA and Ssa1-FLAG protein levels in wild type (wt), *tif51A-1*, and *ssa1Δ ssa2Δ* strains transformed with either an *SSA1* overexpression plasmid (p*SSA1*) or an empty vector control (p∅). Cells were grown in SC-Leu medium at 25°C until exponential phase and then shifted to 37°C for 4 h followed by 30 min at 46°C (indicated as 46°C). Specific antibodies against HA or FLAG were used, and Ponceau S staining was employed as a loading control. **(G)** Ten-fold serial dilution spot assays of wt, *tif51A-1*, *tif51A-3*, and *ssa1Δ ssa2Δ* strains carrying p*SSA1* or p∅. Cells were grown in SC-Leu at 25°C to exponential phase, spotted onto SC-Leu plates, and incubated at the indicated temperatures. **Data information:** In (C–E), data represent the mean ± SD from independent biological experiments (n = 3 to 4). Individual data points are plotted as dots. Statistical significance was calculated using a two-tailed paired Student’s *t*-test (\**P* < 0.05, \*\**P* < 0.01, \*\*\**P* < 0.001). Asterisks indicate statistically significant differences between doxycycline-treated (+ doxy) and untreated (− doxy) samples under the same temperature condition. Exact *P* values for all comparisons are provided in the associated Source Data file.

**Figure EV5.**
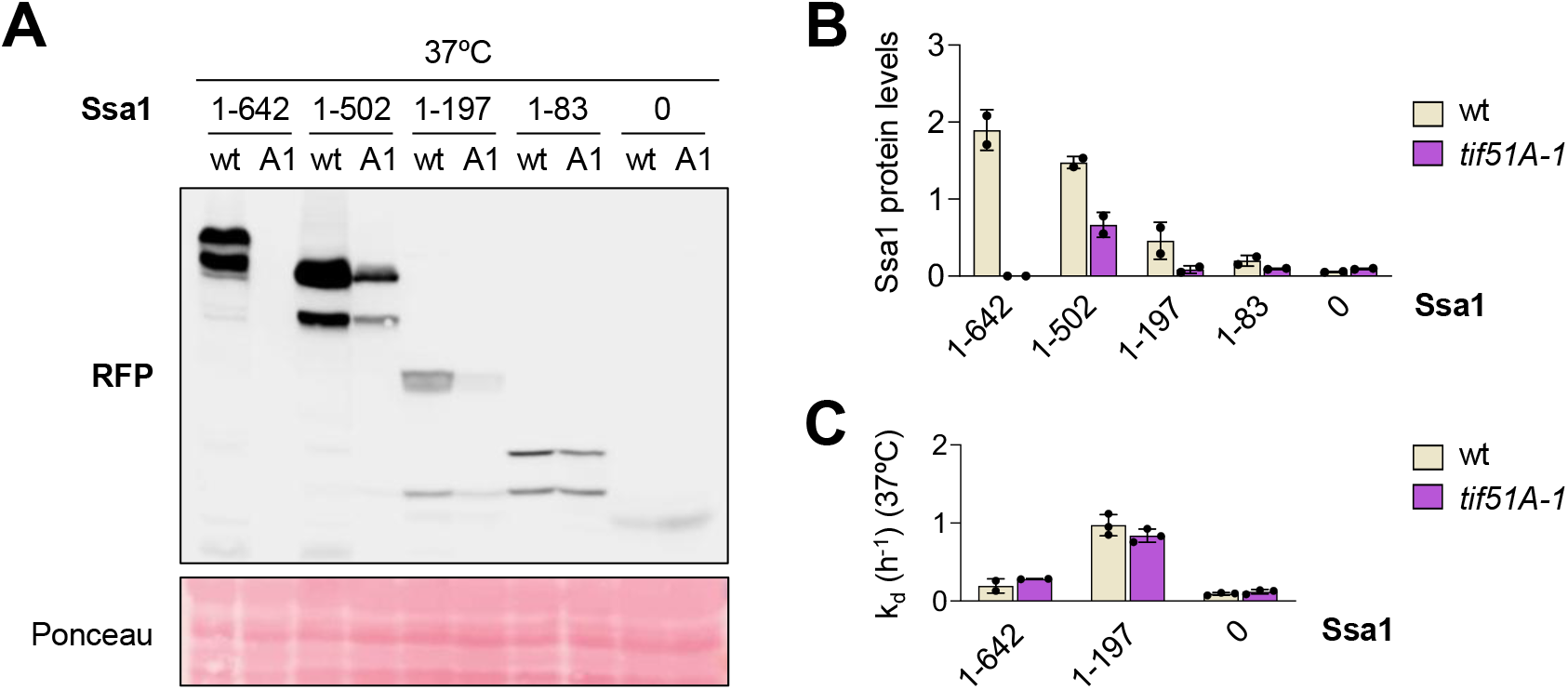
Under heat shock, the *SSA1* translation dependency on eIF5A mutant is linked to the *SSA1* coding region. **(A,B)** Wild type (wt) and *tif51A-1* (A1) cells expressing different Ssa1 truncated proteins (it is indicated the Ssa1 amino acids contained in each construct) fused to RFP, or expressing only the RFP reporter (P*_SSA1_*-RFP, indicated as 0), were grown in YPD at 25°C until early exponential phase and then shifted to 37°C for 4 h. **(A)** Representative western blot evaluating the levels of the Ssa1 versions in wild type and eIF5A mutant. Western was developed using an anti-RFP antibody and Ponceau S staining was used as a loading control. **(B)** Quantification of Ssa1 protein levels from **(A)**, normalized to the Ponceau S loading control. **(C)** Protein stability of Ssa1(1-642), Ssa1(1-197), and only RFP (P*_SSA1_*-RFP, indicated as 0) represented as the degradation rate constant (k_d_). For stability assays, cycloheximide (100 µg/mL) was added after the 4-h incubation at 37°C, and samples were collected at 0, 1, 2, and 4 h post-treatment. **Data information:** Data represent the mean ± SD from independent biological experiments (n = 2 for B; n = 2 to 3 for C). Individual data points are plotted as dots. No statistical tests were applied due to the limited sample size.

