## Appendix Supplementary Figures, Tables and Methods for "Efficient translation of chaperones by eIF5A safeguards proteostasis under heat stress"

**Table of contents**


| | |
|---|---|
| Appendix Figure S1 | Page 2 |
| Appendix Figure S2 | Page 3 |
| Appendix Figure S3 | Pages 4-8 |
| Appendix Figure S4 | Page 9 |
| Appendix Table S1 | Pages 10-11 |
| Appendix Table S2 | Page 12 |
| Appendix Table S3 | Pages 13-14 |
| Appendix Table S4 and references | Page 15 |
| Appendix Supplementary Methods and references | Pages 16-27 |

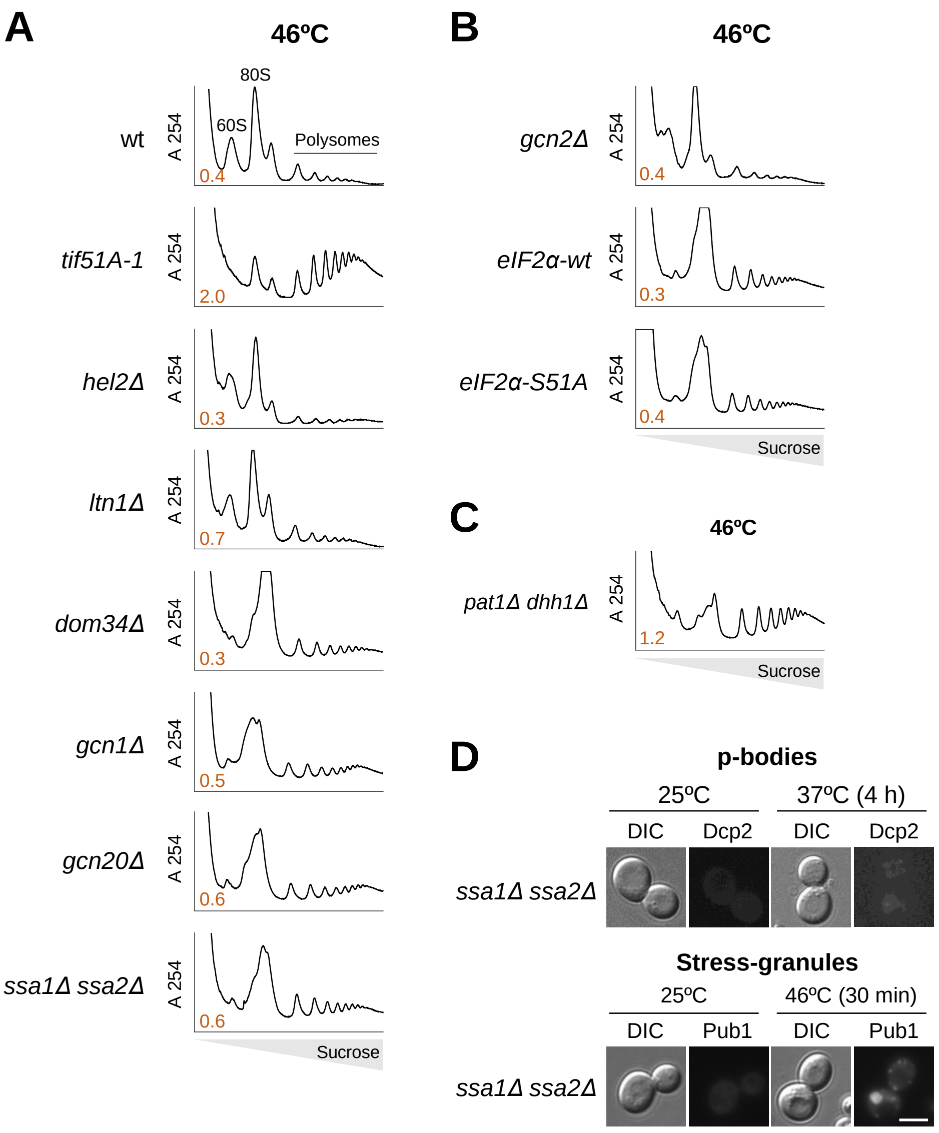


**Appendix Figure S1 – Polysome profile analysis under severe heat stress and assembly of cytosolic foci in chaperone mutants.**

**(A–C)** Representative polysome profiles of the indicated yeast mutant strains under severe heat shock conditions. Wild-type (wt), *tif51A-1*, *hel2Δ*, *ltn1Δ*, *dom34Δ*, *gcn1Δ*, *gcn20Δ*, and *ssa1Δ ssa2Δ* **(A)**, *gcn2Δ*, *eIF2α-wt*, and *eIF2α-S51A* **(B)**, and *pat1Δ dhh1Δ* **(C)** strains were grown in SC medium at 25ºC to early exponential phase, shifted to 37ºC for 4 h, and subsequently subjected to 46ºC for 30 min. In **(B)**, *eIF2α-wt* and *eIF2α-S51A* denote a *sui2Δ* strain carrying a plasmid-borne wild-type *SUI2* or mutant *sui2-S51A* allele, respectively. The calculated polysome/(60S+80S) area ratio is shown in orange in the bottom-left corner of each polysome profile. **(D)** Fluorescence microscopy evaluating the formation of p-bodies and stress granules in the *ssa1Δ ssa2Δ* mutant. Cells expressing Dcp2-RFP (top) or Pub1-mCherry (bottom) were grown to exponential phase at 25ºC in SC-Leu or SC-Ura medium, respectively. Cells were imaged at 25ºC and either shifted to 37ºC for 4 h (p-bodies) or incubated at 37ºC for 4 h followed by heat shock at 46ºC for 30 min (stress granules). Scale bar: 5 µm.

### Appendix Figure S1

**A**

| | PPI | Optimality (%) | eIF5A dependence (37ºC) |
|---|---|---|---|
| Ssa1 | 49.00 | 87.1 | + |
| Ssa2 | 36.31 | 90.0 | + |
| Ssa3 | 19.27 | 50.8 | − |
| Ssa4 | 23.86 | 53.7 | − |
| Hsp150 | 11.23 | 85.7 | + |
| Hsp12 | 3.81 | 85.5 | + |
| Hxk1 | 12.44 | 71.6 | − |
| Ctt1 | 10.14 | 59.0 | − |
| Rtg1 | 12.93 | 48.3 | − |
| GFP | 0 | 49.4 | |
| RFP | 4.73 | 88.2 | |
| YFP | 0 | 71.1 | |
| **Genome** | **10.84** | **47.2** | |

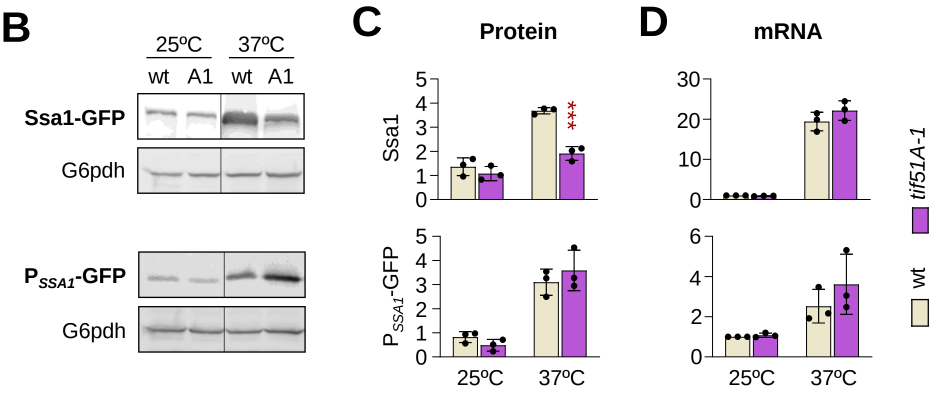


**Appendix Figure S2 – Characteristics and eIF5A translation dependency of heat stress induced mRNAs .**

**(A)** Table summarizing the eIF5A-protein pausing index (PPI), the percentage of codon optimality, and eIF5A translational dependency at 37ºC for heat stress induced and non-induced genes, compared to the genome-wide average. Genes are color-coded based on their optimality: high (green), intermediate (yellow), and low (red). Translation dependency on eIF5A at 37ºC is indicated as (+), and independency as (−), compiled data from Fig 4A-F, EV3A-J and 5H,I. **(B–D)** Wild-type (wt) and the eIF5A temperature-sensitive mutant (*tif51A-1*, A1) strains expressing either endogenously tagged Ssa1-GFP or the $P_{SSA1}$-GFP reporter were cultured in YPD at 25ºC until early exponential phase and then shifted to 37ºC for 4 h. **(B)** Representative western blots evaluating protein levels using an anti-GFP antibody. G6pdh was used as a loading control. Vertical dividing lines indicate spliced non-adjacent lanes from the same membrane. **(C)** Quantification of protein levels from (B), normalized to the G6pdh loading control. **(D)** Corresponding mRNA levels determined by RT-qPCR, normalized to endogenous *ACT1* mRNA and expressed relative to the wt strain at 25ºC.

**Data information:** In (C, D) data represent the mean ± SD from independent biological experiments (n = 3). Individual data points are plotted as dots. Statistical significance was assessed using a two-tailed unpaired Student's t-test (*$P$ < 0.05, **$P$ < 0.01, ***$P$ < 0.001). Asterisks indicate statistically significant differences between the *tif51A-1* mutant and the wt strain under the same temperature condition. Exact *P* values for all comparisons are provided in the associated Source Data file.

### Appendix Figure S2

**A**

```
SSA1(87opt)  ATG TCA AAA GCT GTC GGT ATT GAT TTA GGT ACA ACA TAC TCG TGT GTT GCT CAC TTT GCT
SSA1(54opt)  ATG TCA AAA GCT GTC GGT ATT GAT TTA GGT ACA ACA TAT TCG TGT GTT GCT CAT TTT GCA
SSA4(54opt)  ATG TCA AAA GCT GTT GGT ATT GAT TTA GGT ACA ACC TAT TCA TGT GTT GCT CAT TTT GCA

Ssa1(87opt)   M   S   K   A   V   G   I   D   L   G   T   T   Y   S   C   V   A   H   F   A
Ssa1(54opt)   M   S   K   A   V   G   I   D   L   G   T   T   Y   S   C   V   A   H   F   A
Ssa4(54opt)   M   S   K   A   V   G   I   D   L   G   T   T   Y   S   C   V   A   H   F   A

SSA1(87opt)  AAT GAT CGT GTG GAC ATT ATT GCC AAC GAT CAA GGT AAC AGA ACC ACT CCA TCT TTT GTC
SSA1(54opt)  AAT GAT AGG GTG GAC ATT ATT GCC AAC GAT CAA GGT AAT AGA ACG ACG CCT TCT TTT GTG
SSA4(54opt)  AAC GAT AGG GTT GAA ATT ATC GCT AAC GAT CAA GGT AAT AGA ACG ACG CCT TCT TAT GTG

Ssa1(87opt)   N   D   R   V   D   I   I   A   N   D   Q   G   N   R   T   T   P   S   F   V
Ssa1(54opt)   N   D   R   V   D   I   I   A   N   D   Q   G   N   R   T   T   P   S   F   V
Ssa4(54opt)   N   D   R   V   E   I   I   A   N   D   Q   G   N   R   T   T   P   S   Y   V

SSA1(87opt)  GCT TTC ACT GAC ACT GAA AGA TTG ATT GGT GAT GCT GCT AAG AAT CAA GCT GCT ATG AAT
SSA1(54opt)  GCT TTT ACT GAC ACA GAA AGG CTA ATT GGT GAT GCT GCG AAG AAT CAA GCT GCG ATG AAT
SSA4(54opt)  GCT TTT ACT GAC ACA GAA AGG CTA ATT GGT GAC GCT GCG AAG AAT CAA GCT GCG ATG AAC

Ssa1(87opt)   A   F   T   D   T   E   R   L   I   G   D   A   A   K   N   Q   A   A   M   N
Ssa1(54opt)   A   F   T   D   T   E   R   L   I   G   D   A   A   K   N   Q   A   A   M   N
Ssa4(54opt)   A   F   T   D   T   E   R   L   I   G   D   A   A   K   N   Q   A   A   M   N

SSA1(87opt)  CCT TCG AAT ACC GTT TTC GAC GCT AAG CGT TTG ATC GGT AGA AAC TTC AAC GAC CCA GAA
SSA1(54opt)  CCT TCG AAT ACA GTA TTC GAT GCT AAG CGT CTG ATC GGA AGA AAT TTC AAT GAT CCA GAA
SSA4(54opt)  CCA CAT AAT ACA GTA TTC GAT GCT AAG CGT CTG ATC GGA CGT AAA TTC GAT GAT CCA GAA

Ssa1(87opt)   P   S   N   T   V   F   D   A   K   R   L   I   G   R   N   F   N   D   P   E
Ssa1(54opt)   P   S   N   T   V   F   D   A   K   R   L   I   G   R   N   F   N   D   P   E
Ssa4(54opt)   P   H   N   T   V   F   D   A   K   R   L   I   G   R   K   F   D   D   P   E

SSA1(87opt)  GTG CAG GCT GAC ATG AAG CAC TTC CCA TTC AAG TTG ATC GAT GTT GAC GGT AAG CCT CAA
SSA1(54opt)  GTG CAG GCT GAT ATG AAG CAT TTC CCA TTC AAA TTA ATC GAT GTT GAT GGT AAA CCT CAG
SSA4(54opt)  GTG ACG AAC GAT GCT AAG CAT TAC CCA TTC AAA GTG ATT GAC AAG GGA GGT AAA CCG GTA

Ssa1(87opt)   V   Q   A   D   M   K   H   F   P   F   K   L   I   D   V   D   G   K   P   Q
Ssa1(54opt)   V   Q   A   D   M   K   H   F   P   F   K   L   I   D   V   D   G   K   P   Q
Ssa4(54opt)   V   T   N   D   A   K   H   Y   P   F   K   V   I   D   K   G   G   K   P   V

SSA1(87opt)  ATT CAA GTT GAA TTT AAG GGT GAA ACC AAG AAC TTT ACC CCA GAA CAA ATC TCC TCC ATG
SSA1(54opt)  ATA CAA GTG GAA TTT AAA GGC GAG ACA AAG AAT TTT ACC CCA GAA CAA ATC TCC TCA ATG
SSA4(54opt)  GTG CAA GTG GAA TAT AAA GGC GAG ACA AAG ACA TTT ACT CCA GAA GAA ATT TCC TCA ATG

Ssa1(87opt)   I   Q   V   E   F   K   G   E   T   K   N   F   T   P   E   Q   I   S   S   M
Ssa1(54opt)   I   Q   V   E   F   K   G   E   T   K   N   F   T   P   E   Q   I   S   S   M
Ssa4(54opt)   V   Q   V   E   Y   K   G   E   T   K   T   F   T   P   E   E   I   S   S   M

SSA1(87opt)  GTC TTG GGT AAG ATG AAG GAA ACT GCC GAA TCT TAC TTG GGA GCC AAG GTC AAT GAC GCT
SSA1(54opt)  GTC TTG GGC AAG ATG AAG GAG ACT GCC GAG TCT TAT TTA GGA GCA AAG GTG AAT GAT GCT
SSA4(54opt)  ATC TTG ACA AAG ATG AAG GAG ACT GCT GAG AAC TTT TTA GGA ACA GAA GTG AAA GAT GCT

Ssa1(87opt)   V   L   G   K   M   K   E   T   A   E   S   Y   L   G   A   K   V   N   D   A
Ssa1(54opt)   V   L   G   K   M   K   E   T   A   E   S   Y   L   G   A   K   V   N   D   A
Ssa4(54opt)   I   L   T   K   M   K   E   T   A   E   N   F   L   G   T   E   V   K   D   A

SSA1(87opt)  GTC GTC ACT GTC CCA GCT TAC TTC AAC GAT TCT CAA AGA CAA GCT ACC AAG GAT GCT GGT
SSA1(54opt)  GTA GTA ACG GTC CCA GCT TAT TTC AAC GAT TCA CAA AGG CAA GCA ACA AAA GAT GCT GGT
SSA4(54opt)  GTA GTA ACG GTT CCA GCC TAT TTC AAC GAT TCA CAA AGG CAA GCA ACA AAA GAT GCC GGT

Ssa1(87opt)   V   V   T   V   P   A   Y   F   N   D   S   Q   R   Q   A   T   K   D   A   G
Ssa1(54opt)   V   V   T   V   P   A   Y   F   N   D   S   Q   R   Q   A   T   K   D   A   G
Ssa4(54opt)   V   V   T   V   P   A   Y   F   N   D   S   Q   R   Q   A   T   K   D   A   G
```

### Appendix Figure S3

```
SSA1(87opt)  ACC ATT GCT GGT TTG AAT GTC TTG CGT ATT ATT AAC GAA CCT ACC GCC GCT GCC ATT GCT
SSA1(54opt)  ACA ATT GCG GGC TTG AAT GTC CTT CGT ATT ATT AAT GAA CCT ACA GCC GCT GCC ATT GCG
SSA4(54opt)  ACA ATC GCG GGC TTG AAC GTT CTT CGT ATC ATT AAT GAA CCT ACA GCT GCC GCT ATT GCG

Ssa1(87opt)   T   I   A   G   L   N   V   L   R   I   I   N   E   P   T   A   A   A   I   A
Ssa1(54opt)   T   I   A   G   L   N   V   L   R   I   I   N   E   P   T   A   A   A   I   A
Ssa4(54opt)   T   I   A   G   L   N   V   L   R   I   I   N   E   P   T   A   A   A   I   A

SSA1(87opt)  TAC GGT TTG GAC AAG AAG GG- --T AAG GAA GAA CAC GTC TTG ATT TTC GAC TTG GGT GGT
SSA1(54opt)  TAT GGG CTG GAC AAG AAA GG- --T AAG GAG GAA CAC GTC TTG ATT TTT GAT TTA GGT GGT
SSA4(54opt)  TAT GGG CTG GAC AAG AAA TCG CAG AAG GAG CAC AAC GTC TTG ATC TTT GAT TTA GGT GGT

Ssa1(87opt)   Y   G   L   D   K   K   -   G   K   E   E   H   V   L   I   F   D   L   G   G
Ssa1(54opt)   Y   G   L   D   K   K   -   G   K   E   E   H   V   L   I   F   D   L   G   G
Ssa4(54opt)   Y   G   L   D   K   K   S   Q   K   E   H   N   V   L   I   F   D   L   G   G

SSA1(87opt)  GGT ACT TTC GAT GTC TCT TTG TTG TCC ATT GAA GAC GGT ATC TTT GAA GTT AAG GCC ACC
SSA1(54opt)  GGT ACT TTT GAT GTC TCT CTG CTA TCC ATA GAG GAC GGT ATC TTT GAG GTT AAG GCC ACC
SSA4(54opt)  GGT ACT TTT GAT GTC TCT CTG CTA TCC ATA GAT GAA GGT GTC TTT GAG GTT AAG GCT ACT

Ssa1(87opt)   G   T   F   D   V   S   L   L   S   I   E   D   G   I   F   E   V   K   A   T
Ssa1(54opt)   G   T   F   D   V   S   L   L   S   I   E   D   G   I   F   E   V   K   A   T
Ssa4(54opt)   G   T   F   D   V   S   L   L   S   I   D   E   G   V   F   E   V   K   A   T

SSA1(87opt)  GCT GGT GAC ACC CAT TTG GGT GGT GAA GAT TTT GAC AAC AGA TTG GTC AAC CAC TTC ATC
SSA1(54opt)  GCT GGT GAC ACC CAT TTG GGT GGT GAA GAT TTT GAT AAC AGG CTG GTC AAC CAT TTT ATC
SSA4(54opt)  GCT GGT GAC ACT CAC TTG GGT GGT GAA GAT TTC GAT AGT AGG CTG GTT AAC TTT CTA GCC

Ssa1(87opt)   A   G   D   T   H   L   G   G   E   D   F   D   N   R   L   V   N   H   F   I
Ssa1(54opt)   A   G   D   T   H   L   G   G   E   D   F   D   N   R   L   V   N   H   F   I
Ssa4(54opt)   A   G   D   T   H   L   G   G   E   D   F   D   S   R   L   V   N   F   L   A

SSA1(87opt)  CAA GAA TTC AAG AGA AAG AAC AAG AAG GAC TTG TCT ACC AAC CAA AGA GCT TTG AGA AGA
SSA1(54opt)  CAG GAG TTC AAA AGA AAA AAT AAA AAG GAT CTA TCA ACC AAC CAA AGG GCT CTA AGG AGG
SSA4(54opt)  GAG GAG TTC AAA AGA AAA AAT AAA AAG GAT CTA ACA ACT AAC CAA AGG TCC CTA AGG AGG

Ssa1(87opt)   Q   E   F   K   R   K   N   K   K   D   L   S   T   N   Q   R   A   L   R   R
Ssa1(54opt)   Q   E   F   K   R   K   N   K   K   D   L   S   T   N   Q   R   A   L   R   R
Ssa4(54opt)   E   E   F   K   R   K   N   K   K   D   L   T   T   N   Q   R   S   L   R   R

SSA1(87opt)  TTA AGA ACC GCT TGT GAA AGA GCC AAG AGA ACT TTG TCT TCC TCC GCT CAA ACT TCC GTT
SSA1(54opt)  TTA AGG ACC GCT TGT GAA AGG GCC AAG AGA ACT CTG TCT TCG TCC GCT CAG ACA TCC GTA
SSA4(54opt)  TTA AGG ACC GCC GCT GAA AGG GCC AAG AGA ACT CTG TCT TCG TCT GCT CAG ACA TCT ATA

Ssa1(87opt)   L   R   T   A   C   E   R   A   K   R   T   L   S   S   S   A   Q   T   S   V
Ssa1(54opt)   L   R   T   A   C   E   R   A   K   R   T   L   S   S   S   A   Q   T   S   V
Ssa4(54opt)   L   R   T   A   A   E   R   A   K   R   T   L   S   S   S   A   Q   T   S   I

SSA1(87opt)  GAA ATT GAC TCT TTG TTC GAA GGT ATC GAT TTC TAC ACT TCC ATC ACC AGA GCC AGA TTC
SSA1(54opt)  GAA ATA GAT TCA TTA TTT GAG GGT ATC GAT TTC TAT ACT TCC ATC ACA AGG GCA AGA TTT
SSA4(54opt)  GAA ATA GAT TCA TTA TTT GAG GGT ATC GAT TTC TAT ACT TCC ATT ACA AGG GCA AGA TTT

Ssa1(87opt)   E   I   D   S   L   F   E   G   I   D   F   Y   T   S   I   T   R   A   R   F
Ssa1(54opt)   E   I   D   S   L   F   E   G   I   D   F   Y   T   S   I   T   R   A   R   F
Ssa4(54opt)   E   I   D   S   L   F   E   G   I   D   F   Y   T   S   I   T   R   A   R   F

SSA1(87opt)  GAA GAA TTG TGT GCT GAC TTG TTC AGA TCT ACT TTG GAC CCA GTT GAA AAG GTC TTG AGA
SSA1(54opt)  GAA GAA TTA TGT GCT GAT TTG TTT AGA TCT ACA TTG GAT CCA GTG GAA AAA GTC TTG AGA
SSA4(54opt)  GAA GAA TTA TGT GCT GAT TTG TTT AGA TCT ACA TTG GAG CCA GTG GAA AAA GTT TTG GCT

Ssa1(87opt)   E   E   L   C   A   D   L   F   R   S   T   L   D   P   V   E   K   V   L   R
Ssa1(54opt)   E   E   L   C   A   D   L   F   R   S   T   L   D   P   V   E   K   V   L   R
Ssa4(54opt)   E   E   L   C   A   D   L   F   R   S   T   L   E   P   V   E   K   V   L   A
```

### Appendix Figure S3

| | | | | | | | | | | | | | | | | | | | | |
|---|---|---|---|---|---|---|---|---|---|---|---|---|---|---|---|---|---|---|---|---|
| *SSA1(87opt)* | GAT | GCT | AAA | TTG | GAC | AAA | TCT | CAA | GTC | GAT | GAA | ATT | GTC | TTG | GTC | GGT | GGT | TCT | ACC | AGA |
| *SSA1(54opt)* | GAT | GCA | AAA | TTA | GAT | AAA | TCA | CAA | GTC | GAT | GAA | ATT | GTA | CTG | GTC | GGT | GGT | TCA | ACA | AGA |
| *SSA4(54opt)* | GAT | TCA | AAA | TTA | GAT | AAG | TCA | CAA | ATT | GAT | GAA | ATT | GTA | CTT | GTT | GGT | GGT | TCA | ACA | AGA |
| Ssa1(87opt) | D | A | K | L | D | K | S | Q | V | D | E | I | V | L | V | G | G | S | T | R |
| Ssa1(54opt) | D | A | K | L | D | K | S | Q | V | D | E | I | V | L | V | G | G | S | T | R |
| Ssa4(54opt) | D | S | K | L | D | K | S | Q | I | D | E | I | V | L | V | G | G | S | T | R |

| | | | | | | | | | | | | | | | | | | | | |
|---|---|---|---|---|---|---|---|---|---|---|---|---|---|---|---|---|---|---|---|---|
| *SSA1(87opt)* | ATT | CCA | AAG | GTC | CAA | AAA | TTG | GTC | ACT | GAC | TAC | TTC | AAC | GGT | AAG | GAA | CCA | AAC | AGA | TCT |
| *SSA1(54opt)* | ATT | CCA | AAA | GTA | CAA | AAA | CTG | GTC | ACT | GAT | TAT | TTC | AAT | GGT | AAA | GAA | CCA | AAC | AGA | TCG |
| *SSA4(54opt)* | ATT | CCA | AAA | GTA | CAA | AAA | CTG | GTT | TCT | GAT | TTT | TTC | AAT | GGT | AAA | GAA | CCA | AAC | CGT | TCG |
| Ssa1(87opt) | I | P | K | V | Q | K | L | V | T | D | Y | F | N | G | K | E | P | N | R | S |
| Ssa1(54opt) | I | P | K | V | Q | K | L | V | T | D | Y | F | N | G | K | E | P | N | R | S |
| Ssa4(54opt) | I | P | K | V | Q | K | L | V | S | D | F | F | N | G | K | E | P | N | R | S |

| | | | | | | | | | | | | | | | | | | | | |
|---|---|---|---|---|---|---|---|---|---|---|---|---|---|---|---|---|---|---|---|---|
| *SSA1(87opt)* | ATC | AAC | CCA | GAT | GAA | GCT | GTT | GCT | TAC | GGT | GCT | GCT | GTT | CAA | GCT | GCT | ATT | TTG | ACT | GGT |
| *SSA1(54opt)* | ATC | AAC | CCT | GAT | GAG | GCT | GTT | GCT | TAT | GGT | GCT | GCT | GTA | CAG | GCT | GCT | ATT | TTA | ACG | GGT |
| *SSA4(54opt)* | ATT | AAC | CCT | GAT | GAG | GCC | GTC | GCT | TAT | GGT | GCT | GCC | GTA | CAG | GCT | GCC | ATC | TTA | ACG | GGT |
| Ssa1(87opt) | I | N | P | D | E | A | V | A | Y | G | A | A | V | Q | A | A | I | L | T | G |
| Ssa1(54opt) | I | N | P | D | E | A | V | A | Y | G | A | A | V | Q | A | A | I | L | T | G |
| Ssa4(54opt) | I | N | P | D | E | A | V | A | Y | G | A | A | V | Q | A | A | I | L | T | G |

| | | | | | | | | | | | | | | | | | | | | |
|---|---|---|---|---|---|---|---|---|---|---|---|---|---|---|---|---|---|---|---|---|
| *SSA1(87opt)* | GAC | GAA | TCT | TCC | AAG | ACT | CAA | GAT | CTA | TTG | TTG | TTG | GAT | GTC | GCT | CCA | TTA | TCC | TTG | GGT |
| *SSA1(54opt)* | GAC | GAA | TCG | TCG | AAG | ACT | CAA | GAT | CTA | CTG | TTG | CTG | GAT | GTC | GCA | CCA | TTA | TCC | CTA | GGT |
| *SSA4(54opt)* | GAC | CAG | TCG | TCG | ACG | ACC | CAA | GAT | TTA | CTG | TTG | CTG | GAT | GTT | GCA | CCA | TTA | TCT | CTA | GGT |
| Ssa1(87opt) | D | E | S | S | K | T | Q | D | L | L | L | L | D | V | A | P | L | S | L | G |
| Ssa1(54opt) | D | E | S | S | K | T | Q | D | L | L | L | L | D | V | A | P | L | S | L | G |
| Ssa4(54opt) | D | Q | S | S | T | T | Q | D | L | L | L | L | D | V | A | P | L | S | L | G |

| | | | | | | | | | | | | | | | | | | | | |
|---|---|---|---|---|---|---|---|---|---|---|---|---|---|---|---|---|---|---|---|---|
| *SSA1(87opt)* | ATT | GAA | ACT | GCT | GGT | GGT | GTC | ATG | ACC | AAG | TTG | ATT | CCA | AGA | AAC | TCT | ACC | ATT | CCA | ACA |
| *SSA1(54opt)* | ATT | GAA | ACT | GCA | GGT | GGT | GTC | ATG | ACA | AAG | TTG | ATT | CCA | AGA | AAT | TCG | ACC | ATT | CCA | ACA |
| *SSA4(54opt)* | ATT | GAA | ACT | GCA | GGT | GGT | ATT | ATG | ACA | AAG | TTG | ATC | CCA | AGA | AAT | TCG | ACT | ATC | CCA | ACA |
| Ssa1(87opt) | I | E | T | A | G | G | V | M | T | K | L | I | P | R | N | S | T | I | P | T |
| Ssa1(54opt) | I | E | T | A | G | G | V | M | T | K | L | I | P | R | N | S | T | I | P | T |
| Ssa4(54opt) | I | E | T | A | G | G | I | M | T | K | L | I | P | R | N | S | T | I | P | T |

| | | | | | | | | | | | | | | | | | | | | |
|---|---|---|---|---|---|---|---|---|---|---|---|---|---|---|---|---|---|---|---|---|
| *SSA1(87opt)* | AAG | AAG | TCC | GAG | ATC | TTT | TCC | ACT | TAT | GCT | GAT | AAC | CAA | CCA | GGT | GTC | TTG | ATT | CAA | GTC |
| *SSA1(54opt)* | AAA | AAA | TCG | GAG | ATC | TTT | TCC | ACT | TAT | GCT | GAT | AAC | CAA | CCT | GGT | GTG | TTG | ATA | CAA | GTC |
| *SSA4(54opt)* | AAA | AAA | TCG | GAA | GTG | TTT | TCC | ACC | TAC | GCT | GAC | AAC | CAA | CCT | GGT | GTG | TTG | ATA | CAA | GTT |
| Ssa1(87opt) | K | K | S | E | I | F | S | T | Y | A | D | N | Q | P | G | V | L | I | Q | V |
| Ssa1(54opt) | K | K | S | E | I | F | S | T | Y | A | D | N | Q | P | G | V | L | I | Q | V |
| Ssa4(54opt) | K | K | S | E | V | F | S | T | Y | A | D | N | Q | P | G | V | L | I | Q | V |

| | | | | | | | | | | | | | | | | | | | | |
|---|---|---|---|---|---|---|---|---|---|---|---|---|---|---|---|---|---|---|---|---|
| *SSA1(87opt)* | TTT | GAA | GGT | GAA | AGA | GCC | AAG | ACT | AAG | GAC | AAC | AAC | TTG | TTG | GGT | AAG | TTC | GAA | TTG | AGT |
| *SSA1(54opt)* | TTT | GAG | GGT | GAA | AGG | GCC | AAG | ACA | AAA | GAC | AAC | AAT | CTA | CTG | GGT | AAA | TTT | GAG | TTG | AGT |
| *SSA4(54opt)* | TTT | GAG | GGT | GAA | AGG | ACA | AGG | ACA | AAA | GAC | AAC | AAT | CTA | CTG | GGT | AAA | TTT | GAG | TTG | AGC |
| Ssa1(87opt) | F | E | G | E | R | A | K | T | K | D | N | N | L | L | G | K | F | E | L | S |
| Ssa1(54opt) | F | E | G | E | R | A | K | T | K | D | N | N | L | L | G | K | F | E | L | S |
| Ssa4(54opt) | F | E | G | E | R | T | R | T | K | D | N | N | L | L | G | K | F | E | L | S |

| | | | | | | | | | | | | | | | | | | | | |
|---|---|---|---|---|---|---|---|---|---|---|---|---|---|---|---|---|---|---|---|---|
| *SSA1(87opt)* | GGT | ATT | CCA | CCA | GCT | CCA | AGA | GGT | GTC | CCA | CAA | ATT | GAA | GTC | ACT | TTC | GAT | GTC | GAC | TCT |
| *SSA1(54opt)* | GGT | ATT | CCA | CCC | GCT | CCA | AGA | GGC | GTA | CCA | CAA | ATT | GAA | GTC | ACA | TTT | GAT | GTC | GAT | TCT |
| *SSA4(54opt)* | GGT | ATT | CCA | CCC | GCT | CCA | AGA | GGC | GTA | CCA | CAA | ATT | GAA | GTT | ACA | TTT | GAT | ATC | GAT | GCA |
| Ssa1(87opt) | G | I | P | P | A | P | R | G | V | P | Q | I | E | V | T | F | D | V | D | S |
| Ssa1(54opt) | G | I | P | P | A | P | R | G | V | P | Q | I | E | V | T | F | D | V | D | S |
| Ssa4(54opt) | G | I | P | P | A | P | R | G | V | P | Q | I | E | V | T | F | D | I | D | A |

### Appendix Figure S3

```
SSA1(87opt)  AAC GGT ATT TTG AAT GTT TCC GCC GTC GAA AAG GGT ACT GGT AAG TCT AAC AAG ATC ACT
SSA1(54opt)  AAT GGT ATT CTG AAT GTA TCC GCC GTC GAA AAA GGT ACT GGT AAA TCT AAC AAG ATC ACA
SSA4(54opt)  AAT GGT ATT CTG AAC GTA TCT GCC GTT GAA AAA GGT ACT GGT AAA TCT AAC AAG ATT ACA

Ssa1(87opt)   N   G   I   L   N   V   S   A   V   E   K   G   T   G   K   S   N   K   I   T
Ssa1(54opt)   N   G   I   L   N   V   S   A   V   E   K   G   T   G   K   S   N   K   I   T
Ssa4(54opt)   N   G   I   L   N   V   S   A   V   E   K   G   T   G   K   S   N   K   I   T

SSA1(87opt)  ATT ACC AAC GAC AAG GGT AGA TTG TCC AAG GAA GAT ATC GAA AAG ATG GTT GCT GAA GCC
SSA1(54opt)  ATT ACC AAC GAT AAG GGA AGA TTA TCG AAG GAA GAT ATC GAA AAA ATG GTT GCT GAG GCA
SSA4(54opt)  ATT ACT AAC GAT AAG GGA AGA TTA TCG AAG GAA GAT ATC GAT AAA ATG GTT GCT GAG GCA

Ssa1(87opt)   I   T   N   D   K   G   R   L   S   K   E   D   I   E   K   M   V   A   E   A
Ssa1(54opt)   I   T   N   D   K   G   R   L   S   K   E   D   I   E   K   M   V   A   E   A
Ssa4(54opt)   I   T   N   D   K   G   R   L   S   K   E   D   I   D   K   M   V   A   E   A

SSA1(87opt)  GAA AAA TTC AAG GAA GAA GAT GAA AAG GAA TCT CAA AGA ATT GCT TCC AAG AAC CAA TTG
SSA1(54opt)  GAA AAA TTC AAG GAA GAA GAT GAA AAG GAA TCT CAA AGA ATT GCT TCC AAG AAT CAG CTA
SSA4(54opt)  GAA AAG TTC AAG GCC GAA GAT GAA CAA GAA GCT CAA CGT GTT CAA GCT AAG AAT CAG CTA

Ssa1(87opt)  E   K   F   K   E   E   D   E   K   E   S   Q   R   I   A   S   K   N   Q   L
Ssa1(54opt)  E   K   F   K   E   E   D   E   K   E   S   Q   R   I   A   S   K   N   Q   L
Ssa4(54opt)  E   K   F   K   A   E   D   E   Q   E   A   Q   R   V   Q   A   K   N   Q   L

SSA1(87opt)  GAA TCC ATT GCT TAC TCT TTG AAG AAC ACC ATT TCT GAA GCT GGT GAC AAA TTG GAA CAA
SSA1(54opt)  GAA TCG ATT GCG TAC TCT TTG AAA AAT ACC ATT AGC GAA GCT GGT GAC AAA TTG GAA CAA
SSA4(54opt)  GAA TCG TAC GCG TTT ACT TTG AAA AAT TCT GTG AGC GAA AAT AAC TTC AAG GAG AAG GTG

Ssa1(87opt)   E   S   I   A   Y   S   L   K   N   T   I   S   E   A   G   D   K   L   E   Q
Ssa1(54opt)   E   S   I   A   Y   S   L   K   N   T   I   S   E   A   G   D   K   L   E   Q
Ssa4(54opt)   E   S   Y   A   F   T   L   K   N   S   V   S   E   N   N   F   K   E   K   V

SSA1(87opt)  GCT GAC AAG GAC ACC GTC ACC AAG AAG --- --- GCT GAA GAG ACT ATT TCT TGG TTA GAC
SSA1(54opt)  GCT GAC AAG GAT ACC GTG ACA AAG AAG --- --- GCT GAA GAG ACT ATA TCA TGG TTA GAT
SSA4(54opt)  GGT GAA GAG GAT GCC AGG AAA TTG GAA GCC GCC GCC CAA GAT GCT ATA AAT TGG TTA GAT

Ssa1(87opt)   A   D   K   D   T   V   T   K   K   -   -   A   E   E   T   I   S   W   L   D
Ssa1(54opt)   A   D   K   D   T   V   T   K   K   -   -   A   E   E   T   I   S   W   L   D
Ssa4(54opt)   G   E   E   D   A   R   K   L   E   A   A   A   Q   D   A   I   N   W   L   D

SSA1(87opt)  AGC AAC ACC ACT GCC AGC AAG GAA GAA TTC GAT GAC AAG TTG AAG GAG TTG CAA GAC ATT
SSA1(54opt)  AGC AAC ACC ACT GCC AGC AAG GAG GAA TTC GAT GAC AAG TTG AAG GAG CTA CAA GAC ATT
SSA4(54opt)  GCT TCG CAA GCG GCC TCC ACC GAG GAA TAC AAG GAA AGG CAA AAG GAA CTA GAA GGT GTT

Ssa1(87opt)   S   N   T   T   A   S   K   E   E   F   D   D   K   L   K   E   L   Q   D   I
Ssa1(54opt)   S   N   T   T   A   S   K   E   E   F   D   D   K   L   K   E   L   Q   D   I
Ssa4(54opt)   A   S   Q   A   A   S   T   E   E   Y   K   E   R   Q   K   E   L   E   G   V

SSA1(87opt)  GCC AAC CCA ATC ATG TCT AAG TTG TAC CAA GCT --- GGT GGT GCT CCA GGT GGC GCT GCA
SSA1(54opt)  GCA AAC CCC ATC ATG AGT AAA TTG TAC CAA GCT --- GGT GGT GCT CCA GGA GGC GCT GCA
SSA4(54opt)  GCA AAC CCC ATT ATG AGT AAA TTT TAC GGA GCT GCA GGT GGT GCC CCA GGA G-- --- -CA

Ssa1(87opt)   A   N   P   I   M   S   K   L   Y   Q   A   -   G   G   A   P   G   G   A   A
Ssa1(54opt)   A   N   P   I   M   S   K   L   Y   Q   A   -   G   G   A   P   G   G   A   A
Ssa4(54opt)   A   N   P   I   M   S   K   F   Y   G   A   A   G   G   A   P   G   -   -   A

SSA1(87opt)  GGT GGT GCT CCA GGC GGT TTC CCA GGT GGT GCT CCT CCA GCT CCA GAG GCT GAA GGT CCA
SSA1(54opt)  GGC GGT GCT CCG GGC GGT TTC CCA GGA GGT GCT CCT CCA GCA CCA GAG GCT GAA GGC CCA
SSA4(54opt)  GGC CCA GTT CCG GGT G-- --- -CT GGA GCA GGC CCC ACT GGA GCA CCA GAC AAC GGC CCA

Ssa1(87opt)   G   G   A   P   G   G   F   P   G   G   A   P   P   A   P   E   A   E   G   P
Ssa1(54opt)   G   G   A   P   G   G   F   P   G   G   A   P   P   A   P   E   A   E   G   P
Ssa4(54opt)   G   P   V   P   G   -   -   A   G   A   G   P   T   G   A   P   D   N   G   P
```

### Appendix Figure S3

```
SSA1(87opt)  ACC GTT GAA GAA GTT GAT TAA
SSA1(54opt)  ACG GTT GAA GAG GTT GAT TAA
SSA4(54opt)  ACG GTT GAA GAG GTT GAT TAG

Ssa1(87opt)   T   V   E   E   V   D   -
Ssa1(54opt)   T   V   E   E   V   D   -
Ssa4(54opt)   T   V   E   E   V   D   -
```

**B**

```
HSP12(86opt)   ATG TCT GAC GCA GGT AGA AAA GGA TTC GGT GAA AAA GCT TCT GAA GCT TTG AAG CCA GAC
HSP12(45opt)   ATG TCT GAC GCA GGT AGA AAA GGA TTT GGT GAA AAA GCT TCA GAA GCA TTA AAG CCT GAC

Hsp12(86opt)    M   S   D   A   G   R   K   G   F   G   E   K   A   S   E   A   L   K   P   D
Hsp12(45opt)    M   S   D   A   G   R   K   G   F   G   E   K   A   S   E   A   L   K   P   D

HSP12(86opt)   TCT CAA AAG TCA TAC GCT GAA CAA GGT AAG GAA TAC ATC ACT GAC AAG GCC GAC AAG GTC
HSP12(45opt)   TCG CAA AAG TCA TAT GCT GAG CAG GGT AAA GAA TAT ATA ACG GAC AAG GCG GAC AAG GTG

Hsp12(86opt)    S   Q   K   S   Y   A   E   Q   G   K   E   Y   I   T   D   K   A   D   K   V
Hsp12(45opt)    S   Q   K   S   Y   A   E   Q   G   K   E   Y   I   T   D   K   A   D   K   V

HSP12(86opt)   GCT GGT AAG GTT CAA CCA GAA GAC AAC AAG GGT GTC TTC CAA GGT GTC CAC GAC TCT GCC
HSP12(45opt)   GCG GGC AAG GTT CAG CCA GAG GAC AAT AAG GGC GTA TTC CAG GGA GTA CAC GAT TCA GCC

Hsp12(86opt)    A   G   K   V   Q   P   E   D   N   K   G   V   F   Q   G   V   H   D   S   A
Hsp12(45opt)    A   G   K   V   Q   P   E   D   N   K   G   V   F   Q   G   V   H   D   S   A

HSP12(86opt)   GAA AAA GGC AAG GAT AAC GCT GAA GGT CAA GGT GAA TCT TTG GCA GAC CAA GCT AGA GAT
HSP12(45opt)   GAA AAA GGC AAG GAT AAC GCT GAG GGG CAG GGT GAG TCT CTG GCA GAC CAG GCT AGA GAT

Hsp12(86opt)    E   K   G   K   D   N   A   E   G   Q   G   E   S   L   A   D   Q   A   R   D
Hsp12(45opt)    E   K   G   K   D   N   A   E   G   Q   G   E   S   L   A   D   Q   A   R   D

HSP12(86opt)   TAC ATG GGA GCC GCC AAG TCC AAG TTG AAC GAT GCC GTC GAA TAT GTT TCC GGT CGT GTC
HSP12(45opt)   TAC ATG GGA GCC GCG AAA TCC AAG TTA AAC GAT GCA GTC GAA TAT GTA TCG GGT CGT GTG

Hsp12(86opt)    Y   M   G   A   A   K   S   K   L   N   D   A   V   E   Y   V   S   G   R   V
Hsp12(45opt)    Y   M   G   A   A   K   S   K   L   N   D   A   V   E   Y   V   S   G   R   V

HSP12(86opt)   CAC GGT GAA GAA GAC CCA ACC AAG AAG TAA
HSP12(45opt)   CAC GGA GAG GAA GAC CCT ACG AAG AAG TAA

Hsp12(86opt)    H   G   E   E   D   P   T   K   K   *
Hsp12(45opt)    H   G   E   E   D   P   T   K   K   *
```

**Appendix Figure S3 − Nucleotide and amino acid sequence alignments of native and codon-deoptimized *SSA1* and *HSP12* coding regions.**
**(A)** Sequence alignment showing the nucleotide coding sequences and translated amino acids of native *SSA1(87opt)* (87% codon optimality), the codon-deoptimized *SSA1(54opt)* variant (54% codon optimality), and the naturally non-optimal *SSA4(54opt)* sequence (54% codon optimality). Non-optimal codons and their corresponding amino acids are highlighted in red color. Yellow shading indicates synonymous substitutions in *SSA1* where an optimal codon was replaced by a non-optimal synonymous codón, matching the synonymous non-optimal codon present in *SSA4* at the same position without altering amino acid identity. Cyan shading indicates synonymous substitutions in *SSA1* where an optimal codon was replaced by a non-optimal codon at positions where the corresponding *SSA4* codon is also non-optimal but encodes a different amino acid residue. **(B)** Pairwise sequence alignment of nucleotide and amino acid sequences between native *HSP12(86opt)* (86% codon optimality) and the codon-deoptimized *HSP12(45opt)* variant (45% codon optimality). Non-optimal codons and their corresponding translated amino acids are indicated in red color.

### Appendix Figure S3

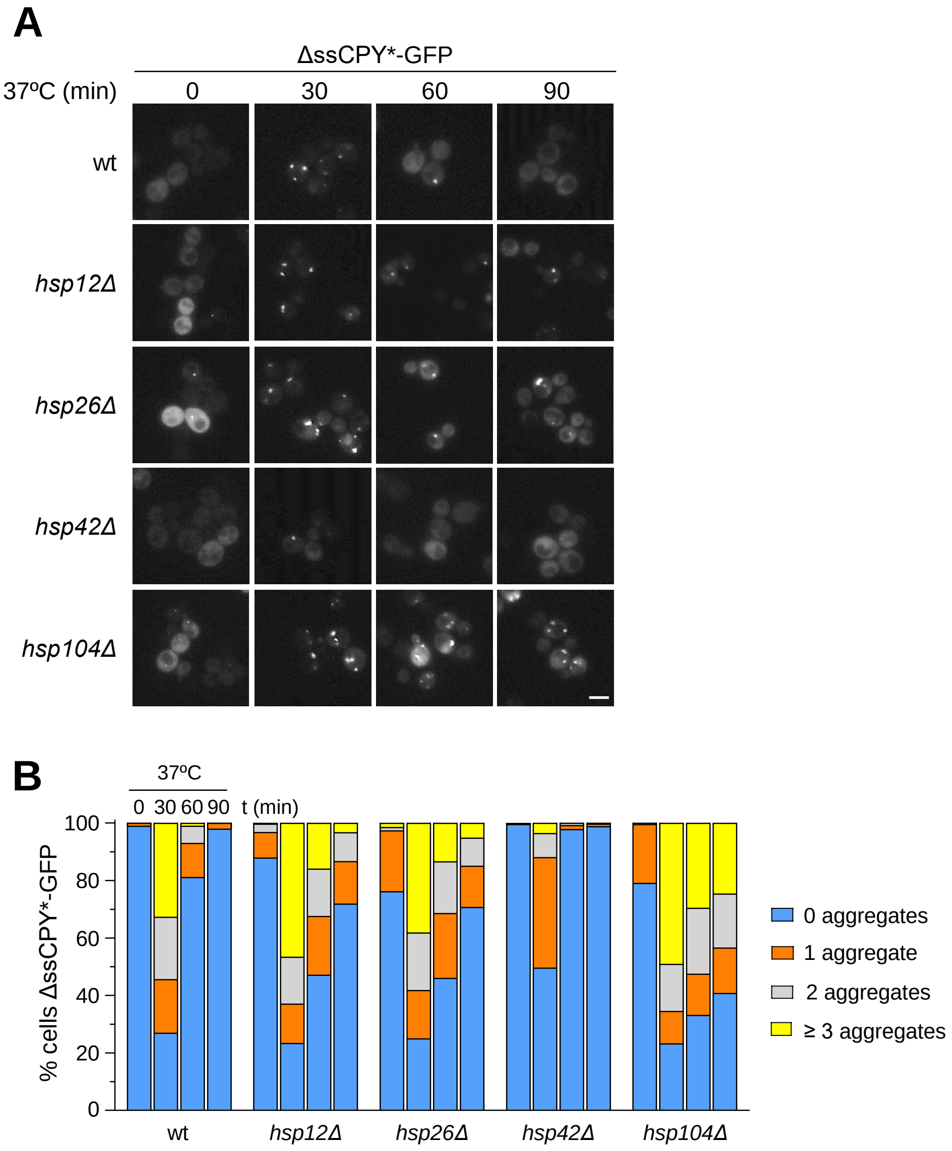


**Appendix Figure S4 – Loss of individual heat shock proteins exacerbates misfolded protein aggregation under thermal stress.** Representative fluorescence microscopy images of wild-type (wt) and the indicated chaperone deletion strains (*hsp12*Δ, *hsp26*Δ, *hsp42*Δ, and *hsp104*Δ) expressing the ΔssCPY*-GFP aggregation reporter. Cells were cultured in SC-Ura medium at 25ºC to the early exponential phase and then shifted to 37ºC. Images were taken at 0, 30, 60 and 90 min after the temperature shift. Scale bar: 5 µm. These images correspond to the supplementary strains quantified in the aggregate analysis presented in Fig 8B.
**Data information:** Images are representative of three independent biological experiments (n ≥ 100 cells were scored per strain and condition).

### Appendix Figure S4

**Appendix Table S1. Yeast strains used in this study**

| Name | Genotype | Source |
|---|---|---|
| **BY4741** | MATa *ura3Δ0 leu2Δ0 his3Δ1 met15Δ0* | Euroscarf |
| ***tif51A-1*** | BY4741 MATa *ura3Δ0 leu2Δ0 his3Δ1 met15Δ0 tif51A-1::kanR* | (Li *et al*, 2011) |
| ***tif51A-3*** | BY4741 MATa *ura3Δ0 leu2Δ0 his3Δ1 met15Δ0 tif51A-3::kanR* | (Li *et al*, 2011) |
| ***dom34Δ*** | BY4741 MATa *ura3Δ0 leu2Δ0 his3Δ1 met15Δ0 dom34Δ::kanMX4* | Euroscarf |
| ***gcn1Δ*** | BY4741 MATa *ura3Δ0 leu2Δ0 his3Δ1 met15Δ0 gcn1Δ::kanMX4* | Euroscarf |
| ***gcn2Δ*** | BY4741 MATa *ura3Δ0 leu2Δ0 his3Δ1 met15Δ0 gcn2Δ::kanMX4* | Euroscarf |
| ***gcn20Δ*** | BY4741 MATa *ura3Δ0 leu2Δ0 his3Δ1 met15Δ0 gcn20Δ::kanMX4* | Euroscarf |
| ***hel2Δ*** | BY4741 MATa *ura3Δ0 leu2Δ0 his3Δ1 met15Δ0 hel2Δ::kanMX6* | This study |
| ***ltn1Δ*** | BY4741 MATa *ura3Δ0 leu2Δ0 his3Δ1 met15Δ0 ltn1Δ::kanMX4* | Euroscarf |
| ***ssa1Δ ssa2Δ*** | BY4741 MATa *ura3Δ0 leu2Δ0 his3Δ1 met15Δ0 ssa1Δ::hphMX4 ssa2Δ::kanMX4* | (Öling *et al*, 2014) |
| ***SUI2*** | RS-86: MATa *ura3-52 leu2-3,112 trp1Δ63 Δsui2 Δp919 [pRS414-SUI2 TRP1]* | (Caballero-Molada *et al*, 2018) |
| ***SUI2-S51A*** | RS-88: MATa *ura3-52 leu2-3,112 trp1Δ63 Δsui2 Δp919 [pRS414 SUI2-S51A TRP1]* | (Caballero-Molada *et al*, 2018) |
| **PAY778** | BY4741 MATa *ura3Δ0 leu2Δ0 his3Δ1 met15Δ0* $tetO_7$*-TIF51A-kanMX4* | (Barba-Aliaga *et al*, 2026) |
| **PAY789** | HHY13a MATα *ura3Δ0 leu2Δ0 his3Δ1 trp1Δ63 can1 tif51A::LEU2 tif51B::HIS3* [p414GAL-mouse-eIF5A, TRP1] | (Clement *et al*, 2003) |
| **PAY819** | BY4741 MATa *ura3Δ0 leu2Δ0 his3Δ1 met15Δ0 hsp104Δ::kanMX4* | Euroscarf |
| **PAY903** | BY4741 MATa *ura3Δ0 leu2Δ0 his3Δ1 met15Δ0 SSA3-6HA-his3MX6* | This study |
| **PAY904** | BY4741 MATa *ura3Δ0 leu2Δ0 his3Δ1 met15Δ0 tif51A-1::kanR SSA3-6HA-his3MX6* | This study |
| **PAY985** | BY4741 MATa *ura3Δ0 leu2Δ0 his3Δ1 met15Δ0 HSP26-3HA-his3MX6* | This study |
| **PAY987** | BY4741 MATa *ura3Δ0 leu2Δ0 his3Δ1 met15Δ0 tif51A-1::kanR HSP26-3HA-his3MX6* | This study |
| **PAY1039** | BY4741 MATa *ura3Δ0 leu2Δ0 his3Δ1 met15Δ0 hsp12Δ::kanMX4* | Euroscarf |
| **PAY1040** | BY4741 MATa *ura3Δ0 leu2Δ0 his3Δ1 met15Δ0 hsp26Δ::kanMX4* | Euroscarf |
| **PAY1083** | BY4741 MATa *ura3Δ0 leu2Δ0 his3Δ1 met15Δ0 GFP-TIF51A-his3MX6* | (Barba-Aliaga *et al*, 2026) |
| **PAY1115** | BY4741 MATa *ura3Δ0 leu2Δ0 his3Δ1 met15Δ0 HSP12-6HA-his3MX6* | This study |
| **PAY1119** | BY4741 MATa *ura3Δ0 leu2Δ0 his3Δ1 met15Δ0 tif51A-1::kanR HSP12-6HA-his3MX6* | This study |
| **PAY1121** | BY4741 MATa *ura3Δ0 leu2Δ0 his3Δ1 met15Δ0 tif51A-3::kanR HSP12-6HA-his3MX6* | This study |
| **PAY1152** | BY4741 MATa *ura3Δ0 leu2Δ0 his3Δ1 met15Δ0 HSP104-RFP-clonNAT* | (Barba-Aliaga *et al*, 2024) |
| **PAY1156** | BY4741 MATa *ura3Δ0 leu2Δ0 his3Δ1 met15Δ0 tif51A-1::kanR HSP104-RFP-clonNAT* | (Barba-Aliaga *et al*, 2024) |
| **PAY1157** | BY4741 MATa *ura3Δ0 leu2Δ0 his3Δ1 met15Δ0 tif51A-3::kanR HSP104-RFP-clonNAT* | (Barba-Aliaga *et al*, 2024) |
| **PAY1183** | BY4741 MATa *ura3Δ0 leu2Δ0 his3Δ1 met15Δ0 hsp42Δ::kanMX4* | Euroscarf |
| **PAY1300** | BY4741 MATa *ura3Δ0 leu2Δ0 his3Δ1 met15Δ0 4xHSEpr-YFP-URA3* | This study |
| **PAY1304** | BY4741 MATa *ura3Δ0 leu2Δ0 his3Δ1 met15Δ0 tif51A-1::kanR 4xHSEpr-YFP URA3* | This study |
| **PAY1317** | BY4741 MATa *ura3Δ0 leu2Δ0 his3Δ1 met15Δ0 SSA2-RFP-clonNAT* | This study |
| **PAY1319** | BY4741 MATa *ura3Δ0 leu2Δ0 his3Δ1 met15Δ0 tif51A-1::kanR SSA2-RFP-clonNAT* | This study |
| **PAY1322** | BY4741 MATa *ura3Δ0 leu2Δ0 his3Δ1 met15Δ0 tif51A-3::kanR SSA2-RFP-clonNAT* | This study |
| **PAY1529** | BY4741 MATa *ura3Δ0 leu2Δ0 his3Δ1 met15Δ0* $tetO_7$*-TIF51A-kanMX4 SSA1-RFP-clonNAT* | This study |
| **PAY1532** | BY4741 MATa *ura3Δ0 leu2Δ0 his3Δ1 met15Δ0 pSSA1-GFP-his3MX6* | This study |
| **PAY1535** | BY4741 MATa *ura3Δ0 leu2Δ0 his3Δ1 met15Δ0 tif51A-1::kanR pSSA1-GFP-his3MX6* | This study |
| **PAY1607** | BY4741 MATa *ura3Δ0 leu2Δ0 his3Δ1 met15Δ0 SSA1-GFP-his3MX6* | This study |
| **PAY1610** | BY4741 MATa *ura3Δ0 leu2Δ0 his3Δ1 met15Δ0 tif51A-1::kanR SSA1-GFP-his3MX6* | This study |
| **PAY1613** | BY4741 MATa *ura3Δ0 leu2Δ0 his3Δ1 met15Δ0 tif51A-3::kanR SSA1-GFP-his3MX6* | This study |
| **PAY1619** | BY4741 MATa *ura3Δ0 leu2Δ0 his3Δ1 met15Δ0 SSA1-RFP-clonNAT* | This study |
| **PAY1622** | BY4741 MATa *ura3Δ0 leu2Δ0 his3Δ1 met15Δ0 tif51A-1::kanR SSA1-RFP-clonNAT* | This study |
| **PAY1625** | BY4741 MATa *ura3Δ0 leu2Δ0 his3Δ1 met15Δ0 tif51A-3::kanR SSA1-RFP-clonNAT* | This study |
| **PAY1648** | BY4741 MATa *ura3Δ0 leu2Δ0 his3Δ1 met15Δ0 pSSA1-RFP-clonNAT* | This study |
| **PAY1651** | BY4741 MATa *ura3Δ0 leu2Δ0 his3Δ1 met15Δ0 tif51A-1::kanR pSSA1-RFP-clonNAT* | This study |
| **PAY1654** | BY4741 MATa *ura3Δ0 leu2Δ0 his3Δ1 met15Δ0 tif51A-3::kanR pSSA1-RFP-clonNAT* | This study |

| | | |
|---|---|---|
| **PAY1657** | BY4741 MATa *ura3Δ0 leu2Δ0 his3Δ1 met15Δ0 SSA1(1-502)-RFP-clonNAT* | This study |
| **PAY1660** | BY4741 MATa *ura3Δ0 leu2Δ0 his3Δ1 met15Δ0 tif51A-1::kanR SSA1(1-502)-RFP-clonNAT* | This study |
| **PAY1663** | BY4741 MATa *ura3Δ0 leu2Δ0 his3Δ1 met15Δ0 tif51A-3::kanR SSA1(1-502)-RFP-clonNAT* | This study |
| **PAY1692** | BY4741 MATa *ura3Δ0 leu2Δ0 his3Δ1 met15Δ0 SSA1(1-197)-RFP-clonNAT* | This study |
| **PAY1695** | BY4741 MATa *ura3Δ0 leu2Δ0 his3Δ1 met15Δ0 tif51A-1::kanR SSA1(1-197)-RFP-clonNAT* | This study |
| **PAY1698** | BY4741 MATa *ura3Δ0 leu2Δ0 his3Δ1 met15Δ0 tif51A-3::kanR SSA1(1-197)-RFP-clonNAT* | This study |
| **PAY1701** | BY4741 MATa *ura3Δ0 leu2Δ0 his3Δ1 met15Δ0 SSA1(1-83)-RFP-clonNAT* | This study |
| **PAY1704** | BY4741 MATa *ura3Δ0 leu2Δ0 his3Δ1 met15Δ0 tif51A-1::kanR SSA1(1-83)-RFP-clonNAT* | This study |
| **PAY1707** | BY4741 MATa *ura3Δ0 leu2Δ0 his3Δ1 met15Δ0 tif51A-3::kanR SSA1(1-83)-RFP-clonNAT* | This study |
| **PAY1975** | BY4741 MATa *ura3Δ0 leu2Δ0 his3Δ1 met15Δ0 pHSP12-GFP-his3MX6* | This study |
| **PAY1976** | BY4741 MATa *ura3Δ0 leu2Δ0 his3Δ1 met15Δ0 tif51A-1::kanR pHSP12-GFP-his3MX6* | This study |
| **PAY2059** | BY4741 MATa *ura3Δ0 leu2Δ0 his3Δ1 met15Δ0 SSA4-RFP-clonNAT* | This study |
| **PAY2061** | BY4741 MATa *ura3Δ0 leu2Δ0 his3Δ1 met15Δ0 tif51A-1::kanR SSA4-RFP-clonNAT* | This study |
| **PAY2064** | BY4741 MATa *ura3Δ0 leu2Δ0 his3Δ1 met15Δ0 tif51A-3::kanR SSA4-RFP-clonNAT* | This study |
| **PAY2068** | BY4741 MATa *ura3Δ0 leu2Δ0 his3Δ1 met15Δ0 HSP26-GFP-his3MX6* | This study |
| **PAY2071** | BY4741 MATa *ura3Δ0 leu2Δ0 his3Δ1 met15Δ0 tif51A-1::kanR HSP26-GFP-his3MX6* | This study |
| **PAY2086** | BY4741 MATa *ura3Δ0 leu2Δ0 his3Δ1 met15Δ0 HSP12-GFP-his3MX6* | This study |
| **PAY2089** | BY4741 MATa *ura3Δ0 leu2Δ0 his3Δ1 met15Δ0 tif51A-1::kanR HSP12-GFP-his3MX6* | This study |
| **PAY2092** | BY4741 MATa *ura3Δ0 leu2Δ0 his3Δ1 met15Δ0 tif51A-3::kanR HSP12-GFP-his3MX6* | This study |
| **PAY2113** | BY4741 MATa *ura3Δ0 leu2Δ0 his3Δ1 met15Δ0 HXK1-GFP-his3MX6* | This study |
| **PAY2116** | BY4741 MATa *ura3Δ0 leu2Δ0 his3Δ1 met15Δ0 tif51A-1::kanR HXK1-GFP-his3MX6* | This study |
| **PAY2124** | BY4741 MATa *ura3Δ0 leu2Δ0 his3Δ1 met15Δ0 RTG1-GFP-his3MX6* | This study |
| **PAY2127** | BY4741 MATa *ura3Δ0 leu2Δ0 his3Δ1 met15Δ0 tif51A-1::kanR RTG1-GFP-his3MX6* | This study |
| **PAY2133** | BY4741 MATa *ura3Δ0 leu2Δ0 his3Δ1 met15Δ0 CTT1-GFP-his3MX6* | This study |
| **PAY2136** | BY4741 MATa *ura3Δ0 leu2Δ0 his3Δ1 met15Δ0 tif51A-1::kanR CTT1-GFP-his3MX6* | This study |
| **PAY2167** | BY4741 MATa *ura3Δ0 leu2Δ0 his3Δ1 met15Δ0 pSSA1-RTG1-GFP-his3MX6* | This study |
| **PAY2170** | BY4741 MATa *ura3Δ0 leu2Δ0 his3Δ1 met15Δ0 tif51A-1::kanR pSSA1-RTG1-GFP-his3MX6* | This study |
| **PAY2173** | BY4741 MATa *ura3Δ0 leu2Δ0 his3Δ1 met15Δ0 pHSP12-RTG1-GFP-his3MX6* | This study |
| **PAY2176** | BY4741 MATa *ura3Δ0 leu2Δ0 his3Δ1 met15Δ0 tif51A-1::kanR pHSP12-RTG1-GFP-his3MX6* | This study |
| **PAY2203** | BY4741 MATa *ura3Δ0 leu2Δ0 his3Δ1 met15Δ0 DED1-GFP-his3MX6* | This study |
| **PAY2206** | BY4741 MATa *ura3Δ0 leu2Δ0 his3Δ1 met15Δ0 tif51A-1::kanR DED1-GFP-his3MX6* | This study |
| **PAY2209** | BY4741 MATa *ura3Δ0 leu2Δ0 his3Δ1 met15Δ0 tif51A-3::kanR DED1-GFP-his3MX6* | This study |
| **PAY2287** | BY4741 MATa *ura3Δ0 leu2Δ0 his3Δ1 met15Δ0 SSA1(54% opt)-GFP-his3MX6* | This study |
| **PAY2290** | BY4741 MATa *ura3Δ0 leu2Δ0 his3Δ1 met15Δ0 tif51A-1::kanR SSA1(54% opt)-GFP-his3MX6* | This study |
| **PAY2309** | BY4741 MATa *ura3Δ0 leu2Δ0 his3Δ1 met15Δ0 HSP12(45% opt)-GFP-his3MX6* | This study |
| **PAY2312** | BY4741 MATa *ura3Δ0 leu2Δ0 his3Δ1 met15Δ0 tif51A-1::kanR HSP12(45% opt)-GFP-his3MX6* | This study |
| **yRP2323** | MATα, *his3Δ1 leu2Δ0 ura3Δ0 dhh1::NEO pat1::HIS3* | (Balagopal & Parker, 2009) |
| **PAY2352** | BY4741 MATa *ura3Δ0 leu2Δ0 his3Δ1 met15Δ0 HSF1-GFP-his3MX6* | This study |
| **PAY2355** | BY4741 MATa *ura3Δ0 leu2Δ0 his3Δ1 met15Δ0 tif51A-1::kanR HSF1-GFP-his3MX6* | This study |
| **PAY2357** | BY4741 MATa *ura3Δ0 leu2Δ0 his3Δ1 met15Δ0 tif51A-3::kanR HSF1-GFP-his3MX6* | This study |

**Appendix Table S2. Plasmids used in this study**

| Name | Plasmid description | Source |
|---|---|---|
| **PA171** | GA2256-6HA-*HIS3MX6* | Dr. Gustav Ammerer |
| **PA201** | pFA6a-3HA-*HIS3MX6* | (Longtine *et al*, 1998) |
| **PA213** | pRS415-*LEU2* | (Mumberg *et al*, 1995) |
| **PA242** | pFA6a-GFP-*HIS3MX6* | (Longtine *et al*, 1998) |
| **PA245** | pRS415-DCP2-RFP-*LEU2* | (Sheth & Parker, 2003) |
| **PA367** | pYM43-Redstar2-clonNAT | Euroscarf (P30255) |
| **PA369** | pRS415-GPD-*SSA1*-FLAG-*LEU2* | Dr. Thomas Nyström |
| **PA418** | pRS316-ΔssCPY*-GFP-*URA3* | (Medicherla *et al*, 2004) |
| **PA420** | pRP1661-*PUB1*-mCherry-*URA3* | (Buchan *et al*, 2008) |
| **PA423** | pDP122-pNH605-4xHSEpr-YFP-*LEU2* | (Zheng *et al*, 2016) |

**Appendix Table S3. Oligonucleotides used in this study**

| Primer | Sequence (5' - 3') |
|---|---|
| **Oligonucleotides used for gene expression detection by RT-qPCR, ChIP, and RIP-qPCR analyses** | |
| ACT1-F | TCGTTCCAATTTACGCTGGTT |
| ACT1-R | CGGCCAAATCGATTCTCAA |
| CTT1-F | AGACCAGACGGCCCTATCTT |
| CTT1-R | GACTACACGCTCCGGAACTC |
| GFP-F | CCTTGTTAATAGAATCGAGT |
| GFP-R | ATTCCATTCTTTTGTTTGTC |
| HSF1-F | CACCAATGCTACAACCAACG |
| HSF1-R | TTGCTGAGATTGTTGGCAAG |
| HSP104-F | AGCAGGCTCGTCAAGGTAAA |
| HSP104-R | TTACCGATACCTGGCTCACC |
| HSP12-F1 | CTTCCAAGGTGTCCACGACT |
| HSP12-R1 | TTGGTTGGGTCTTCTTCACC |
| HSP150-F | TCTGAGCCTTGGTCCACTTT |
| HSP150-R | GAGATTGGTTGAACGGCAAT |
| HSP26-F | AAGGCGGCTTAAGAGGCTAC |
| HSP26-R | ATCATAAAGAGCGCCAGCAT |
| HSP42-F | GACAGCGGCACTACAAGACA |
| HSP42-R | TGCCTTCATATGGTGGAACA |
| HXK1-F | GTACCCAGCTTCCCAAAACA |
| HXK1-R | GGCCTTCGACATTTGGAATA |
| RFP-F | ACGGTGTCTTGAAGGGTGAA |
| RFP-R | TGGACTTGCTTCTTGGCCTT |
| RPL28-F | AAGCAACAAGCTCATTTCTGGAA |
| RPL28-R | TGGGATCAATGTCCACAATTTG |
| RPL30-F | ACTTCCAAGGTGGTAACAACGAA |
| RPL30-R | AAATAGAGACAACACCGACTCTGAATAA |
| SSA1/2-F6 | TGCCAACCCAATCATGTCTAA |
| SSA1-R3 | CGTTATTATCAATTGCCGCACC |
| SSA1(5'UTR)-F | CGTTCTTGTGGATTGTTG |
| SSA1(5'UTR)-R | CTTTTTGGGTTCTTTCTCTC |
| SSA2(3'UTR)-R | CGGATATTTTACAGGGCGAT |
| SSA3-F | ATTGGAAACGAATGGAAGG |
| SSA3-R | GAAAAATGAGCAACACACGA |
| SSA4-F | GGCTGTCAGAATATGGGGCCGTAGTA |
| SSA4-R | CACCCCGAAGCTGCTTTCACAATAC |
| SSA4(5'UTR)-F | TATGTACTCTCCCACCATT |
| SSA4(5'UTR)-R | TTACTGTCGTCAAACTAAGG |
| TIF51A-F | CGATTCTACTTCTGTAGCCA |
| TIF51B-F | CTACACTTTAGTTCCCTTAC |
| TIF51A/B-R | TCGACAATCTTACATGGTCT |
| **Oligonucleotides used for PCR-mediated gene tagging** | |
| CTT1-GFP-F | GACCTTGAAGGTTATGCCAAGACTTGGTCCATTGCAAGTGCCAATCGGATCCCCGGGTTAATTAA |
| CTT1-GFP-R | TTATGGAGATATAATTACGAATAATTATGAATAAATAGTGCTGCCGAATTCGAGCTCGTTTAAAC |
| DED1-GFP-F | TCTGGCTGGGGTAACAGCGGTGGTTCAAACAACTCTTCTTGGTGGCGGATCCCCGGGTTAATTAA |
| DED1-GFP-R | TGCTAGAGCAGAAAACGAAGAATCCTCACCCTAGTTTGTCTGAAAGAATTCGAGCTCGTTTAAAC |

| | |
|---|---|
| HSF1-GFP-F | CCGACAGAGTACAACGATCACCGCCTGCCCAAACGAGCTAAGAAACGGATCCCCGGGTTAATTAA |
| HSF1-GFP-R | ATATTAAATGATTATATAGCGTATTTAATGACCTTGCCCTGTGTAGAATTCGAGCTCGTTTAAAC |
| HSP104-RFP-F | GATGACGATAATGAGGACAGTATGGAAATTGATGATGACCTAGATCGTACGCTGCAGGTCGAC |
| HSP104-RFP-R | TACTGCTTCTTGTTCGAAAGTTTTTAAAAATCACACTATATTAAATCAATCGATGAATTCGAGCTCG |
| HSP12-6HA-F | TATGTTTCCGGTCGTGTCCACGGTGAAGAAGACCCAACCAAGAAGTCCGGTTCTGCTGCTAG |
| HSP12-6HA-R | CATCATAAAGAAAAAACCATGTAACTACAAAGAGTTCCGAAAGATCCTCGAGGCCAGAAGAC |
| HSP12-GFP-F2 | TATGTTTCCGGTCGTGTCCACGGTGAAGAAGACCCAACCAAGAAGCGGATCCCCGGGTTAATTAA |
| HSP12-GFP-R2 | CATCATAAAGAAAAAACCATGTAACTACAAAGAGTTCCGAAAGATGAATTCGAGCTCGTTTAAAC |
| HSP150-3HA-F2 | CAATGTACTCCAGTCCACTTGGAAGCTATCGATTTGATAGACTGTCGGATCCCCGGGTTAATTAA |
| HSP150-3HA-R1 | AAATGAGAAAAAGTCATCAGGATAAAAGAACTAATAGTTTTCTGCGAATTCGAGCTCGTTTAAAC |
| HSP26-GFP/3HA-F2 | CACGTCAAGAAGATTGAGGTTTCTTCTCAAGAATCGTGGGGTAACCGGATCCCCGGGTTAATTAA |
| HSP26-GFP/3HA-R1 | ACAATGGTCCTCGCGAGAGGGACAACACTATAGAGCCAGGTCACTGAATTCGAGCTCGTTTAAAC |
| HXK1-GFP-F2 | GAAAAAAGAATTGCCGAAGGTAAGTCTCTTGGTATCATTGGCGCTCGGATCCCCGGGTTAATTAA |
| HXK1-GFP-R1 | TTAAGGGAGGGAAAAACACATTTATATTTCATTACATTTTTTTCAGAATTCGAGCTCGTTTAAAC |
| pHSP12-GFP-F | TCTCAAACAAACAACTCAAAACAAAAAAAACTAAATACAACAATGCGGATCCCCGGGTTAATTAA |
| pSSA1-GFP-F2 | AAGTATTACAAGAAACAAAAATTCAAGTAAATAACAGATAATATGCGGATCCCCGGGTTAATTAA |
| pSSA1-RFP-F1 | AAGTATTACAAGAAACAAAAATTCAAGTAAATAACAGATAATATGTCGTACGCTGCAGGTCGAC |
| RTG1-GFP-F2 | AGCTCCTTCGAGTACGGAGGGTATGGTGAGTACGGTAATGGTAGCCGGATCCCCGGGTTAATTAA |
| RTG1-GFP-R1 | ACGAGGGTTATCACAACATAGCAATAGTGAGAGTCAGAAGTACTTGAATTCGAGCTCGTTTAAAC |
| SSA1(1-502)-RFP-F5 | GTCGAAAAGGGTACTGGTAAGTCTAACAAGATCACTATTACCAACCGTACGCTGCAGGTCGAC |
| SSA1(1-197)-RFP-F7 | TTGGACAAGAAGGGTAAGGAAGAACACGTCTTGATTTTCGACTTGCGTACGCTGCAGGTCGAC |
| SSA1(1-83)-RFP-F6 | GCTAAGCGTTTGATCGGTAGAAACTTCAACGACCCAGAAGTGCAGCGTACGCTGCAGGTCGAC |
| SSA1-GFP-F3 | CCTCCAGCTCCAGAGGCTGAAGGTCCAACCGTTGAAGAAGTTGATCGGATCCCCGGGTTAATTAA |
| SSA1-GFP-R2 | CATTAAAAGACATTTTCGTTATTATCAATTGCCGCACCAATTGGCGAATTCGAGCTCGTTTAAAC |
| SSA1-RFP-F4 | CCTCCAGCTCCAGAGGCTGAAGGTCCAACCGTTGAAGAAGTTGATCGTACGCTGCAGGTCGAC |
| SSA1-RFP-R | CATTAAAAGACATTTTCGTTATTATCAATTGCCGCACCAATTGGCTCAATCGATGAATTCGAGCTCG |
| SSA2-RFP-F | TCCTCCAGCTCCAGAAGCTGAAGGTCCAACTGTCGAAGAAGTTGATCGTACGCTGCAGGTCGAC |
| SSA2-RFP-R | CAAAAGTAAAACTTTTCGGATATTTTACAGGGCGATCGCTAAGCTTCAATCGATGAATTCGAGCTCG |
| SSA3-6HA-F | ACGGGAGGTGGAGAAGATACAGGTCCAACAGTGGAAGAGGTTGATTCCGGTTCTGCTGCTAG |
| SSA3-6HA-R | GGTTAAACATAAAAAGTAGCTAAATAGAACACTATAGAAGAATAACCTCGAGGCCAGAACAG |
| SSA4-RFP-F | CCCACTGGAGCACCAGACAACGGCCCAACGGTTGAAGAGGTTGATCGTACGCTGCAGGTCGAC |
| SSA4-RFP-R | ACTAAGAAATTCGATGCTGCTACTTCATCGCATCTTTGTATTTATTCAATCGATGAATTCGACCTCG |
| **Oligonucleotides used for additional genomic fusions** | |
| pHSP12-RTG1-F | TAATCTCAAACAAACAACTCAAAACAAAAAAAACTAAATACAACAATGAGCAGCATTCCAGCTGG |
| pSSA1-RTG1-F | ATCAAGTATTACAAGAAACAAAAATTCAAGTAAATAACAGATAATATGAGCAGCATTCCAGCTGG |
| **Oligonucleotides for the generation of deoptimized variants** | |
| HSP12(5'UTR)-F | CAAAAAAAACTAAATACAACAATGTC |
| PA242-linker-GFP_F | CGGATCCCCGGGTTAATTAACAGT |
| PA242-linker-GFP-R | AGTGAAAAGTTCTTCTCCTT |
| SSA1(5'UTR)-F | ATCAAGTATTACAAGAAACA |

**Appendix Table S4. Antibodies used in this study**

| Primary Antibody | Dilution | Source | Secondary Antibody | Dilution | Source |
|---|---|---|---|---|---|
| **Protein detection by Western blotting** | | | | | |
| eIF5A | 1:500 | Abcam (Ab32407) | α-rabbit | 1:10000 | Promega (W4011) |
| FLAG | 1:5000 | Sigma-Aldrich (F1804) | α-mouse | 1:10000 | Promega (W4021) |
| GFP | 1:5000 | Roche (11814460001) | α-mouse | 1:10000 | Promega (W4021) |
| G6pdh | 1:15000 | Roche (10127671001) | α-rabbit | 1:10000 | Promega (W4011) |
| HA | 1:5000 | Roche (12013819001) | - | - | - |
| RFP | 1:1000 | Thermo Fisher Scientific (A00682-40) | α-rabbit | 1:10000 | Promega (W4011) |
| Hog1-P | 1:1000 | Cell Signaling Technology (4511) | α-rabbit | 1:10000 | Promega (W4011) |
| eIF2α-total | 1:1000 | Santa Cruz Biotechnology (sc-11386) | α-rabbit | 1:10000 | Promega (W4011) |
| eIF2α-P (Ser 52) | 1:1000 | Santa Cruz Biotechnology (sc-101670) | α-rabbit | 1:10000 | Promega (W4011) |
| **DNA inmunoprecipitation (ChIP)** | | | | | |
| GFP | 10 µL | Roche (11814460001) | Dynabeads Pan Mouse IgG | | Invitrogen (11041) |
| **RNA inmunoprecipitation (RIP)** | | | | | |
| eIF5A | 8 µL | Selleck Chemicals (F2900) | Dynabeads anti-rabbit IgG | | Invitrogen (11204D) |

#### Appendix Supplementary Methods

##### Yeast strains, plasmids and growth conditions

All *Saccharomyces cerevisiae* strains and plasmids used in this study are listed in Supplementary Tables S1 and S2, respectively (see Appendix). Cells were cultured in either liquid YPD medium (2% glucose, 2% bacteriological peptone, 1% yeast extract), YP medium (2% bacteriological peptone, 1% yeast extract), or synthetic complete (SC) medium. The SC medium consisted of 0.7% yeast nitrogen base (YNB, Fisher Scientific) without amino acids, the appropriate Drop-Out mixture (Kaiser, Formedium), and 2% glucose (or 2% galactose for experiments requiring galactose induction, referred to as SGal) as the carbon source. When required, SC medium lacking specific amino acids was used for the selection of auxotrophic markers. Solid media were prepared by adding 2% agar.

For all experimental assays, cells were grown exponentially for at least four generations at 25ºC or 30ºC until reaching the desired Optical Density 600 nm ($OD_{600}$). Temperature-sensitive strains were initially cultured at the permissive temperature of 25ºC and transferred to restrictive temperatures (37ºC, 39ºC, 41ºC, or 46ºC) for the specific durations detailed in each experiment to induce the depletion of eIF5A (Li et al., 2014). For some experiments described in the text, the media were supplemented with 4 mM spermidine (Sigma-Aldrich; S0266).

For survival assays using the *tif51AΔ tif51BΔ* strain expressing the murine eIF5A isoform under the control of a galactose-inducible promoter (*pGAL-meIF5A*), cells were grown in SGal-Trp medium at 25ºC until reaching the exponential phase. Subsequently, to deplete eIF5A levels, the cultures were shifted to a glucose-containing medium (SC-Trp) for 6 h at 25ºC prior to a heat shock at 46ºC. Control cells were continuously maintained in galactose medium (SGal-Trp).

To repress *TIF51A* expression in the *tetO$_7$-TIF51A* strain, doxycycline was added. For experiments performed in liquid cultures, cells were grown in YPD or SC medium at 25ºC in the dark, either with or without doxycycline (15 µg/mL) until reaching the early exponential phase. The following day, fresh doxycycline was added, and cells were shifted to 37ºC for 4 h, followed by a 30-min incubation at 46ºC. For phenotypic growth assays on solid media, doxycycline (0.1 µg/mL) was added to the autoclaved and cooled agar medium.

**Strain engineering and plasmid transformation**

For protein quantification and visualization under the microscope, a PCR-based homologous recombination approach was employed for genomic tagging of full-length open reading frames (ORFs) (*SSA1*, *SSA2*, *SSA3*, *SSA4*, *HSP150*, *HSP104*, *HSP26*, *HSP12*, *CTT1*, *HXK1*, *RTG1*, and *HSF1*) with HA, GFP or RFP tags. This same strategy was also used for the complete replacement of target ORFs with other genes or specific reporter sequences. The plasmids pFA6a-3HA-HIS3MX, pFA6a-GFP-HIS3MX (Longtine *et al*, 1998), and pYM43-Redstar2-clonNAT (Euroscarf; P30255) were used as templates for the PCR reaction. Amplifications were performed using the specific primers listed in Supplementary Table S3 (see Appendix), which contained 45 bp homologous to the target genomic region and 20 bp complementary to the vector sequence.

All genetic modifications and plasmid introductions were performed following the standard lithium acetate transformation method (Gietz *et al*, 1992). Briefly, approximately $2 \times 10^9$ cells were harvested by centrifugation at 3,000 × g for 4 min, washed, and resuspended in 1 mL of sterile distilled water. Aliquots of 100 µL were centrifuged again, and the cell pellets were resuspended in a transformation master mix containing 140 µL of 1 M LiAc, 140 µL of 10X TE buffer, 960 µL of 50% PEG 3350, 160 µL of water, and 20 µL of salmon sperm DNA (10 mg/mL; Sigma-Aldrich; D1626), along with approximately 200 ng of the DNA of interest (PCR cassette or plasmid). Mixtures were homogenized by vortexing, incubated at 25ºC for 30 min, and subjected to a heat shock at 42ºC for 20 min. Transformants were selected on appropriate solid media allowing for auxotrophic or antibiotic resistance selection (supplemented with 100 µg/mL nourseothricin/clonNAT [GoldBio; N-500-1]) and incubated at 25ºC or 30ºC for 3 days. All genomic integrations were verified by conventional PCR, western blot analysis, and/or Sanger sequencing (Macrogen Inc.), as required.

To construct yeast strains with codon-deoptimized chaperones, synthetic DNA sequences containing the deoptimized ORFs of *SSA1* and *HSP12* were designed *in silico* according to the codon optimality classification described by (Presnyak *et al*, 2015). These fragments were synthesized and provided by Integrated DNA Technologies (IDT). To integrate these sequences into the yeast genome, the deoptimized fragments, including their respective endogenous 5' UTRs, were fused to a GFP-tagging cassette utilizing an overlap extension PCR strategy (Ho *et al*, 1989). The resulting fusion cassettes, flanked by sequence homology to their native 5' and 3' UTRs, were ultimately transformed into the corresponding strains as described above.

**Growth and viability analyses**

To evaluate cellular growth on solid media, cells were grown to the early exponential phase and cultures were normalized to an identical starting cell density across all strains. Serial 10-fold dilutions were prepared and 5 μL of each dilution was spotted onto the corresponding agar plates. Plates were then incubated at the indicated temperatures for 2-3 days before being imaged.

To determine cell viability over time under the specified experimental conditions, colony-forming unit (CFU) assays were performed. Briefly, early exponential phase cultures were subjected to the indicated treatments. At specified time points, aliquots were collected, serially diluted, and plated onto solid media. After 2-3 days of incubation at 25ºC, visible colonies were counted to calculate cell viability, expressed as the percentage of survival relative to the initial time point (t = 0).

**Polyribosome profile analysis**

For polysome fractionation, cells were grown at 25ºC in SC-Complete medium to the early exponential phase and subjected to the indicated treatments. Cell extractions and polysome gradients were performed as described by (Garre *et al*, 2013; and Ramos-Alonso *et al*, 2018). Briefly, a culture volume containing approximately 2 × $10^9$ cells was incubated for 5 min at room temperature in the presence of 100 μg/mL CHX. Cells were harvested by centrifugation at 3,000 × g for 3 min at 4ºC and washed twice with 2 mL of lysis buffer (20 mM Tris-HCl pH 8.0, 140 mM KCl, 5 mM $MgCl_2$, 0.5 mM DTT, 100 μg/mL CHX, 500 μg/mL heparin, 1% Triton X-100). The cell pellet was then resuspended in 700 μL of lysis buffer and transferred to a 1.5 mL screw-cap tube containing 500 μL of 0.5 mm sterile glass beads. Mechanical disruption was performed by vortexing for eight 30 seconds cycles, with 30 seconds incubations on ice between each cycle. Lysates were cleared by an initial centrifugation at 2,300 × g for 5 min at 4ºC. The recovered supernatant was subjected to a second centrifugation step at 6,000 × g for 5 min at 4ºC to remove residual debris. RNA concentration in the cleared lysate was estimated using a NanoDrop spectrophotometer (Thermo Fisher Scientific). Finally, glycerol was added to the extracts to a final concentration of 5% before flash-freezing and storage at -80ºC. For translational run-off experiments, the same protocol was followed, except that CHX was completely omitted from both the harvesting step and the lysis buffer and after the heat shock, the samples were incubated at room temperature for 10 minutes.

To perform polysome profiling, 8.5 $A_{260nm}$ units of each lysate were loaded onto 10–50% sucrose gradients and separated by ultracentrifugation at 210,000 × g maximum RCF (RCFmax) for 2 h 40 min at 4ºC using a Beckman SW41Ti rotor. Gradients were subsequently fractionated by pumping 60% sucrose from the bottom of the tube, continuously monitoring the absorbance at 240 nm to record the polysome profiles using a density gradient fractionation system (Teledyne ISCO, Lincoln, NE, USA). A total of 22 fractions of 0.5 mL were collected. Total RNA from each pooled fraction was extracted using the SpeedTools Total RNA Extraction Kit (Biotools; 21211), incorporating an rDNase treatment after the elution step according to the manufacturer's instructions. Target mRNAs were analyzed by RT-qPCR using specific primers listed in Supplementary Table S3 (see Appendix). The relative abundance of each mRNA was expressed as a percentage of the total transcript distributed across the fractions. All polysome profiling experiments were performed at least in biological triplicates and a representative profile is shown.

**RT-qPCR analysis**

To analyze mRNA levels, total RNA was isolated from yeast cells following the phenol-chloroform-based extraction method (Barba-Aliaga & Alepuz, 2022). Approximately 3 × $10^8$ cells were harvested and flash-frozen. For cell lysis, the pellet was resuspended in 500 µL of LETS buffer (0.1 M LiCl, 10 mM EDTA pH 8.0, 0.2% SDS, 10 mM Tris-HCl pH 7.4) and transferred to a screw-cap tube containing 500 µL of 0.5 mm sterile glass beads and 500 µL of acid-phenol:chloroform (5:1, pH 4.5). Mechanical cell lysis was then performed using a Precellys 24 tissue homogenizer (Bertin Technologies) for three cycles of 30 s at 6,500 rpm. Following homogenization, samples were centrifuged at 15,800 × g for 5 min at 4ºC and the supernatant was carefully transferred to a fresh tube with 500 µL of acid-phenol:chloroform (5:1, pH 4.5) and then to a tube with 500 µL of chloroform:isoamyl alcohol (24:1) to remove residual phenol and organic solvents. Following a final centrifugation step, RNA in the aqueous phase was precipitated by adding 0.1 volumes of 5 M LiCl and 2.5 volumes of ice-cold 96% ethanol, followed by overnight incubation at -20ºC. The resulting RNA pellets were washed with cold 70% ethanol and dissolved in RNase-free Milli-Q water for later evaluation of RNA concentration using a NanoDrop 2000 spectrophotometer (Thermo Fisher Scientific).

The reverse transcription and quantitative PCR (qPCR) reactions were performed as detailed in (Garre *et al*, 2013). Briefly, 2.5 µg of the total DNAse-I (Roche; 4716728001) treated RNA was retrotranscribed using an oligo $(dT)_{18}$ with Maxima Reverse

Transcriptase (Thermo Fisher Scientific; EP0743). cDNA was labelled with Supreme qPCR Green Master Mix (NZYtech; MB22403) and the amplification and fluorescence detection were carried out using a CFX96 Touch Real-Time PCR Detection System (Bio-Rad). Quantification cycle (Cq) values were used to determine the relative quantity of cDNA in each sample by interpolation from the corresponding standard calibration curve. Endogenous *ACT1* mRNA levels were used as an internal control for normalization. At least three biological replicates of each sample were analyzed, and the specific primers designed to amplify the target gene fragments are listed in Supplementary Table S3 (see Appendix).

In the case of RIP and polysome profiling experiments, cDNA was synthesized following the same reverse transcription protocol but omitting the prior DNase I treatment.

**Western blotting**

To analyze yeast protein by western blotting, extracts were prepared using an alkaline lysis method as described previously in (Zuzuarregui *et al*, 2015) with minor modifications. Briefly, approximately $3 \times 10^8$ cells were harvested by centrifugation. The resulting pellets were washed and resuspended in 200 µL of 0.2 M NaOH and incubated for 5 min at room temperature. Following centrifugation at 13,300 × g for 1 min, the cell pellets were resuspended in 100 µL of 2X-SDS lysis buffer (24 mM Tris-HCl pH 6.8, 0.8% SDS, 10% glycerol, 5.76 mM β-mercaptoethanol, 0.04% bromophenol blue) and boiled at 95ºC for 5 min. Cell lysates were then centrifuged at 830 × g for 10 min at 4ºC, and the supernatants, containing the soluble protein fractions, were transferred to fresh tubes and stored at -20ºC until further analysis.

For protein extraction from sucrose gradient fractions (obtained during polysome profiling), the 22 collected fractions were pooled into four distinct groups: free RNA (L), 60S+80S (M), light polysomes (P(L)), and heavy polysomes (P(H)), as indicated in the corresponding figures. Protein precipitation was performed from 200 µL of the L and M fractions, and from 400 µL of the P(L) and P(H) fractions. Trichloroacetic acid (TCA) was added to each sample to a final concentration of 10%, and proteins were precipitated overnight at -20ºC. Following centrifugation at 15,800 × g for 10 min at 4ºC, the resulting protein pellets were washed twice with 1 mL of ice-cold 100% acetone to remove residual TCA and sucrose. The washed pellets were then resuspended in SDS lysis buffer (100 µL for the L fraction, and 20 µL for the M, P(L), and P(H) fractions) and stored at -20ºC.

Prior to immunoblotting analysis, samples were denatured at 95ºC for 5 min before loading onto SDS-polyacrylamide gels. An equivalent of approximately 80 µg of total protein from whole-cell extracts was loaded per lane, whereas 10 µL of the resuspended L fraction and 20 µL of the M, P(L), and P(H) fractions were used for the polysomal samples. The acrylamide percentage of the resolving gels ranged from 8% to 15%, depending on the molecular weight of the target proteins. Electrophoresis was performed in 1X SDS running buffer (25 mM Tris pH 8.3, 192 mM glycine, 0.02% SDS) using the Mini-PROTEAN Tetra Cell system (Bio-Rad, Hercules, CA, USA) according to standard procedures. Resolved proteins were transferred onto 0.45 µm nitrocellulose membranes (Cytiva; 10600001) using a Mini Trans-Blot Electrophoretic Transfer Cell (Bio-Rad). The transfer was carried out in cold 1X transfer buffer (25 mM Tris-Base, 192 mM glycine, 0.1% SDS, 20% methanol) at 400 mA for 1 h. Blotting membranes were blocked with 5% (w/v) skimmed milk in 1X TBS-T (137 mM NaCl, 2.7 mM KCl, 10 mM $Na_2HPO_4$, 1.8 mM $KH_2PO_4$ pH 7.5, 0.1% Tween-20) for 1 h at room temperature and incubated overnight at 4ºC with the appropriate primary antibodies: anti-GFP (Roche; 11814460001), anti-RFP (Abcam; ab152123), anti-HA (Roche; 12013819001), anti-eIF5A (Abcam; ab32407), or anti-glucose-6-phosphate dehydrogenase (G6pdh) (Sigma-Aldrich; A9521). Following three 10-min washes with 1X TBS-T, bound antibodies were detected by incubating the membranes with the corresponding horseradish peroxidase (HRP)-conjugated secondary antibodies: anti-rabbit IgG (Promega; W4011) or anti-mouse IgG (Promega; W4021) for 1 h at room temperature. The complete list of primary and secondary antibodies used, along with their working dilutions, is provided in Supplementary Table S4 (see Appendix).

After three final 10-min washes with 1X TBS-T, chemiluminescent signals were detected using the Amersham ECL Prime Western Blotting Detection Reagent (Cytiva; 10308449) according to the manufacturer's instructions. Images were acquired using the ImageQuant LAS 4000 software (Cytiva), and band intensities were quantified using ImageJ software (Schindelin *et al*, 2012). To capture variation across all samples, the signal in each lane was normalized to the mean signal across all lanes in a single blot. Then, the resulting signal was normalized against the corresponding G6pdh signal or total protein (Ponceau staining), as indicated in each specific experiment. At least three biological replicates of each sample were analyzed.

**Protein stability determinations**

To assess protein stability, translation was arrested by supplementing cultures with cycloheximide (CHX; Sigma-Aldrich; C7698) at a final concentration of 100 µg/mL following the indicated treatments. Then samples were harvested at specified time points over a 5-h period post-CHX addition, and the decrease in protein levels was analyzed by western blotting. Protein quantification data were normalized to the initial time point (t = 0, set as 100%). The degradation rate constant ($k_d$) for each protein was determined by fitting the normalized data to an exponential decay model, where $k_d$ was extracted from the exponent of the fitted curve ($y = Ae^{-kd\,t}$).

**Fluorescence microscopy**

For live-cell imaging, cells were grown to the early exponential phase in their respective experimental culture media. Prior to visualization, cultures were centrifuged and washed. Fluorescence images were acquired using a motorized Axio Imager Z1 microscope (Carl Zeiss, Oberkochen, Germany) equipped with a Plan-Apochromat 63x/1.4 oil-immersion objective, a 100-W mercury lamp, and an AxioCam MRm digital camera. Image acquisition was controlled using AxioVision v4.7 software (Carl Zeiss). The following excitation (ex) and emission (em) wavelengths were used to detect the different fluorophores: GFP (ex: 475 nm; em: 509 nm), RFP (ex: 555 nm; em: 583 nm), and mCherry (ex: 587 nm; em: 610 nm). To ensure accurate visual and quantitative comparisons, identical exposure times were maintained across all corresponding samples.

Image processing and quantitative analyses were performed using ImageJ software (Schindelin *et al*, 2012). To analyze the presence of protein aggregates, cells were manually categorized based on the number of visible inclusions per cell (0, 1, 2, or ≥ 3 aggregates). A minimum of 100 cells per strain and condition were evaluated, and data was obtained from at least three independent biological replicates.

**Flow cytometry**

For flow cytometry analysis, cells expressing the *4xHSE-YFP* reporter were grown to the early exponential phase in YPD medium at 25ºC and subjected to heat stress at 39ºC for the indicated times. Following each incubation period, translation was immediately halted by adding CHX to a final concentration of 50 µg/mL. Samples were subsequently incubated at 25ºC for 2 h to allow for the complete structural maturation of the YFP fluorophore, as described in (Zheng *et al*, 2016). Following maturation, cells were harvested by centrifugation at 13,300 × g for 1 min, washed with 1 mL of 1X PBS (137 mM NaCl, 2.7 mM KCl, 10 mM $Na_2HPO_4$, 1.8 mM $KH_2PO_4$), and fixed with 4% paraformaldehyde (PFA) in PBS for 15 min on ice. Fixed cells were centrifuged at 13,300 × g for 1 min, resuspended in 1 mL of PBS, and stored at 4ºC until processing. Fluorescence was quantified using a BD FACSVerse™ flow cytometer, with 50,000 individual events acquired and analyzed per sample. All flow cytometry experiments were performed using at least three independent biological replicates.

**RNA immunoprecipitation (RIP)**

The RNA immunoprecipitation (RIP) protocol was adapted from (Gilbert & Svejstrup, 2006), with minor modifications. Approximately 6 × $10^8$ cells in the exponential growth phase were cross-linked by adding formaldehyde to a final concentration of 1% for 15 min with agitation, maintaining the original culture temperature and conditions. The cross-linking reaction was stopped by adding glycine to a final concentration of 0.14 M for 5 min at room temperature. Then, cells were harvested by centrifugation and washed three times with cold 1X TBS. The cell pellet was resuspended in 500 µL of ice-cold lysis buffer (20 mM Tris-HCl pH 7.5, 140 mM NaCl, 1.8 mM $MgCl_2$, 1 mM DTT, 10 U/mL Ribonuclease Inhibitor [Promega; N2111], Complete Protease Inhibitor Cocktail [Roche; 11836153001], 0.2% Triton X-100), briefly centrifuged (13,800 × g, 10 s, 4ºC), and immediately flash-frozen in liquid nitrogen.

For cellular disruption, pellets were resuspended in 300 µL of ice-cold lysis buffer and transferred to screw-cap tubes containing 300 µL of 0.5 mm sterile glass beads. Cells were mechanically lysed using a Precellys 24 tissue homogenizer (Bertin Technologies) for two cycles of 20 s at 6,500 rpm, with a 1-min incubation on ice between cycles. An additional 150 µL of lysis buffer was added to the tubes, and the glass beads were removed by centrifugation at 380 × g for 1 min at 4ºC. To fragment the nucleic acids, the lysate was sonicated in a cold-water bath using a Bioruptor (Diagenode) with 30 s ON/OFF cycles for a total of 12 min and cleared by centrifugation at 13,300 × g for 15 min at 4ºC. Genomic DNA was subsequently removed by adding a reaction mix containing final concentrations of 10 U/µL DNase I (Roche; 4716728001), 40 U/µL Ribonuclease Inhibitor (Promega; N2111), 25 mM $MgCl_2$, and 10 mM $CaCl_2$ to the samples, followed by incubation at 37ºC for 15 min.

The digestion was stopped by adding EDTA to a final concentration of 20 mM, and the mixture was centrifuged at 13,300 × g for 5 min at 4ºC. A 10 µL aliquot of the resulting supernatant was collected and stored on ice (INPUT fraction). The remaining supernatant (IP fraction) was incubated with 50 µL of Dynabeads™ M-280 Sheep Anti-Rabbit IgG magnetic beads (Invitrogen; 11204D) that had been previously bound to 8 µL of eIF5A antibody (Selleck Chemicals; F2900) overnight at 4ºC. The IP mixture was incubated for a minimum of 3 h at 4ºC with gentle rotation. Following incubation, the bead-associated material was collected using a magnetic rack, and the beads were washed three times with lysis buffer. To elute the immunoprecipitated complexes, 75 µL of elution buffer (50 mM Tris-HCl pH 7.5, 5 mM EDTA, 1% SDS, 10 U/µL Ribonuclease Inhibitor [Promega; N2111]) was added to the beads and incubated for 10 min at 37ºC in a water bath. After centrifugation at 850 × g for 2 min at room temperature, the supernatant was transferred to a fresh tube. This elution step was repeated once, and the supernatants were pooled. Both the IP and INPUT samples were then adjusted to a final volume of 150 µL with elution buffer.

Finally, crosslinking was reversed by adding NaCl and Proteinase K (Roche; RPROTKSOL-RO) to all samples to final concentrations of 200 mM and 0.26 mg/mL, respectively. Samples were first incubated at 42ºC for 15 min to allow the Proteinase K to digest the cross-linked polypeptides, followed by an incubation at 65ºC for 1 h to reverse the formaldehyde crosslinking. Finally, 100 µL of 1X TE buffer was added, and the RNA was purified using a Qiagen RNA Purification Kit (Qiagen; 74104) according to the manufacturer's instructions.

The specific binding of eIF5A to target RNAs was analyzed by RT-qPCR using the specific primers listed in Supplementary Table S3 (see Appendix). Prior to qPCR evaluation, INPUT samples were diluted 1:50, while IP samples were diluted 1:10. The relative binding of the analyzed RNAs to the protein was quantified by normalizing the values obtained in the IP samples against their corresponding INPUT values.

**Chromatin immunoprecipitation (ChIP)**

Chromatin immunoprecipitation (ChIP) assays to monitor the recruitment of Hsf1-GFP to chromatin were performed as previously described (Li *et al*, 2016) with the modifications described in (Barba-Aliaga & Alepuz, 2022). Briefly, exponentially growing yeast cultures were subjected to heat stress by shifting the temperature to 37ºC for the indicated times (0, 10, 20, 30 and 45 min), maintaining control cultures at 25ºC. Cross-linking was induced by adding formaldehyde to a final concentration of 1% and incubating for 15 min under original culture conditions. The reaction was subsequently quenched with 0.14 M glycine by incubating for 5 min at room temperature. Cells were harvested by centrifugation, washed with ice-cold 1X TBS (150 mM NaCl, 20 mM Tris, pH 7.6), and flash frozen.

For cellular disruption, cell pellets were resuspended in lysis buffer (50 mM HEPES pH 7.5, 140 mM NaCl, 1 mM EDTA pH 8.0, 1% Triton X-100, 0.1% sodium deoxycholate, 1 mM phenylmethylsulfonyl fluoride [PMSF], 1 mM benzamidine, Complete Protease Inhibitor Cocktail [Roche; 11836153001]) and mechanically disrupted using 0.5 mm sterile glass beads in a Precellys 24 tissue homogenizer (Bertin Technologies) with 3 cycles of 30 s at 5,000 rpm, with 1-min incubations on ice between cycles. Chromatin was then sheared into 300–600 bp fragments by sonication in a Bioruptor water bath sonicator (Diagenode) with 30-s ON/OFF cycles for 12 min. After clearing the lysates by centrifugation (13,300 × g for 15 min at 4ºC), a 10 µL aliquot of each supernatant was saved as the INPUT fraction. The remaining supernatant (IP fraction) was incubated for 3 h at 4ºC with 35 µL of Dynabeads Pan Mouse IgG (Invitrogen; 11041) magnetic beads pre-conjugated with 10 µL of the anti-GFP antibody (Roche; 11814460001) for detecting Hsf1-GFP binding. Detailed antibody information and working dilutions are provided in Supplementary Table S4 (see Appendix).

Following immunoprecipitation, beads were sequentially washed: twice with lysis buffer, twice with high-salt lysis buffer (lysis buffer supplemented with 300 mM NaCl), twice with wash buffer (10 mM Tris-HCl, 250 mM LiCl, 0.5% NP-40, 0.5% sodium deoxycholate, 1 mM EDTA pH 8), and a final wash with 1X TE buffer (10 mM Tris-HCl pH 7.5, 1 mM EDTA). Protein-chromatin complexes were eluted twice by incubation with 40 µL of elution buffer (50 mM Tris-HCl pH 8, 10 mM EDTA, 1% SDS) for 10 min at 65ºC, and both the IP and INPUT samples were then adjusted to a final volume of 100 µL with elution buffer. Cross-linking was reversed by overnight incubation at 65ºC. Samples were then treated with 20 mg/mL Proteinase K for 1.5 h at 37ºC, and DNA was purified using the NZYGelpure kit (NZYtech; MB01101) according to the manufacturer's instructions.

Hsf1 binding to the *SSA1* gene was quantified by RT-qPCR using the specific primers designed to amplify the heat shock element (HSE) sequences located within the 5' UTR regions (listed in Supplementary Table S3, see Appendix). Prior to qPCR evaluation, IP samples were diluted 1:10, while INPUT samples were diluted 1:100. Target DNA levels were normalized against the corresponding total INPUT DNA values. At least three independent biological replicates were analyzed for each sample.

#### Codon optimality and PPI index determinations

A list of optimal and non-optimal codons in *S. cerevisiae* was procured from (Presnyak *et al*, 2015) and the CDSs of the specific yeast mRNAs were obtained from the Saccharomyces Genome Database (https://www.yeastgenome.org/). The occurrence of each optimal codon was counted and divide it by the total number of codons to calculate the percentages of optimal codons.

The translation dependency of each yeast gene on eIF5A was estimated by calculating the Protein Pause Index (PPI) as described in (Barba-Aliaga *et al*, 2026). Briefly, the number of the top 43 eIF5A-dependent tripeptide motifs present in the encoded protein amino acid sequence was determined. Then, the number of each motif was multiplied by the pause strength value observed when eIF5A is depleted, as revealed by 5PSeq analysis (Pelechano & Alepuz, 2017). Thirdly, the PPI for each gene was obtained as the sum of motifs x strength.

#### Statistical analyses

Quantitative data are expressed as the mean ± standard deviation (SD). All graphs and statistical analyses were generated using GraphPad Prism (version 8.0.2; GraphPad Software, Boston, MA, USA). Statistical significance was determined using a two-tailed Student's *t*-test. Specifically, an unpaired *t*-test was applied when comparing independent groups, whereas a paired *t*-test was utilized when comparing the same strain subjected to different experimental conditions or treatments. A $p$-value $< 0.05$ was considered statistically significant. Unless specified, all experiments were obtained from at least three independent biological replicates performed on different days. For microscopy analyses, at least 100 cells per strain and condition were evaluated, and data were pooled from at least three independent biological replicates.
